# Dating the origin of a viral domestication event in parasitoid wasps attacking Diptera

**DOI:** 10.1101/2024.05.24.595704

**Authors:** Benjamin Guinet, Jonathan Vogel, Ralph S. Peters, Jan Hrcek, Matthew L. Buffington, Julien Varaldi

## Abstract

Over the course of evolution, hymenopteran parasitoids have developed a close relationship with heritable viruses, sometimes even integrating viral genes into their chromosomes. For example, in *Drosophila* parasitoids belonging to the Leptopilina genus, 13 viral genes from the Filamentoviridae family have been integrated and domesticated to deliver immunosuppressive factors to host immune cells, thereby protecting parasitoid offspring from host immune responses. The present study aims to comprehensively characterise this domestication event in terms of the viral genes involved, the wasp diversity affected by this event, and its chronology. Our genomic analysis of 41 Cynipoidea wasps from six subfamilies revealed 18 viral genes that were endogenised during the early radiation of the Eucoilini+Trichoplastini clade around 75 million years ago. Wasps from this highly diverse clade develop not only from *Drosophila* but also from a variety of Schizophora. This event coincides with the radiation of Schizophora, a highly speciose Diptera clade, suggesting that viral domestication facilitated wasp diversification in response to host diversification. Additionally, at least one viral gene was replaced by another Filamentovirus gene in one of the species, highlighting the dynamic nature of viral endogenisation. This study highlights the impact of viral domestication on the diversification of parasitoid wasps.

## 1. Introduction

Parasitoid wasps exhibit remarkable species richness, constituting a major component of insect biodiversity [30]. Their biology is characterized by a peculiar reproductive strategy, wherein the parasitoid wasp larvae develop in (endoparasitoid) or on (ectoparasitoid) their host, mostly other insects. Because they usually ultimately kill their hosts, they are major players in the regulation of insect communities. Throughout evolution, endoparasitoid wasps have maintained a special relationship with viruses. This is evident not only from the abundance and diversity of “free-living” heritable viruses that are injected during oviposition into their hosts [46], [17], [85],[20],[86, 36] but also from the abundance of endogenous viral elements found in their genomes. In line with this idea, it was recently found that Hymenoptera with an endoparasitoid lifestyle had a higher propensity to endogenize (i.e. integrate into their chromosomes) and domesticate (i.e. retain by selection) genes acquired from dsDNA viruses, compared with ectoparasitoids or free-living Hymenoptera [37]. Some of these endogenous viral elements (EVEs) have played an essential role in the interaction between these wasps and immunity of their hosts. In some wasps, entire viral machineries have been domesticated, resulting in the production of “virus-like structures” (VLS) in the reproductive apparatus of females. These VLS enable the delivery of either DNA encoding immunosuppressive factors or immunosuppressive proteins to the host’s immune system [10, 88, 63, 16, 22]. When VLS contain DNA, in systems known as polydnaviruses (PDVs), the DNA integrates into the host hemocyte’s DNA, gets expressed [18] and subsequently influences the host’s physiology and behaviour, ultimately favoring the development of wasp offspring [18, 57]. Alternatively, when VLS contain proteins in systems known as virus-like particles (VLPs), the viral machinery facilitates the entry of virulence proteins into host immune cells, thereby suppressing the host immune response [67],[24]. The domestication of viruses has probably contributed to adaptive radiation observed in some highly speciose clades of endoparasitoid wasps, for instance in the microgastroid complex (Ichneumonoidea: Braconidae) following domestication of an ancestral nudivirus [66], [10].

Recently, a new case of viral domestication has been put forward in the *Drosophila* parasitoids belonging to the genus *Leptopilina* (Figitidae)[22]. Since the 1980’s, it was recognized that *Leptopilina* females protect their offspring by producing VLPs in their venom gland, but the evolutionary origin of the genes responsible for their production was unknown [67, 68, 34]. The analysis of their genomes revealed the presence of 13 EVEs that are clustered in the wasp genome, with some level of synteny conservation among species, supporting the hypothesis of a single endogenization event pre-dating the diversification of *Leptopilina* species [22]. However, the wasp diversity affected by this domestication event and its chronology remain unclear.

These EVEs are specifically expressed in the venom gland during the early pupal stage of the wasp when VLPs are synthesized. Furtermore, a viral DNA polymerase (LbFVorf58) most likely amplifies some of the EVEs (10/13) resulting in a concordant peak in DNA copy number and transcripts levels [22]. The intact open reading frames and the very low *dN/dS* values for all 13 EVEs further indicate that these genes are under strong purifying selection since their entry in wasp genomes. Finally, one of these EVEs (LbFVorf85) has been detected as a protein in purified VLPs, providing further evidence for the involvement of these EVEs in VLP production. Once formed in the venom gland, VLPs are injected into the host along with the egg, allowing the delivery of proteins to host immune cells, ultimately protecting the developing parasitoid from the host immune response [67, 64, 64]. The ancestral virus that provided *Leptopilina* species with these genes belongs to a recently proposed new family of dsDNA viruses, i.e. the Filamentoviridae [36]. Filamentoviridae appear to be specialized on hymenopteran parasitoids, as suggested but the fact that all known Filamentovirus only infect endoparasitoid wasps and that endogenous versions of filamentous genes are preferentially and often found in the genomes of Hymenoptera with endoparasitoid lifestyle, such as for instance *Dolichomitus*sp. (Ichneumonidae) [15] and *Platygaster orseoliae* (Platygastridae) [37][36], while they are extremely rare in the genomes of non-parasitoid hymenopterans [36].

So far, all examined species within *Leptopilina* (6 species) tested positive for these Filamentovirus EVEs (FV EVEs). Two species, attacking similar hosts, belonging to the related genus *Ganaspis*, were negative [22]. This suggests that the endogenization event occurred within Figitidae after the split between *Ganaspis* and *Leptopilina* species (approximately around 91.1 mya, [11]) but before the diversification of *Leptopilina* (<40 mya, [11]). However, because *Ganaspis* is quite distantly related to *Leptopilina*, and because several intermediate clades interleaved between those taxa were not tested, the breadth of the Figitidae diversity concerned by this event is currently unknown. In addition, despite the absence of FV-derived genes in *Ganaspis*, it is still possible that the domestication is predating the split between *Ganaspis* and *Leptopilina*, if a subsequent loss in *Ganaspis* did occur. Additionally, since only one Filamentoviridae genome was available at the time that the *Leptopilina* genomes were screened for viral genes, we can expect the discovery of additional genes, now that more FV have been sequenced [36]. In short, our understanding of this major endogenization event remains incomplete both in terms of the diversity of wasps involved and the viral genes involved.

Figitidae are the most speciose family within Cynipoidea superfamily and play an important ecological role in a wide range of environments by controlling the populations of their widely distributed and diversified hosts [14]. Most of them are koinobiont endoparasitoids (meaning they allow the host to continue development while the parasitoid larva grows) attacking larvae of Schizophora flies. These flies account for one-third of fly diversity within Cyclorrhapha, with over 55,000 known species [6] and are encountered in all parts of the world in various habitats encompassing leaf-mines, decaying fruits, dung or carcass [70]. The known diversity of Figitidae is also remarkable knowing the challenge of morphological identification in this clade [53], with more than 1700 species described for Figitidae. Within the Figitidae family, the Eucoilinae subfamily, to which *Leptopilina* belongs, stands out as the most species-rich, with around 1000 described species, constituting the major part of Figitidae diversity [29, 69, 71].

In this paper, we first tested whether the viral domestication documented in *Leptopilina sp.* is restricted to Figitidae attacking *Drosophila* larvae living in decaying organic matter, as *Leptopilina* species do, or is rather shared with other Figitidae with different ecology (i.e. attacking non-*Drosophila* hosts and/or living in different habitats). Second, we asked the question : if a new wasp species turn out to share the same orthologous EVEs, will they also be associated with VLP production?

To address these questions, the presence of FV-like EVEs was first investigated in 26 Figitidae species covering most of the Figitidae diversity using a combination of PCR and whole genome sequencing. The results revealed a single ancestral endogenization event in the common ancestor of all Eucoilini+Trichoplastini, including *Drosophila* and non-*Drosophila* parasitoids, that occurred around 76.4 million years ago during the late Cretaceous period, roughly coinciding with the timing of Schizophora hosts diversification. Furthermore, we bring to light some unpublished results from a 1999 phD thesis showing that the most early diverging species of this virus-bearing clade, do also produce VLPs. Using the whole genomes dataset, we additionally identify new genes (apart from the 13 genes identified by [22]) that derive from the same ancestral endogenization event, thanks to recent advances in the delineation of Filamentovirus diversity [36]. A second independent endogenization event involving a close relative of LbFV was also observed in one species, which likely resulted in the replacement of a gene acquirred from the ancestral event. These results support the critical role played by Filamentovirus core genes in the production of VLPs in endoparasitoids and their diversification.

## 2. Results

### (a) All Eucoilini+Trichoplastini species contain filamentoviridae genes in their chromosomes

In order to investigate the distribution of Filamentoviridae-derived genes among Figitidae wasps (+four additional outgroup species belonging to Liopteridae and Cynipidae), a PCR screening was conducted on 41 specimens representing 20 genera divided into 6 subfamilies, i.e., around 145 mya of wasp evolution [11] (FigureS2,FigureS3). As expected from previous results, the three *Leptopilina* species were positive, while *Ganaspis* was negative. In addition, we found that *Trybliographa* sp. was also positive (FigureS2). Following this preliminary PCR screening, we sequenced and assembled the genomes of *Trybliographa* sp., along with two related wasps belonging to the Trichoplastini: *Rhoptromeris* sp. and *Trichoplasta* sp. that were selected because of their position in between *Leptopilina* sp. and *Trybliographa* sp.. We also included in our analysis the most basal *Leptolamina* sp. of uncertain tribe [12, 84] and the published genome assemblies of *Leptopilina* (for which we utilized the long read assemblies for *L. heterotoma* and *L. boulardi*), as well as the genome assemblies of *Ganaspis* and *Synergus* (Cynipidae: Ceroptresini). The assemblies of the nine species ranged from 354.80 Mb to 935.64 Mb, with a minimum of 93.8% BUSCO genes (Complete and fragmented) (see TableS1) for all assemblies details). Using the predicted proteins from 7 Filamentoviridae viruses genomes as queries [36], we found significant viral hits in 6 out of 11 genomes: *L. boulardi*, *L. heterotoma*, *L. clavipes*, *Trybliographa sp.*, *Rhoptromeris sp.*, and *Trichoplasta sp.* (Figure1-A). On the contrary, no viral hits were detected in the *Synergus* and *Ganaspis* genomes, nor in that of *Leptolamina*. In total, for the six positive-tested genomes, we identified 153 putative EVEs deriving from the integration of 25 Filamentovirus genes (FigureS1). Overall we found 25.6 EVEs/genomes (sd=14.68) with the highest number of EVEs in *L. boulardi* (n=54) and the lowest in *Trichoplasta* (n=15). For detailed filamentous gene alignments, see FiguresS13. All 107 scaffolds containing the candidate EVEs had coverage and GC profiles similar to those of BUSCO scaffolds, providing evidence for their integration within wasp chromosomes (FigureS4). In addition, 55 out of the 107 EVE-containing scaffolds, also harbored at least one eukaryotic gene or transposable element, further supporting their chromosomal integration into wasp chromosomes.

**Figure 1.**
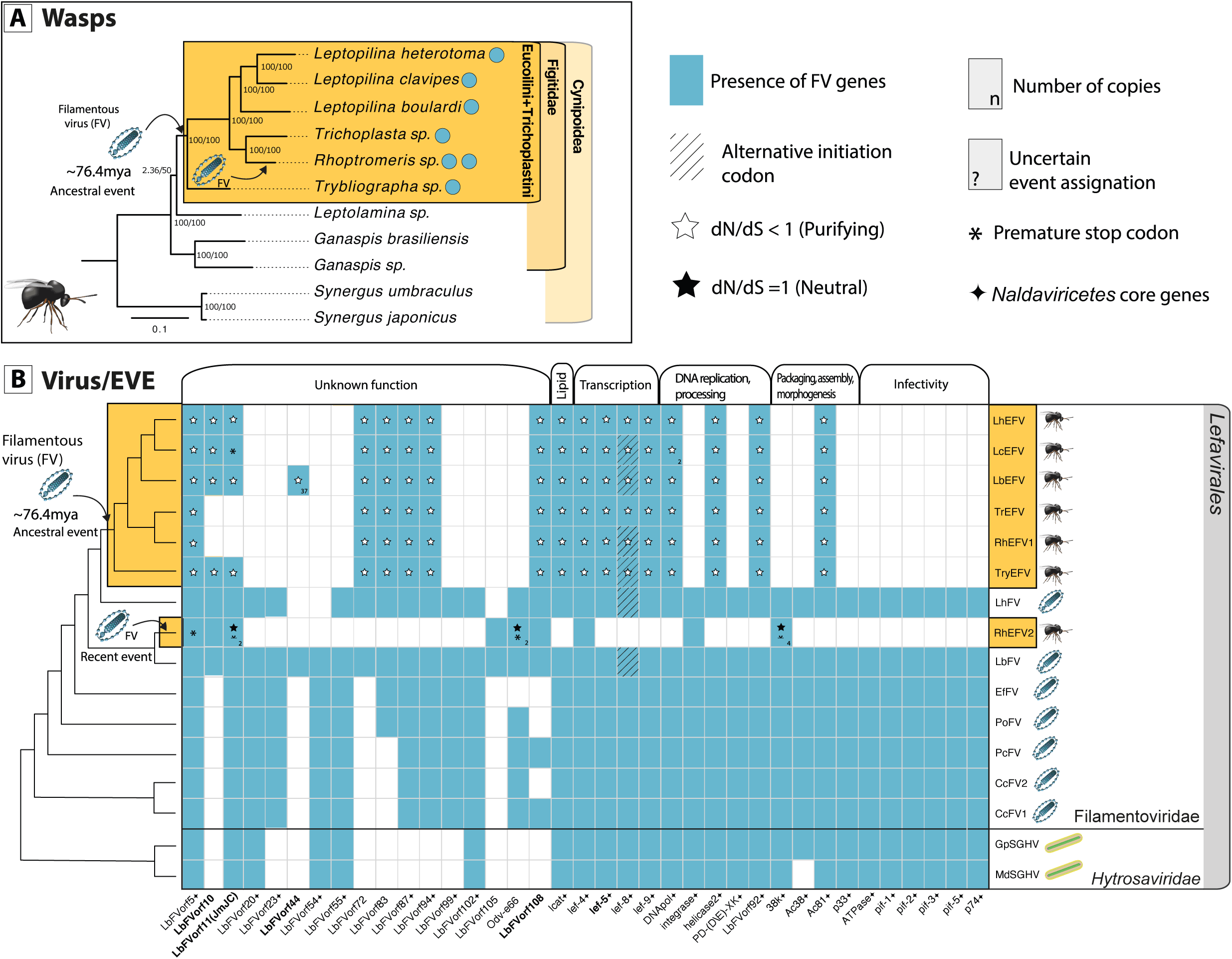
Domestication of Filamentoviridae (FV) in Eucoilini+Trichoplastini wasps. **(A)** The phylogeny of the Cynipoidea has been estimated using 1,000 Busco genes. Confidence scores (aLRT%/ultra-bootstrap support%) are shown at each node. Blue circles indicate the presence of FV EVE in a genome. **(B)** The heatmap represents the distribution of viral ORF in *Hytrosaviridae* virus and Filamentoviridae virus, as well as the distribution of endogenous viral elements (EVEs) from Filamentovirus origin in Eucoilini wasps. All 29 core Filamentovirus genes (with a black diamond next to the gene name) are depicted, whether endogenized or not. The cladogram phylogeny of the virus and EVEs is reported on the left and has been made according to the results from the FigureS6. The rows represent the viral or Eucoilini+Trichoplastini species, and the columns represent the viral ORFs distributed according to their potential functions. When the box is hashed, it indicates that the predicted ORF has an alternative start codon. A white star indicates evidence for purifying selection (*dN/dS*<1) while a black star indicates no evidence for selection (*dN/dS*=1). When multiple paralog EVEs can be found in a species, its number is displayed in the bottom right corner. Asterisk corresponds to the presence of premature stop codons in the EVEs. When multiple copies are present, a semi-asterisk indicates that some copies have a premature stop codon, while at least one copy has a complete ORFs. When an EVE could not be assigned to a particular event, a question mark is displayed in the bottom left corner. Gene names highlighted in bold are the EVES from the ancestral event that are newly described in this study.

### (b) Two independent integration of Filamentovirus occurred in Eucoilini+Trichoplastini

Analyzing each of the 25 individual gene phylogenies revealed three topologies that will be briefly described below (FigureS5, see supplementary material for additional phylogenies).

In **type I** phylogenies, all Eucoilini+Trichoplastini sequences formed a monophyletic clade nested within a Filamentovirus clade, branching with Leptopilina heterotoma Filamentous virus (LhFV). These phylogenies are consistent with a single endogenization event occurring before the divergence of the 6 species and involving a LhFV-like donor.

In **type II** phylogenies, a single sequence from *Rhoptromeris* was nested within a Filamentovirus clade, typically branching with Leptopilina boulardi Filamentous virus (LbFV). These phylogenies suggested a single endogenization event specific to *Rhoptromeris* involving a LbFV-like donor.

In **type III** phylogenies, both type I and type II patterns were observed, indicating an endogenization event involving a LhFV-like donor in the common ancestor of Eucoilini+Trichoplastini species (as in type I) and a second event in the branch leading to *Rhoptromeris* involving a LbFV-like donor (as in type II).

Out of the 25 genes, five were left unclassified because of insufficient signal, while 12 exhibited type I, 3 exhibited type II, and 5 exhibited type III patterns. All filamentous gene phylogenies can be found in FigureS14.

The gene-level phylogenetic analysis thus strongly suggests that two independent endogenization events did occur in this wasp clade: an ancestral event almost basal to the Eucoilini+Trichoplastini and another one more recent and specific to *Rhoptromeris*.

Because EVEs that are physically close to each other (in the same scaffold) are likely to have been acquired during the same endogenization event, we used this closeness as a criterion to group them. 71 out of the 153 EVEs were found to co-occur on the same scaffolds in one or the other genome assemblies (Figure2). As expected, this genomic-location-based grouping systematically grouped together genes with similar phylogenies supporting the hypothesis that co-location indeed indicates common evolutionary history. This way, it was possible to assign two unresolved gene phylogenies to the two events (LbFVorf44 in the ancestral event and LbFVorf105 in the recent *Rhoptromeris* event). Finally, integrating both phylogenetic and co-location information, we were able to assign 150 of the 153 FV-EVEs to the two independent events. In the end, we estimated that 18 and 9 Filamentovirus genes have been jointly acquired following respectively the first and second endogenization events. Note that the set of Filamentovirus genes acquired during these events overlap by five genes (those having “type III” phylogenies). Within each event, the gene phylogenies exhibited congruence, further supporting a shared evolutionary scenario for the different genes assigned to each event. We will respectively name these two independent endogenization events: *ancestral event* and *recent event* in the following sections.

**Figure 2.**
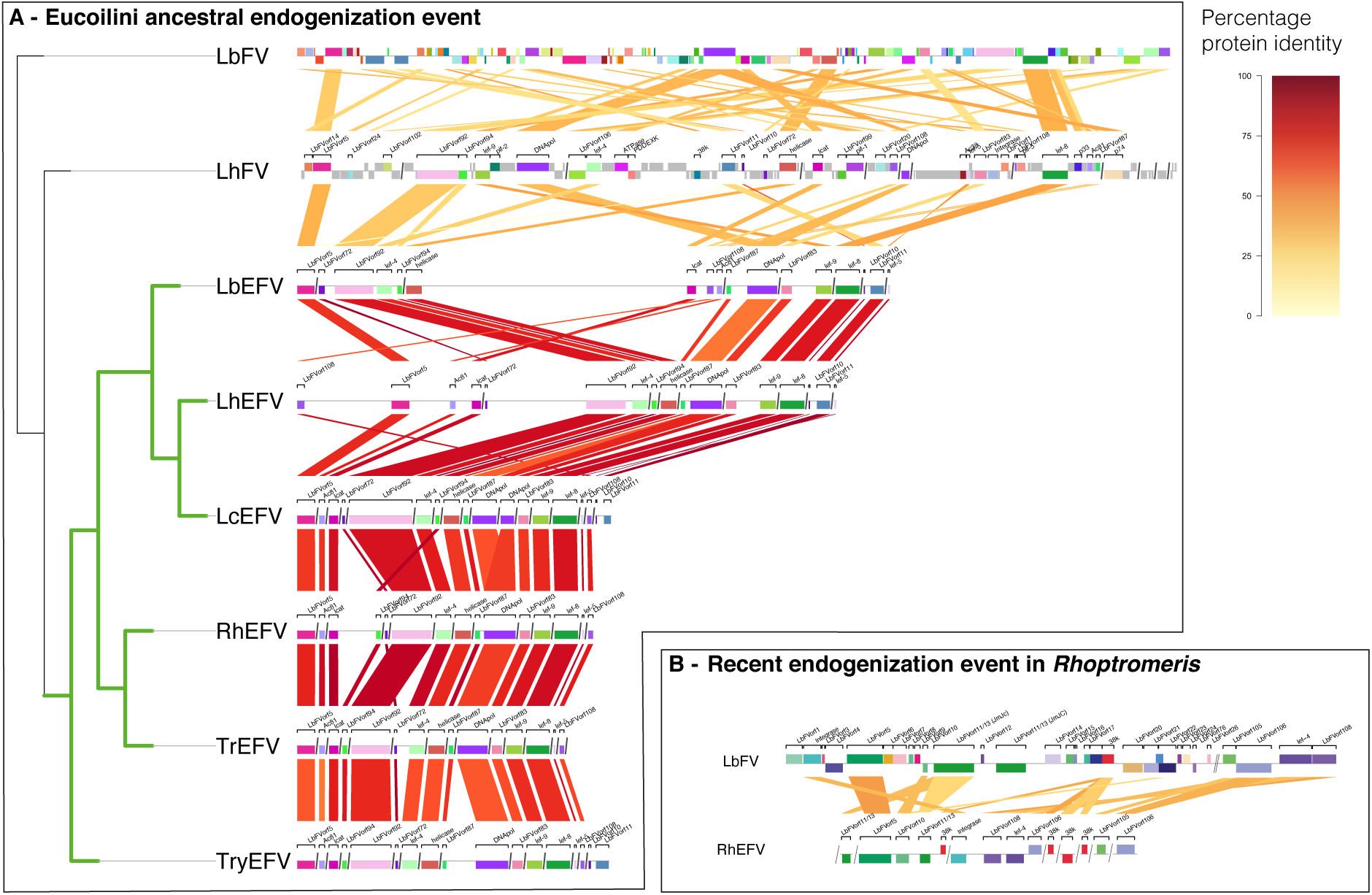
Comparative genomics of wasp scaffolds sharing similarities with filamentous ORFs. LbFV (Leptopilina boulardi Filamentous virus), LhFV (Leptopilina heterotoma Filamentous virus), LbEFV (*Leptopilina boulardi EVEs*), LhEFV (*Leptopilina heterotoma EVEs*), LcEFV (*Leptopilina clavipes EVEs*), RhEFV (*Rhoptromeris EVEs*), TrEFV (*Trichoplasta EVEs*), TryEFV (*Trybliographa EVE*). The phylogenetic tree on the left has been made according to the results from the FigureS6. Green and black branches correspond to Eucoilini+Trichoplastini and Filamentous branches, respectively. The red/yellow color code depicts the percentage of protein identity between homologous sequence pairs (viral or virally-derived loci). Colored boxes identify the virally-derived genes and their orientation (above: sense, below: antisense). Gray connections indicate homology between non virally-derived regions. The figure has been drawn using the genoPlotR package [39]. The scaffolds are ordered from left to right in an arbitrary manner.

**Figure 3.**
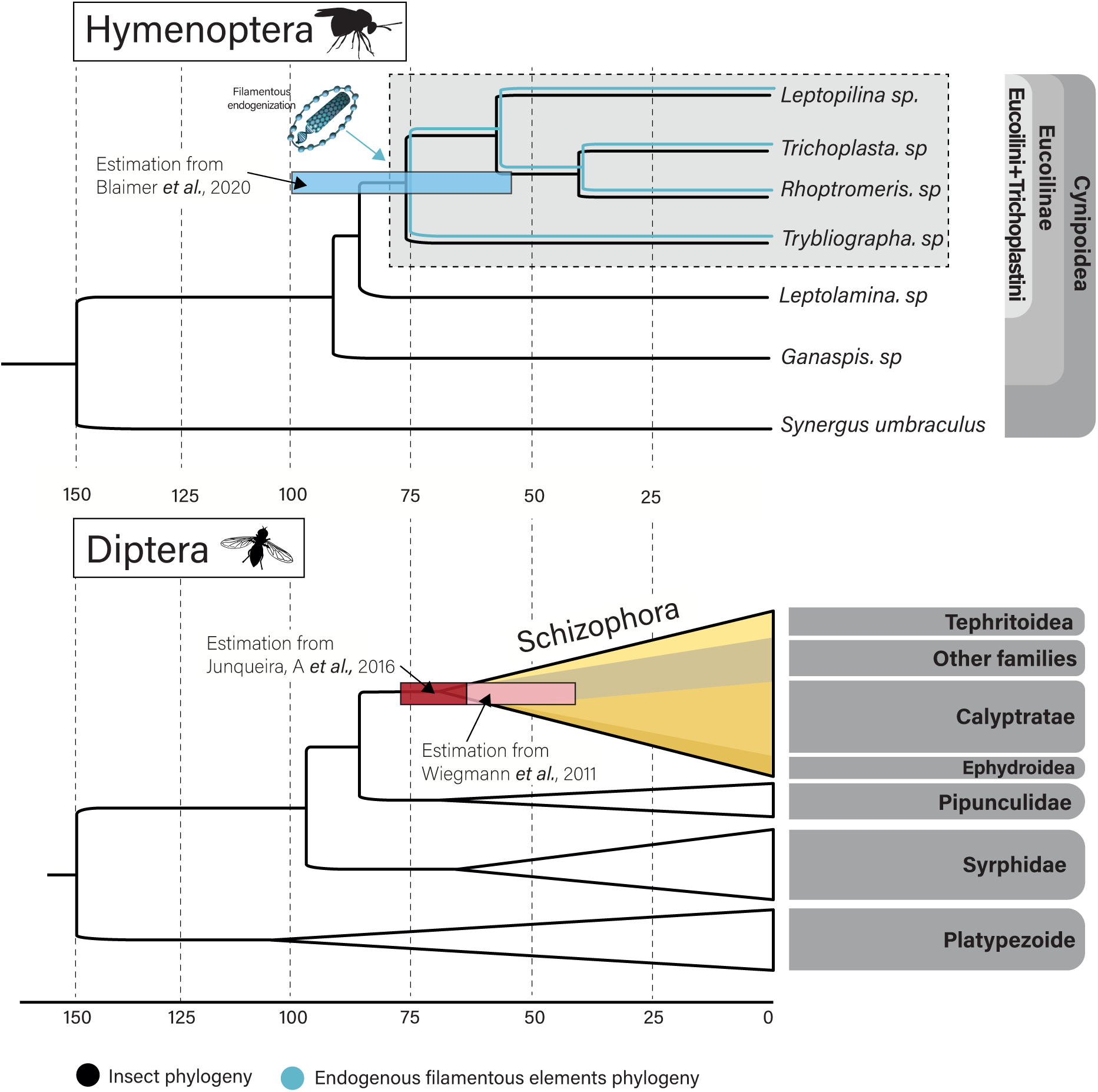
Calibrated phylogenies for Eucoilinae parasitoids, and their dipteran hosts. Black branches correspond to Hymenoptera (up) or *Diptera* (bottom) branches, while blue branches illustrate the presence of the endogenous viral genes. Trees were reproduced using the following sources : [11] for Hymenoptera datation, and [90, 42] for the *Diptera* datation. The corresponding credibility intervals are shown in red tons for the Schizophora node age and in blue for the Eucoilini+Trichoplastini node age.

#### (b1) First ancestral endogenization event within the Eucoilini+Trichoplastini common ancestor

The first ancestral event, in the common ancestor of the 6 Eucoilini species, involved the endogenization of 18 filamentous genes, including the 13 genes previously identified as domesticated in *Leptopilina* species [22]. Concatenating the 15 genes shared by all Eucoilini+Trichoplastini species produced a consistent phylogenetic signal, with Eucoilini+Trichoplastini species forming a highly supported monophyletic clade (bootstrap score =100) nested within Filamentovirus diversity, with LhFV being the closest relative (Figure1-B). As expected from a single endogenization event, the phylogeny of the EVES mirrors the evolutionary history of the species (see Figure1-A and B).

Moreover, the presence of a few conserved gene synteny blocks across Eucoilini+Trichoplastini genomes further supports the conclusion of a single event of endogenization. Notably, gene syntany was particularly evident in the best-assembled genomes of *L. boulardi* and *L. heterotoma*. In both genomes, we observed co-localizations of various genes, such as LbFVorf92/Lef4, DNApol/LbFVorf87, LbFVorf83/Lef9, and LCAT/LbFVorf108/Ac81 (Figure2-A). We also noted a significant gene association between LbFVORF10 and LbFVORF11. This association was not only observed in the free-living viruses LbFV and LhFV but also consistently found in *Leptopilina* species and in the *Trybliographa* genome (Figure2-A).

Overall, the EVEs identified in the ancestral event were enriched for core viral genes (defined at the level of *Naldaviricetes*) with 12 core genes among the 18 EVEs (compared with a putative donor virus such as LhFV with 110 genes and 29 core genes, Fisher test, odd-ratio=5.5, p-value=0.001753). These genes are likely involved in cholesterol metabolism, DNA replication and processing, transcription, packaging and assembly, morphogenesis and unknown functions (Figure1). This enrichment in core genes is in line with the data obtained on other cases of viral domestication in parasitic wasps [10, 88, 63, 16] and suggests that core viral functions are crucial for the production of virus-like particles.

In comparison with our previous study [22], we identified five additional genes acquired during the same ancestral event (Figure1- highlighted in bold), three of which were shared by all Eucoilini+Trichoplastini species. The gene names, their putative functions, and their phylogenies can be found in FigureS14 and the alignments in FiguresS13. These additional genes are briefly described below.

One of these genes, **LbFVorf108**, is shared by all species from this event (except for *Leptolamina*) and localized next to previously documented EVEs in some of the wasp assemblies (Figure2-A). The presence of complete open reading frames (ORF) and indications of strong purifying selection (*dN/dS*=0.2792 (SE=0.03)) in all Eucoilini+Trichoplastini EVEs suggests that LbFVorf108 is functional in Eucoilini+Trichoplastini genomes. However, its function is unknown, as no homologues other than Filamentoviridae are available in public databases.

The **LbFVorf10** and **LbFVorf11 (JmJC)** were consistently found next to each other on the same wasps scaffolds (Figure2-A, B). Both genes are also next to each other in the genomes of their closest relatives LhFV and LbFV, which further suggests they entered into wasp genomes together during each of the two independant events. They have been jointly lost in the related *Trichoplasta sp.* and *Rhoptromeris sp.*. All LbFVorf10 EVEs presented a complete ORF and a *dN/dS* analysis on the EVEs suggested strong purifying selection (mean *dN/dS*=0.1654 (SE=0.033). HHpred analysis revealed homology between LbFVorf10 and a Glycoprotein containing a Zinc finger domain from a Hantavirus protein (Evalue 5.9e-8) (TableS3). Homologs of LbFVorf11 in *L. heterotoma* and *Trybliographa sp.* were under purifying selection but showed either no signs of purifying selection or even premature stop codons in the other *Leptopilina* genome. Interestingly, the phylogeny built on LbFVorf11 homologs suggests that the gene was acquired by Filamentoviridae from eukaryotes. This suggests a two-step integration process from eukaryotes to Filamentoviruses and then to the genomes of wasps (see FigureS14-3).

A ***Lef-5*** homolog was shared by all species (apart from *Leptolamina*). Interestingly, in both LbFV and LhFV, *Lef-5* has an alternative start codon TTG. This feature was conserved in the genomes of the wasps at the exception of *L. heterotoma* and *Trichoplasta sp.* that encode a classic ATG start codon and of *Leptopilina clavipes* that encodes the alternative start codon CTG. The endogenized *Lef-5* showed a complete ORF and *dN/dS* analysis suggests they are under strong purifying selection (mean *dN/dS*=0.1118 (SE=0.03)). This gene might be involved in RNA polymerase initiation transcription factor, as in baculoviruses [82].

***LbFVorf44*** EVE was found only in *Leptopilina boulardi*. It had multiple paralogs (n=36) distributed in 23 scaffolds. One of the LbFVorf44 copies co-occurred in the same scaffold as the *lef-8* EVE. This finding suggests an integration of *LbFVorf44* along with the other genes during the same event, followed by several duplications. A dN/dS analysis of these paralogs suggests that they evolve under purifying selection (mean *dN/dS*=0.3004, (SE= 0.077)). This gene has no known homologs in public databases.

#### (b2) Second recent endogenization event unique to the *Rhoptromeris* genome

The second recent event detected specifically in the *Rhoptromeris* species, involved 9 filamentous genes that ultimately led to the presence of 16 EVEs (this count is higher due to the presence of paralogs). Four of these genes were specific to *Rhoptromeris*, while five were detected in both the first and second events (Figure1). The phylogeny constructed on the concatenated EVEs positions *Rhoptromeris* outside the Eucoilini clade, close to LbFV species, indicating acquisition from an independent event involving a virus related to LbFV, rather than LhFV. Among the EVEs, 7 out of 16 had premature stop codons or incomplete ORFs, suggesting non-functionality for some copies. However, the remaining 9 EVEs may still be functional since they had complete open reading frames. Interestingly, LbFVorf10 probably replaced the homologous

EVE from the first event in *Rhoptromeris* (Figure2-A). Similarly to the first ancestral event, LbFVorf10 and LbFVorf11 were also found next to each other in the *Rhoptromeris* genome, as they are in the genome of their closest relative LbFV, again suggesting that they entered jointly into wasp chromosomes. *Integrase* and LbFVorf106 (*Odv-e66*) were unique to *Rhoptromeris* (Figure2-A). No evidence of selective pressure was observed on the *Odv-e66* EVE (dN/dS not different from 1). The same dN/dS analysis was not feasible on the *Integrase* gene, due to the absence of paralogs.

### (c) All EVEs of the first ancestral endogenization event show signs of domestication

The dN/dS values obtained for the 18 EVEs acquired during the first ancestral event were low (mean=0.18, sd=0.084), and within the range of values observed for the highly conserved BUSCO genes (FigureS7). However, EVEs on average had a slightly higher *dN/dS* values compared to BUSCO genes (T.test two-sided, df=1015, p-value = 6.442e-09), suggesting either a lower stabilizing selection intensity or diversifying selection in some sites. In total, 2,354 codons (out of 7,233) showed evidence of purifying selection (see supplementary excel table): Contrast_FEL_table), while 23 were likely evolving under diversifying selection, possibly indicating some interactions with host proteins (FigureS8).

### (d) The basal *Trybliographa* produce VLPs, as *Leptopilina* species do

In dicussion with colleagues at the University of Rennes (France) who provided us with some of the *Trybliographa rapae* samples used in this study, we were informed that virus-like particles (VLPs) had been discovered during Nabila Kacem-Haddj El Mrabet’s doctoral thesis, which was defended at the same university in 1999 (ref). Since this part of the thesis has never been published in a journal, we have asked the author to reproduce here the main results obtained (see full details of results in supplemental informations)

This PhD project was initially motivated by an applied perspective, since this endoparasitoid wasp attacks the most important root herbivore in *Brassica* crops, i.e., the cabbage root fly *Delia radicum*. By transmission electron microscopy investigations, two types of viral-like structures were detected in *Trybliographa rapae*. One is produced in the venom gland (also called the unpaired gland, FigureS10), similarly to VLPs found in *Leptopilina sp.*, while the other is formed in the paired gland (FigureS11). The particles secreted by the venom gland accumulate in the reservoir, in the same way as the VLPs found in *Leptopilina sp* and were morphologically very similar to the VLPs produced by *Leptopilina sp.* (FigureS10). Both types of particles were found in the ovipositor canal, suggesting that they are both injected with the eggs during oviposition (FigureS10-I and FigureS11-J). While particles produced by the paired gland are abundantly present on the egg surface after oviposition (FigureS11 B-C-D), the ones produced by the venom gland were not found in contact with the parasitoid egg after oviposition, like in *Leptopilina*. As no evidence of encapsulation was observed in several hundred eggs analyzed in the course of this thesis, the researchers attributed an immunosuppressive role to these viral particles. In conclusion, the data accumulated so far show that all wasps concerned by the ancestral endogenization event, including the most basal one, do produce VLPs that share evident morphological and life-history features.

### (e) Timing of the endogenization events

Previous analysis based on fossil calibration at the scale of the Cynipoidea estimated that the common ancestor of *Leptopilina*, *Trichoplasta*, *Rhoptromeris* and *Trybliographa* lived in the Cretaceous period, approximately 76 million years ago (55-100 mya) [11]. Since our analysis shows that the endogenization event (event I) took place in this same common ancestor, we estimate that this major endogenization event occurred at this time period. Based on the same rationale, we estimated that the second endogenization event which only concerns *Rhoptromeris* occurred in the last 40 million years (22-59 mya) since this is the time period of the split between *Rhoptromeris* and *Trichoplasta sp*.

## 3. Discussion

In this study, we analyzed the diversity of cynipoid wasps observed to have a viral domestication event initially reported in *Leptopilina* [22]. From previous work, it was established that (i) VLPs does protect the eggs from encapsulation in *Leptopilina* species [67], and that (ii) in *Leptopilina boulardi*, for which various experimentations and genomic analysis have been conducted, this VLP production was linked to the presence of 13 virally-derived genes [22] . Because all *Leptopilina* species analyzed so far do produce VLPs [22] and because all *Leptopilina* genomes analyzed so far do contain the same conserved LbFV-like genes, it was concluded that all *Leptopilina* species rely on those viral genes to produce the VLPs [22]. Here, after analyzing wasp genomes from 5 Figitidae subfamilies, we propose to extend this reasoning to the whole clade of Eucoilini+Trichoplastini. Indeed, in this clade, all species share a set of 15 highly conserved viral genes (including the 13 previously identified by [22]), and most importantly, the most basal species within this clade, namely *Trybliographa* sp. do produce VLPs (Kacem-Haddj El Mrabet’s PhD research). This finding thus suggests that VLP production is a common feature shared by all species of this clade, not only *Leptopilina* species, supporting the hypothesis that these virally-derived genes are responsible for VLPs production.

Building on the recent sequencing of additional LbFV-related viruses [36], this analysis also provided a comprehensive view of the set of viral genes involved in this event. Five additional viral genes endogenized during the same event were identified, including a *Naldaviricetes* core gene involved in transcription (lef5). The overall picture is that all the core genes involved in transcription, as well as some essential genes involved in DNA replication (including the viral DNA polymerase), have been retained by selection in all wasp genomes of this clade. This conservation strongly suggests that this machinery is essential for wasp fitness and is likely involved in genomic amplification and associated transcription of virally-derived genes, as observed in the venom gland of developing *L. boulardi* females [22]. Additionally, the Ac81 gene which plays a crucial role in baculovirus envelopment [23] is shared by all wasp genomes. This suggests it plays a critical role in the the production of VLP envelope, as also attested by the presence of its proteinic product in mature particles purified from *Leptopilina* species [22]. Both features (transcriptomic and amplification machineries on one side and presence of Ac81) are shared with other VLP systems previously described in the distantly related Ichneumonoidea *Venturia canescens* [63] and *Fopius arisanus* [16]. On the contrary, the picture is completely different on *per os* infectivity factors (PIFs), which are major players for the entry of baculovirus into insect cells. While both *Venturia* and *Fopius* systems genomes do encode most of the known PIFs, none are encoded by the Eucoilini+Trichoplastini genomes. Knowing that VLP from *Leptopilina* permit the delivery of virulence proteins into *Drosophila* immune cells [21], this indicates that other mechanisms not relying on pif proteins have been recruited in Eucoilini+Trichoplastini.

The best understood case of viral domestication in parasitoids involves Braconidae belonging to the microgastroid complex. In this system, the (braco-)virus has been acquired by the ancestral wasps around 103 mya from a nudivirus ancestor [58, 10]. Among the subfamilies that compose this clade, the Microgastrinae is by far the most speciose and has rapidly radiated around 50 mya [58]. It has been hypothesized that this rapid radiation is correlated with a rapid radiation in their lepidopteran hosts which are also very diverse [54]. One hypothesis that was put forward to explain the diversification of this particular subfamily compared to Cheloninae, is that they develop from inside Lepidoptera larvae (while Cheloninae develop from within the eggs) and as such are exposed to the host immune system, since eggs are much less active immunologically than larvae. The idea is that endoparasitoids attacking larvae require specific adaptation for each host species attacked, thus favoring ecological speciation process.

Similar to microgastroids, we found that ancestral filamentovirus domestication also affected a diverse, albeit smaller, clade of wasps (the Eucoilini and Trichoplastini tribes) that diversified around 75 mya. All these species are koinobiont endoparasitoids that attack the larvae of distinct species of Schizophora Diptera in various environments [13]. For instance, *Leptopilina* develops from Drosophilidae, *Rhoptromeris* from Chloropidae flies, *Trybliographa* from Anthomyiidae flies, and *Trichoplasta* develops from Drosophilidae, Muscidae, and Lonchaeidae flies [13]. It is interesting to note that the Schizophora clade, which is the most species-rich group in Cyclorrhapha [6], underwent a rapid radiation just after the Cretaceous-Paleogene (K-Pg) crisis, starting around 65 to 68 million years ago depending on the estimates [42, 90]. The domestication of the virus in Eucoilini+Trichoplastini thus coincides quite well with this period [13]. We can speculate that the acquisition of the virus favored the adaptation to these various hosts, allowing the wasps to cope with the specificities of each host immune system.

The lack of evidence for filamentovirus domestication in *Ganaspis* is remarkable. Members of this genus tend to parasitize the same hosts as *Leptopilina*, often in direct competition with them [52, 1, 60]. In some cases it appears *Ganaspis* can actually push *Leptopilina* from a given niche space (Spotted Wing *Drosophila* working group, unpublished data). As such, we predict another form of host immune system manipulation is at play within *Ganaspis*, and certainly worthy of additional study.

Finally, our analysis also provided evidence for a second event of endogenization involving a closely related donor virus, also belonging to the newly proposed Filamentoviridae family [36]. While we estimate that the major endogenisation event occurred approximately 75 million years ago, soon after the separation of the Eucoilini+Trichoplastini lineage, the more recent endogenisation took place in the last 40 million years, within the branch leading to the genus *Rhoptromeris*. It is unclear whether this second event provided selective advantage to the wasps, since dN/dS analysis was only possible for a few genes having a few paralogs and did not provide evidence for purifying selection. However, it is still possible that some genes acquired during the ancestral endogenization event that were subsequently lost in this clade were replaced by genes from this second endogenization event.

In conclusion, this study shows that the newly proposed viral family Filamentoviridae [36], which is involved in both events, has a long and intimate history of association with endoparasitoid wasps. More generally, this work combined with previous literature [10, 63, 88, 16, 37] show that DNA viruses associated with parasitoid wasps had a strong impact on the evolution and diversification of the parasitoid lifestyle in very distant wasp clades.

## 4. Methods & Materials

### 41 *Cynipoidea* specimens extractions

All extractions were performed with the NucleoSpin Tissue Macherey kit. 31/41 Cynipoidea specimens were extracted with the whole body, while 10/41 were extracted with the metasoma in order to keep the upper body for morphological identification (when only one individual was available). Consequently, we adapted the volumes according to the selected tissues, in the following we call the volumes for the full bodies “c”, and the metasoma “a”. The bodies or metasoma were manually crushed with a piston in a 1.5Ml eppendorf tube with proteinase K (c:180µL/a:60µL) and T1 buffer (c:25µL/a:8µL) and then incubated at 56 °C for 3h. The lysates were then mixed and buffer B3 (c:200µL/a:70µL) was added and incubated at 70 °C for 10 min. The solution was then bound to a silica column, washed with buffer BW (a:500µL/c:250µL) and buffer B5 (a:600µL/c:300µL) and finally eluted in 20µL of TE.

### PCR amplification of ORF96

Based on the sequences of *L. boulardi*, *L.heterotoma* and *L. clavipes*, we used the same LbFV_ORF96 primers as in [22] (ATTGGTGAAATTCAATCGTC and TCATTCATTCGCAATAATTGTG). LbFV_ORF96 was used as a primer because it is the most conserved EVE due to its strong purifying selection compared to the other 12 documented EVEs [22]. They amplified a 411bp internal fragment of the coding sequence. PCR reaction was performed in a 50µL volume containing 10µM primers, 10mM dNTPs, and 0.5uL of Taq DNA polymerase with the following cycling conditions : 95 C 30”,48 C 30”, 72 C 60” (40 cycles). The CO1 marker was correctly amplified in 33 out of 41 specimens, indicating that extraction was satisfying, at least for these specimens. Finally, because we had sometimes several specimens per species, all species but two could be analyzed with at least one specimen.

### Genome sampling, assembly quality check

The DNA of single female was extracted using the same protocol from *Rhoptromeris*, *Trybliographa*, *Leptolamina* and *Trichoplasta*. TruSeq Nano DNA (350) Illumina libraries were built and sequenced with 30 Gb per sample 60M(R1+R2) at Macrogen (Amsterdam, Netherlands). The paired-end reads (2x150bp) were cleared from duplicates using SuperDeduper v1.3.0 [62] (-f nodup), and quality trimmed using Fastp v0.22.0 (–cut_tail –length_required 100 –correction). The assembly was done using Megahit v1.2.9 [50] (–kmin-1pass), a scaffolding step was done using SOAPdenovo-fusion (-D -s) and we obtained a scaffolded homozygous genome assembly using the Redundans pipeline [65] (default parameters). The genome assembly quality check was done using the BUSCO pipeline v 5.3.0 [74] (-m genome) on both Hymenoptera and Arthropoda databases, and assembly statistics were computed using Quast [38] (default parameters) (see genome statistics and accessions on TableS1.

### LbFV-like ORF homology sequence research

We screened all Cynipoidea genomes for the presence of ORFs from LbFV-like virus genomes. The sequence homology search was performed using the Mmseqs2 search algorithm [75] (evalue max = 0.001 -s 7), using as queries all the predicted proteins from the *Naldaviricetes* (see details in [36]) and as database the 11 Cynipoidea genomes. In order to infer the phylogenetic relationships between the endogenized sequences and free-living viruses, we first sought to gather the homologous ORFs between them. To do this, we performed a blastp between all ORFs with the mmseqs2 search algorithm (min bit = 50, min evalue = 1e-04). We then formed clusters between all genes that had at least one homology with one of the members, following the same pipeline used in [36].

### Endogenization arguments

A way to rule out contaminating scaffolds was to look for the presence of insects genes along the scaffolds containing candidate EVEs, assuming that the presence of several insect genes in a viral scaffold is unlikely (except for specific giant viruses or specific insect genes such as apoptotic genes). Eukaryotic genes research was made using the Metaeuk easy-predict workflow [49] (default parameters) followed by the taxtocontig workflow that allows to assign taxonomic labels to the predicted MetaEuk proteins. For each contig, we adopted a majority voting strategy among the taxonomically labeled protein to assign taxonomy to the contig. As an example, a contig was tagged as “Eukaryota origin” if at least 50% of the labels assigned to the encoded proteins were “Eukaryota”.

We also used the presence of transposable elements in contigs as a marker of its eukaryotic nature. Indeed, so far, very few viral genomes have been shown to contain transportable elements [55, 31, 33, 32, 51]. Thus, the probability for a viral EVE to be flanked by a TE is low, even more so if the number of flanking TEs is greater than 1. These results suggest that TE insertions rarely reach high frequencies in viral populations, due to the fact that the majority of endogenizations of TEs in viral genomes are deleterious and are quickly eliminated by selection [33, 32]. We thus looked for TE sequence homology in the scaffolds harboring candidate EVEs (see details in M.M Transposable elements’ detection and analysis) and identified the scaffolds presenting one or more TE.

### LbFV-like EVEs phylogenies and Event assignations

To reconstruct the phylogenetic history of each EVE originating from Filamentoviruses (n=25), we first aligned the protein sequences using ClustalO (v1.2.4)[73] and then inferred the phylogenies using Iqtree (v2.1.2) [56] (options :m MFP -alrt 5000 -bb 5000). The phylogenetic tree was then used to infer endogenization events involving Eucoilini+Trichoplastini species. It is expected that such events will be represented by a monophyletic clade of related wasp species nested within a clade of Filamentoviruses. Within each phylogenetic tree, we grouped all Eucoilini+Trichoplastini wasp species into a single event, if they formed a monophyletic clade with a bootstrap score greater than 80.

### Aggregation of different EVEs in a single endogenization event

In case multiple EVEs arrived together into the ancestral wasp genome, we expect to find signs of shared synteny between descendant species. To uncover this, we conducted the equivalent of an all vs all TblastX (Mmseqs2 search –search-type 4, max Evalue =1e-07) between all the candidate loci within a putative event (deduced from the phylogenetic inference), and then looked for hits (HSPs) between homologous EVEs around the insertions. Because it is possible to find homology between two genomic regions that do not correspond to an orthology relationship, for example because of the presence of conserved domains, we had to define a threshold to identify with confidence the orthology signal. We therefore conducted simulations to define this value, based on the well-assembled genome of *Cotesia congregata* (GCA_905319865.3) by simply performing the same all vs all blast analysis against itself (as if the two species considered had the same genome). Based on this, we defined two types of simulated EVEs, (i) independently endogenized EVEs in the genomes of the two “species”. This is simply simulated by randomly selecting two different regions in the genomes, and (ii) a shared simulated EVE that was acquired by their common “ancestor”. This is simulated by selecting the same random genomic location in both “genomes”. We then counted the total length of the HSPs found around the simulated insertions all along the corresponding scaffold (i and ii). As the result will obviously depend on scaffold length, we performed these simulations on several scaffold lengths (100000000bp, 10000000bp, 1000000bp, 100000bp and 10000bp). We conducted 500 simulations in each scenario, and we measured the cumulative length of homologous sequences, where homologous sequences are defined by sequences having a bit score > 50. We then defined a threshold for each window’s size in order to minimize for the false-positive (FP) and maximize true-positifs (TP) (thresholds 100000000bp = 172737bp (FP = 0.012, TP= 0.922); 10000000bp = 74262 bp (FP= 0.012, TP=0.878); 1000000bp = 21000 bp (FP=0.014, TP=0.28); 100000bp = 1332 bp (FP= 0.012 TP= 0.198) and 10000bp = 180 bp (FP=0.008, TP= 0.208)). This way, the EVEs could be “aggregated” into single events.

## Supporting information

FiguresS13

## Acknowledgment

This work was performed using the computing facilities of the CC LBBE/PRABI. We thank Denis Poinsot for providing specimens of *Trybliographa sp.* as well as Anne-Marie Coretesor for allowing us to consult the PhD manuscript of Nabila Kacem - Haddj El Mrabet.

## Data accessibility

All scripts used in this paper can be found on the GitHub page: https://github.com/GrendelAnonymous/Viral_domestication_Eucoilini_sup All information regarding loci studied and predicted from this article can be found in TableS2.

All Eucoilini genome assemblies as well as raw data can be found under the Bioproject ID number : PRJNA831620.

## Author contributions

B.G. : Conceptualization; Data curation; Formal analysis; Investigation; Methodology; Project administration; Resources; Software; Validation; Visualization; Writing – original draft; Writing – review editing; J.Vo. : Resources; Writing – review editing; R.P. : Resources; Writing – review editing; J.H. : Resources; Writing – review editing; M.L.B. : Resources; Writing – review editing; J.Va. : Methodology; Resources; Validation; Visualization; Writing – review editing.

## Supplemental informations

The following section is a translation of the original PhD thesis made by Nabila Kacem - Haddj El Mrabet in 1999 related to the identification of virus-like structures (VLS) in *Trybliographa rapae*.

### Introduction

While ectoparasitoids behave as micropredators consuming their host from the outside, endoparasitoids immersed in the host’s internal environment are exposed to immune reactions triggered by their infestation. Endoparasitoid Hymenoptera have a variety of strategies for thwarting the host’s response to parasitoid eggs and larvae. These strategies range from oviposition behavior, enabling the egg to be deposited in areas of least host reaction [41, 35, 87], to the release of immunosuppressive substances or liquid particulate secretions from the adnexal glands of the reproductive apparatus [76, 68, 67, 47]. Viral particle production has been described in the ovarian calyx of the Braconidae [83, 28] and in the follicular cells of the Encyrtidae *Leptomastix dactylopii* (Hymenoptera: Encyrtidae) [5] and in the venom gland reservoir in the Eucoilidae *Leptopilina boulardi* and *Leptopilina heterotoma* (Hymenoptera: Figitidae) [68, 67, 72, 25]. Parasitoids and viruses co-evolve [89]. *Trybliographa rapae* (Hymenoptera: Figitidae) is a larvophagous endoparasitoid of the cabbage maggot *Delia radicum* (Diptera: Anthomyiidae) in which we have never found any evidence of encapsulation, suggesting the existence of a protective system. This work, using scanning and transmission electron microscopy techniques, studies the tracking of the *T. rapae* egg from the genital tract to 96 h after oviposition in the *D. radicum* larva, as well as the ultrastructure of the adnexal glands of the female genital tract.

### Materials and methods

The Breton strain of *T. rapae* was reared on *D. radicum* larvae infesting Cruciferae roots using the technique described by [59]. Two other strains, one from Quebec and reared in the laboratory, and the other wild strain obtained from Finland, were used. Second-stage host larvae are subjected to parasitism by females for 5 min. This short time allows us to determine the precise time between egg laying and recovery. Ovarian eggs are obtained by dissection of the ovary, and laid eggs by dissection of the parasitized larvae. These dissections are carried out in Ringer’s physiological fluid. The laid eggs observed are 5 min, 24h, 48h, 72h and 96h old.

Samples observed by scanning electron microscopy are fixed with 2.5% glutaraldheyde buffered with sodium cacodylate for one hour, then dehydrated for 10 min in baths of 70°, 80°, 90°, 95°and twice 100°alcohol, followed by 100° acetone. They were then dried using the CO critical point technique, liquid dried on a Balzers CPD10, then metallized with palladium gold using a cathodic sputter and observed on a JEOL JSM 6400. Samples observed by transmission electron microscopy (ovipositor sections, laid eggs and odd and even adnexal glands) are fixed with cacodylate-buffered gluraldehyde 2.5, placed in a 2% buffered osmium tetroxide solution for 1 h and dehydrated with pure acetone. They are embedded in Epon-Araldite. Semi-fine sections are taken on a Reichert OM.U2 ultramicrotome. Sections are stained with 1% toluidine blue and observed by Phillips CM - 12 transmission electron microscopy. The laid egg was also observed 24 hours after oviposition using light microscopy.

### Results

The ovarian wall egg, freed from surrounding follicular cells, shows a regularly and strongly wrinkled exochorion (FigureS11-A). The newly laid egg is completely covered by a dense coating of viscous-pasty particles ranging in size from 0.20 µm to 0.42 um in length (FigureS11-B). Twenty-four hours after oviposition, the hydropic egg has greatly increased in volume as a result of the detachment occurring between oviposition, exochorion and endochorion. The space thus created is filled with fluids from the internal environment of the parasitized larva that has passed through the exochorion. The yolk of the bulbar part of the *T. rapae* pedicellate egg remains contained within the endochorion (FigureS11-C). As the volume of the egg increases, the exochorion unfolds, causing a change in the particle coating (FigureS11-D). The particles, now ranging in size from 0.57 um to 1.14 um, organize themselves into a network, with connecting bridges appearing between the particles in the Breton strain (FigureS11-D), the Quebec strain (FigureS11-E) and the Finnish strain (FigureS11-F). Forty-eight hours after oviposition, the phenomenon of egg hydropy is complete; the particles reach their greatest spread and measure 1.3 µm to 2 µm in diameter (FigureS11-G). In cross-section, the 48h-old particles, which appear hollowed out, are surrounded by a double membrane and rest closely on the exochorion (FigureS11-H). The lumen of the paired gland shows grains of secretion (FigureS11-I). Their size, 2.4 um in diameter, is comparable to that of the particles found on the egg laid 5 minutes ago. These agglomerated secretions are found in the ovipositor canal (FigureS11-J); the cluster slides over the ctenidia carried by the inner face of the valves delimiting the canal. The venom gland (odd-numbered) has three easily distinguishable parts: the distal part is 480 µm long and corresponds to the gland itself, the narrow middle part is 320 um long and corresponds to the canal, and the swollen proximal part or reservoir varies from 320 to 500 µm depending on the quantity of venom stored. The inner surface of the gland displays two types of secretory pores. Type A pores appear on the inner surface in the form of nipples 2 um in diameter, pierced by a central orifice 0.3 um in diameter, which certainly allows secretions to escape (FigureS10-A). Type B pores are found on the entire inner surface in the form of rounded cavities 3.3 um in diameter (FigureS10-B). In cross-section (FigureS10-C), these pores are composed of crypt-like secretory units discharging their products into the matrix-filled glandular lumen. In the distal pores, easily recognizable particles can be found (FigureS10-D). At the same distal level of the gland, higher magnification reveals the isolation and grouping of the constituent elements (FigureS10-E) of the particles (FigureS10-F). These roughly spherical particles vary in diameter from 0.6 to 1.2 um. In the middle part of the gland, they are well individualized and distinct from their substrate (FigureS10-G). In the proximal part, or reservoir, of the venom gland, they remain unchanged (FigureS10-H). A semi-fine section of the ovipositor stylet shows the presence of these particles in the oviposition canal (FigureS10-I). These particles, whose formation is followed from the apex to the venom gland reservoir, are expelled by the ovipositor; however, they were not found in contact with the laid egg.

### Discussion

The presence of “Virus-Like Particles” has been reported in many Braconidae: *Biosteres longicaudatus* [47], *Microplitis mediator* (Walker) [83], *Microplitis croceipes* (Cresson) [40], *Apanteles melanoscelus* [78], *Apanteles congregatus* (Say) [80], *Apanteles glomeratus* [43], *Apanteles hyphantriae* (Riley) [79], *Cardiochiles nigriceps*, *Microplitis croceipes*, *Chelonus texanus* and *Cotesia rubecula* [3]. Some Ichneumonidae, including *Venturia canescens* [7, 27], *Hyposoter exiguae* [45] and a single Encyrtidae, *Leptomastix dactylopii* [5], also possess virus particles. Virus particles are always replicated in Ichneumonidae and most often in Braconidae at the ovarian calyx. In the latter, replication of two types of particles in *Biosteres longicaudatus* [47] takes place in the adnexal glands. Particle replication occurs in the follicular cells of the Encyrtidae *Leptomastix dactylopii* [5] and in the epithelial cells of the adnexal glands associated with the female reproductive tract in the Figitidae *Leptopilina boulardi* and *Leptopilina heterotoma*. Several types of virus are involved in suppressing the host’s immune system. Baculoviruses are found in the Braconidae; these membrane-enveloped virions are cylindrical nucleocapsids 40 nm in diameter, containing a genome composed of a circular DNA strand that replicates in the cell nucleus. PolyDNA viruses are highly unusual viral entities [79, 28, 76]. Their genome is made up of several dozen circular double-stranded DNA molecules. Other so-called rod-shaped particles are assimilated to VLPs [44, 81, 61, 4, 77, 2]; they differ in size from the VLPs found in the accessory glands of *Biosteres longicaudatus* [47, 26]. Our work shows the formation of 2 types of particles in *T. rapae*, formed in two different sites: the odd gland (venom gland) and the even gland (uterine gland). These 2 particle types are found in the ovipositor canal. The type produced by the paired gland (uterine gland) is easily found on the laid egg of all three strains of *T. rapae*.

It therefore provides close protection for the egg. The type produced by the odd-numbered gland, on the other hand, is not found on the egg, suggesting that these particles are dispersed within the host larva. Since no evidence of host aggression or encapsulation was observed on several hundred laid eggs, we attribute to these most likely viral particles an immunosuppressive role in host reactions. *T. rapae* thus appears to be the first Figitidae and the second parasitoid species to possess two identified types of particles produced by the even and odd glands of the reproductive tract. In *Biosteres longicaudatus* (Hymenoptera: Braconidae), both types of particle originate from the filaments of the accessory gland and Dufour’s gland, considered to form part of the venomous apparatus [47]. The comparison of the various situations described in the field of virus-parasitoid interactions calls for a homogenization of the qualification by the various authors of the accessory glands in general, in order to better understand the strategies put in place by co-evolution, and which may eventually contribute to the ongoing systematic revision of the species concerned.

### Supplementary figures

**Figure S1.**
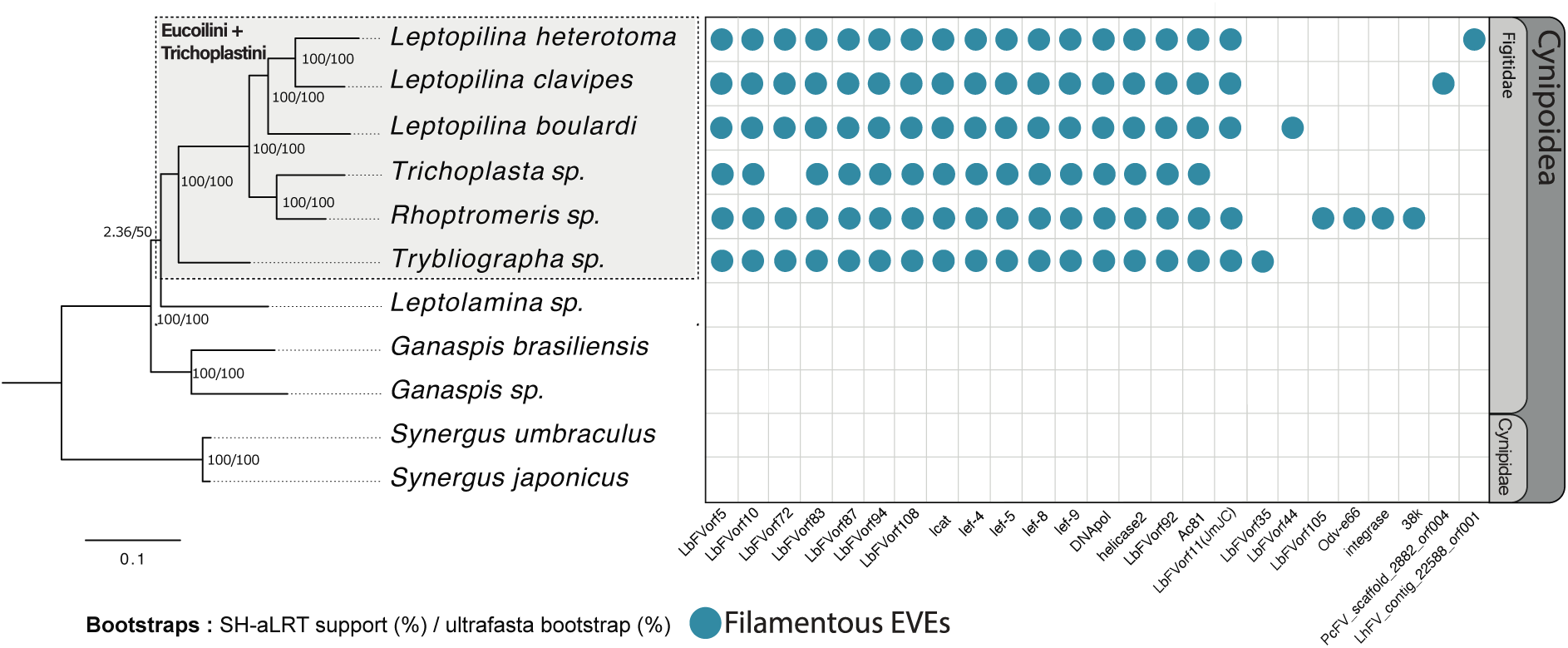
Filamentous EVE distribution among Cynipoidea species. The phylogeny of the Cynipoidea has been estimated using 1,000 Busco genes. Confidence scores (aLRT%/ultra-bootstrap support%) are shown at each node. Each row represents a Cynipoidea species, whereas each column represents a filamentous EVE. Blue circles indicate the presence of EVE in a genome, whereas white boxes indicate its absence.

**Figure S2.**
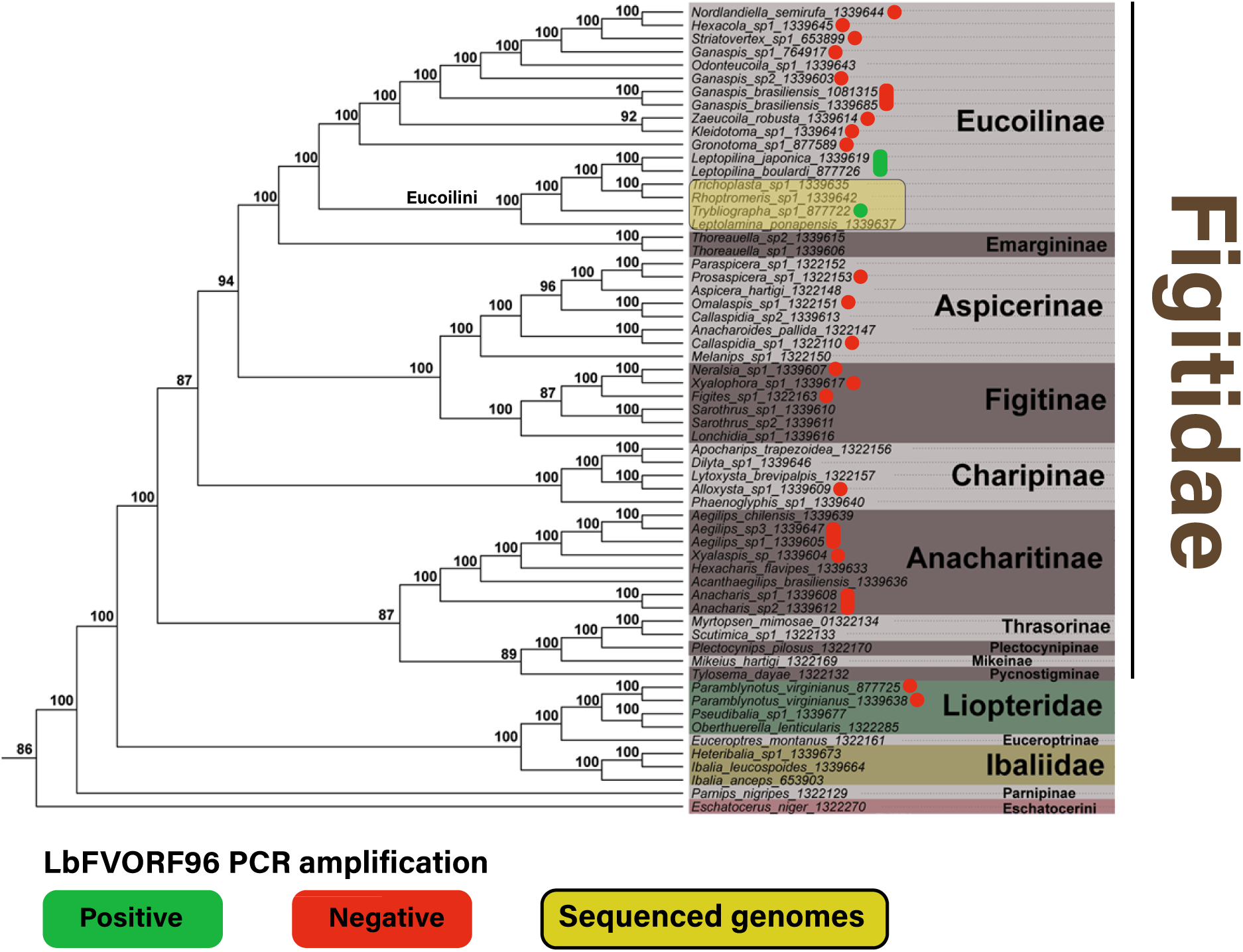
Figitidae phylogenetic tree with all sampled species. The figure is modified from the previous work of [11]. Positive PCR amplification of ORF96 for each genus is displayed in green, while negative in red. This only concerns genus in the phylogeny, it does not mean the specific taxa as been tested. To see the specific species tested, please refer to the (FigureS3). Genus that were newly sequenced and assembled are in yellow box, this includes *Leptolamina*, *Trybliographa*, *Trichoplasta* and *Rhoptromeris*.

**Figure S3.**
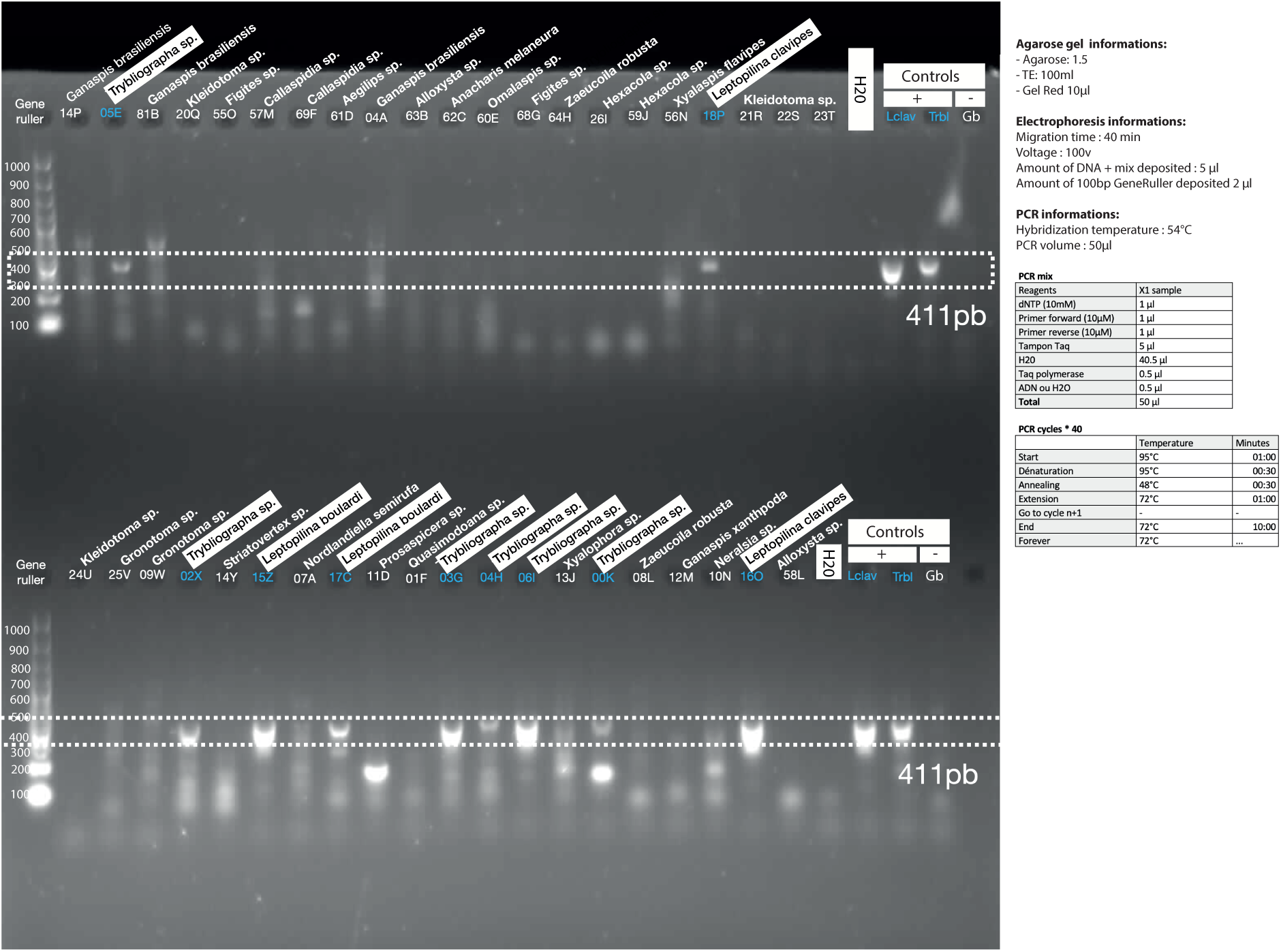
ORF96 PCR results. Positive PCR amplification of ORF96 is displayed by a blue number and a white box. The size of amplicons can be found thanks to PCR ladder on the left side of the figure. The expected ORF96 amplicon size was 411bp and is delimited with the doted white rectangles. Control abbreviations : Gb=*Ganaspis brasiliensis*, Lclav=*Leptopilina clavipes*, *Trbl=Trybliographa*. Information about the PCR mix, programs electrophoresis concentration are displayed next to the electrophoresis picture,

**Figure S4.**
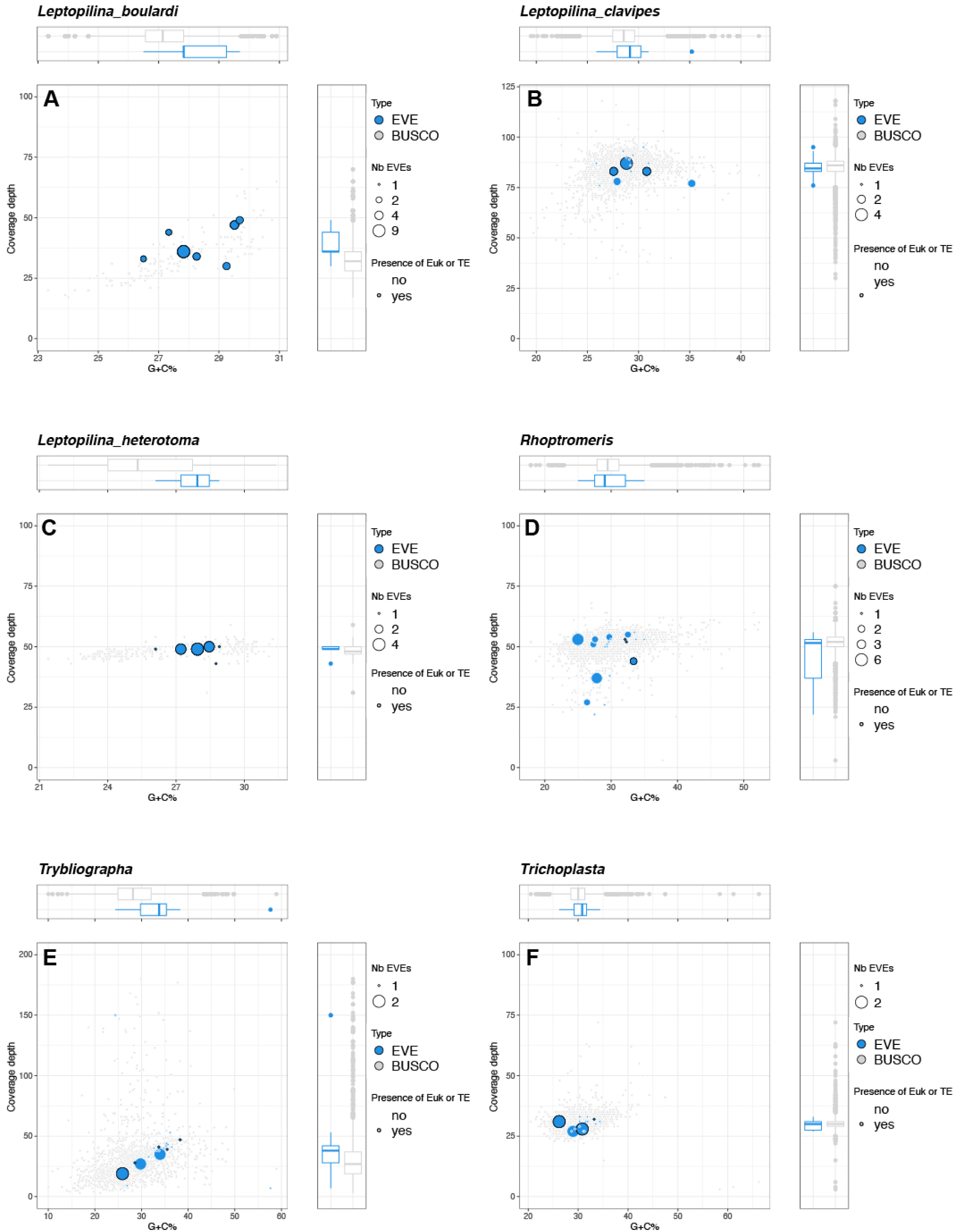
Coverage and G+C% content of scaffold harboring filamentous EVEs. General features of scaffolds containing single copy universal arthropod genes (BUSCO gene set, in grey). Scaffolds containing filamentous virally-derived loci are in blue. The size of the dots corresponds to the number of candidate EVEs inside the scaffold. The dots circled in black correspond to scaffolds that contain one or more eukaryotic genes and/or one or more repeat elements. (A) *L. boulardi*; (B) *L. clavipes*; (C) L. heterotoma; (D) *Rhoptromeris sp*; (E)Trybliographa; (F) *Trichoplasta*.

**Figure S5.**
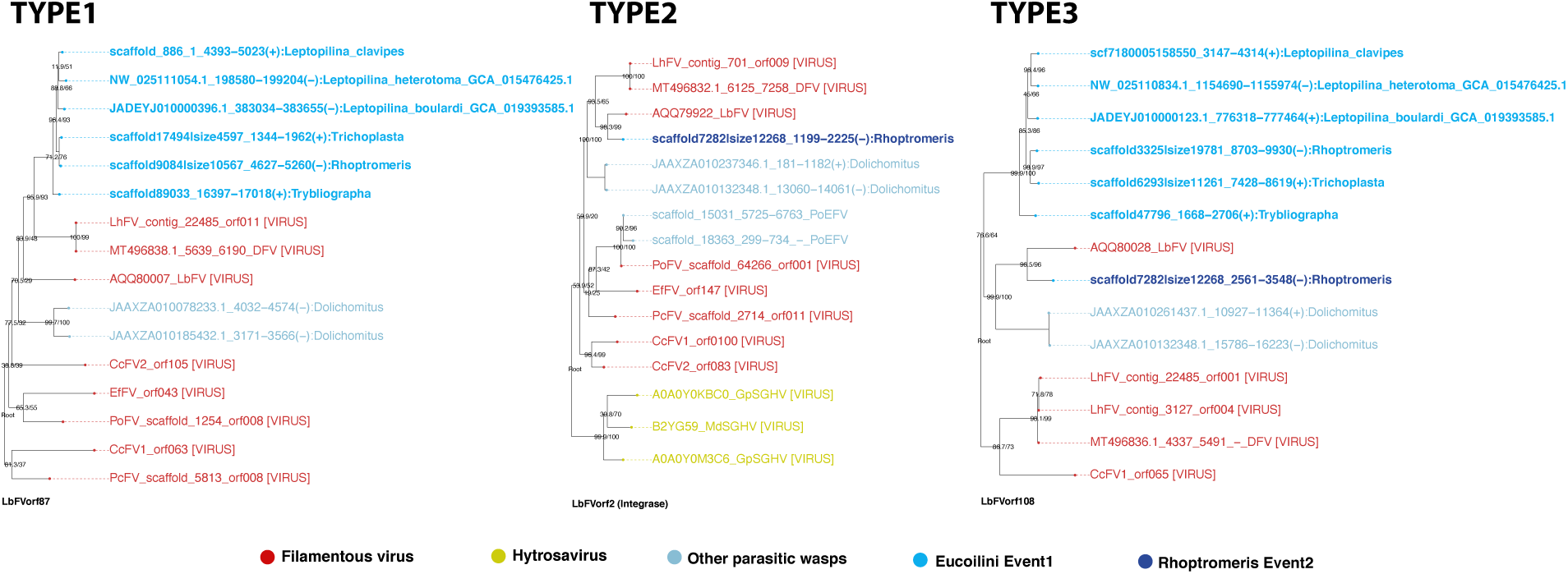
Example of individual gene phylogenies observed. Phylogenies include sequences from free-living viruses such as Filamentoviruses (in red) and Hytrosavirus (in yellow) and sequences from parasitic wasps in blue tones. EVEs assigned to the first ancestral endogenization event within the Eucoilini tribe are depicted in light blue, while the EVEs assigned from the recent endogenization event in the *Rhoptromeris* genome are depicted in dark blue. Three types of phylogenies are present in the 25 gene phylogenies of the analysis. The type1 corresponds to a single endogenization event that occurred in the ancestor of all Eucoilini species and involving a LhFV-like donor. The Type2 suggests another independent event that occurred in the *Rhoptromeris* branch, involving a LbFV-like donor. The type3 phylogenies suggest that both events occurred.

**Figure S6.**
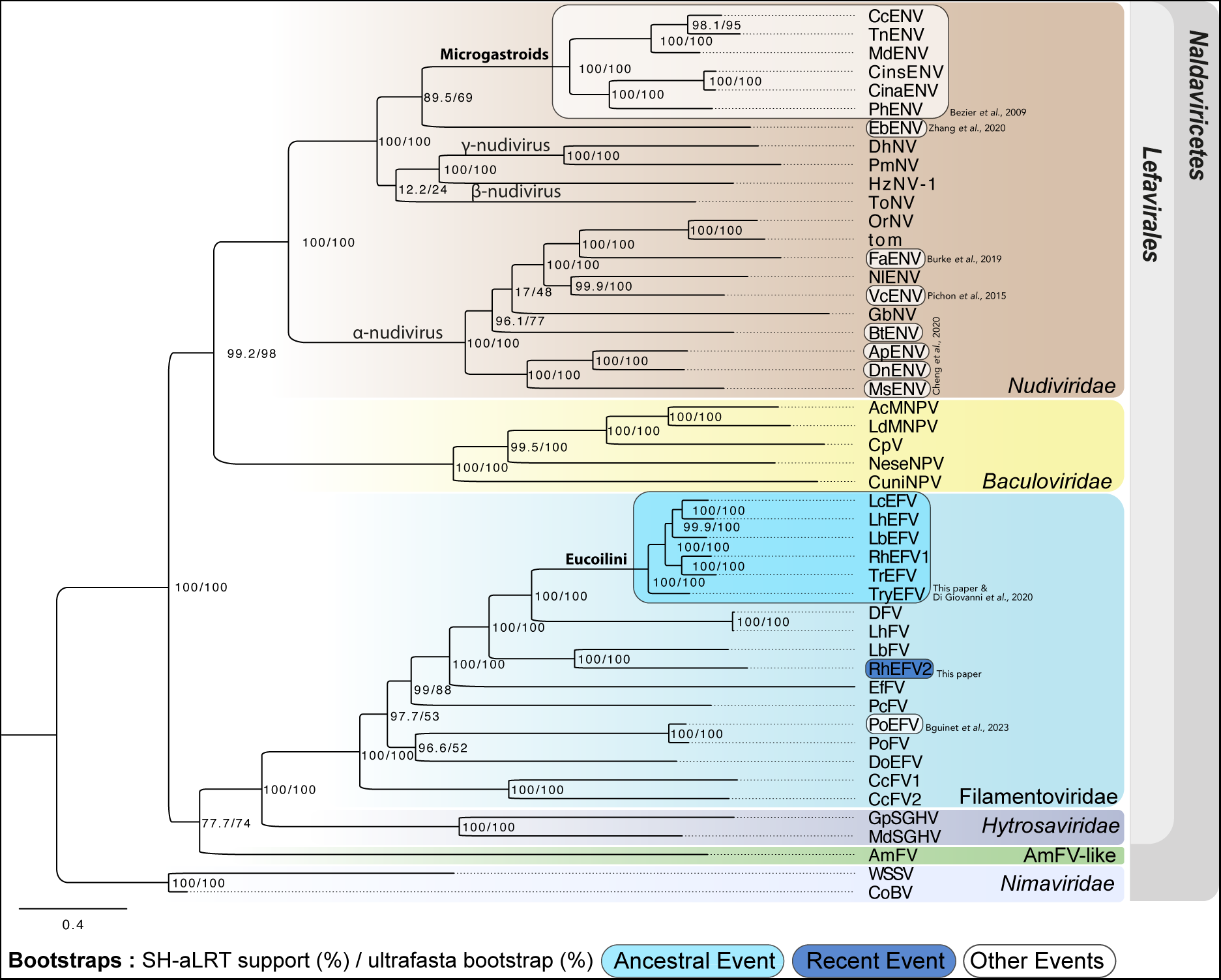
Phylogenetic tree inference of the *Naldaviricetes*. The phylogeny was inferred by maximum likelihood with 113 concatenated amino-acide sequences (25,382 amino-acides sites including only homologous clusters with at least 4 sequences) of 47 viruses. Confidence scores (aLRT%/ultra-bootstrap support%) are shown at each node. The scale bar indicates the average number of amino acid substitutions per site. The phylogeny includes free-living viruses from the families *Baculoviridae* (yellow), *Nudiviridae* (brown), *Hytrosaviridae* (purple), *Nimaviridae* (light purple), Filamentoviridae (light blue) and the AmFV virus (green). All previously documented viral endogenization events are highlighted in white boxes, while the newly described events in this paper are highlighted in blue tones (light blue for the first ancestral event in Eucoilini, and dark blue from the recent endogenization event in the *Rhoptromeris*.

**Figure S7.**
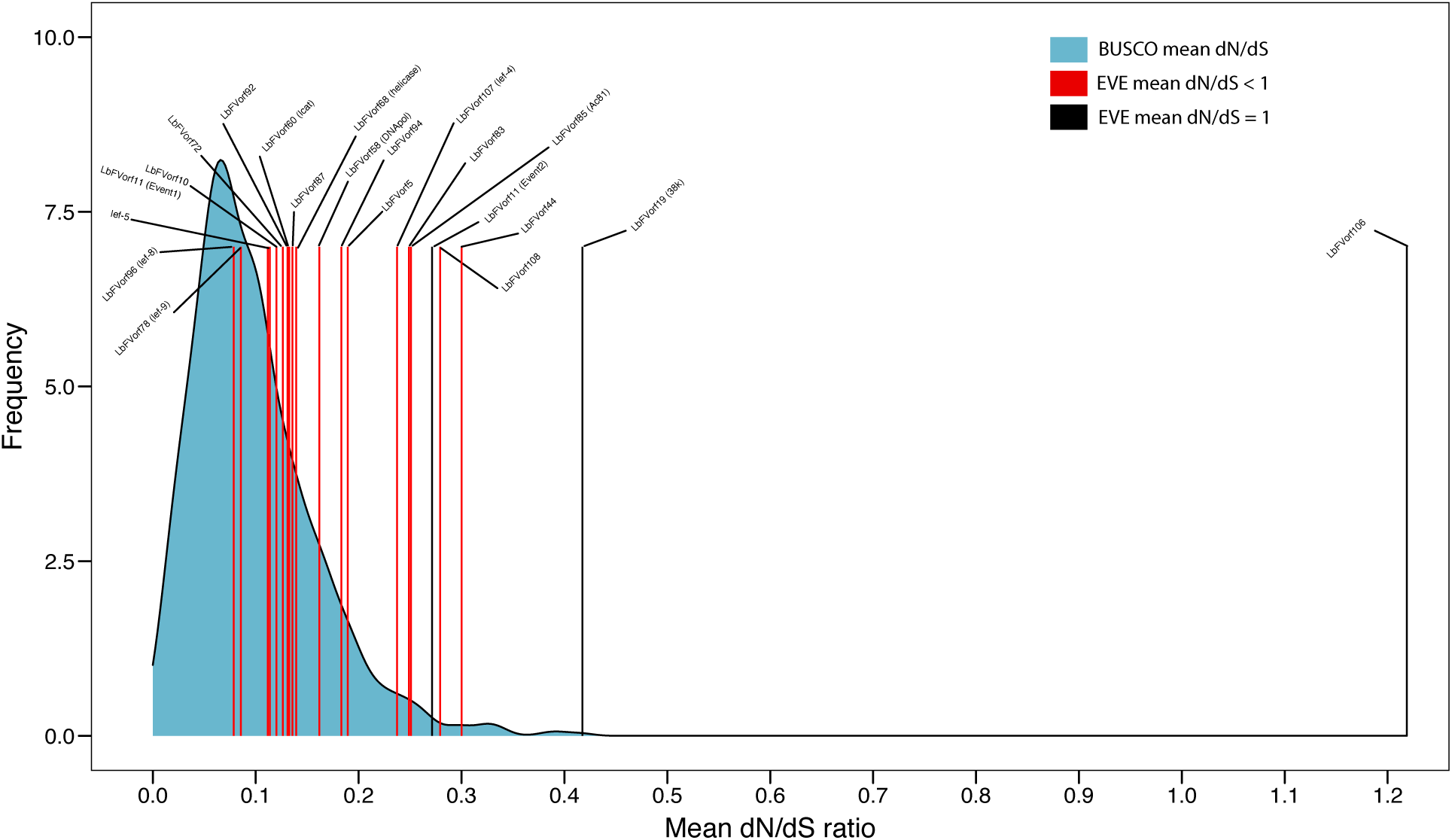
Virally derived genes are under strong purifying selection in Eucoilini wasp genomes. *dN/dS* ratio for a set of 1000 universal arthropod genes (blue density curve) and for 18 virally derived genes found in *Leptopilina*, *Trybliographa*, *Trichoplasta*, and *Rhoptromeris* species (indicated by the red lines). Red lines indicate *dN/dS* significantly below 1 while black lines indicate *dN/dS* =1. The labels above the lines indicate the number of the ORF within the LbFV genome, and the putative protein names are indicated between parenthesis.

**Figure S8.**
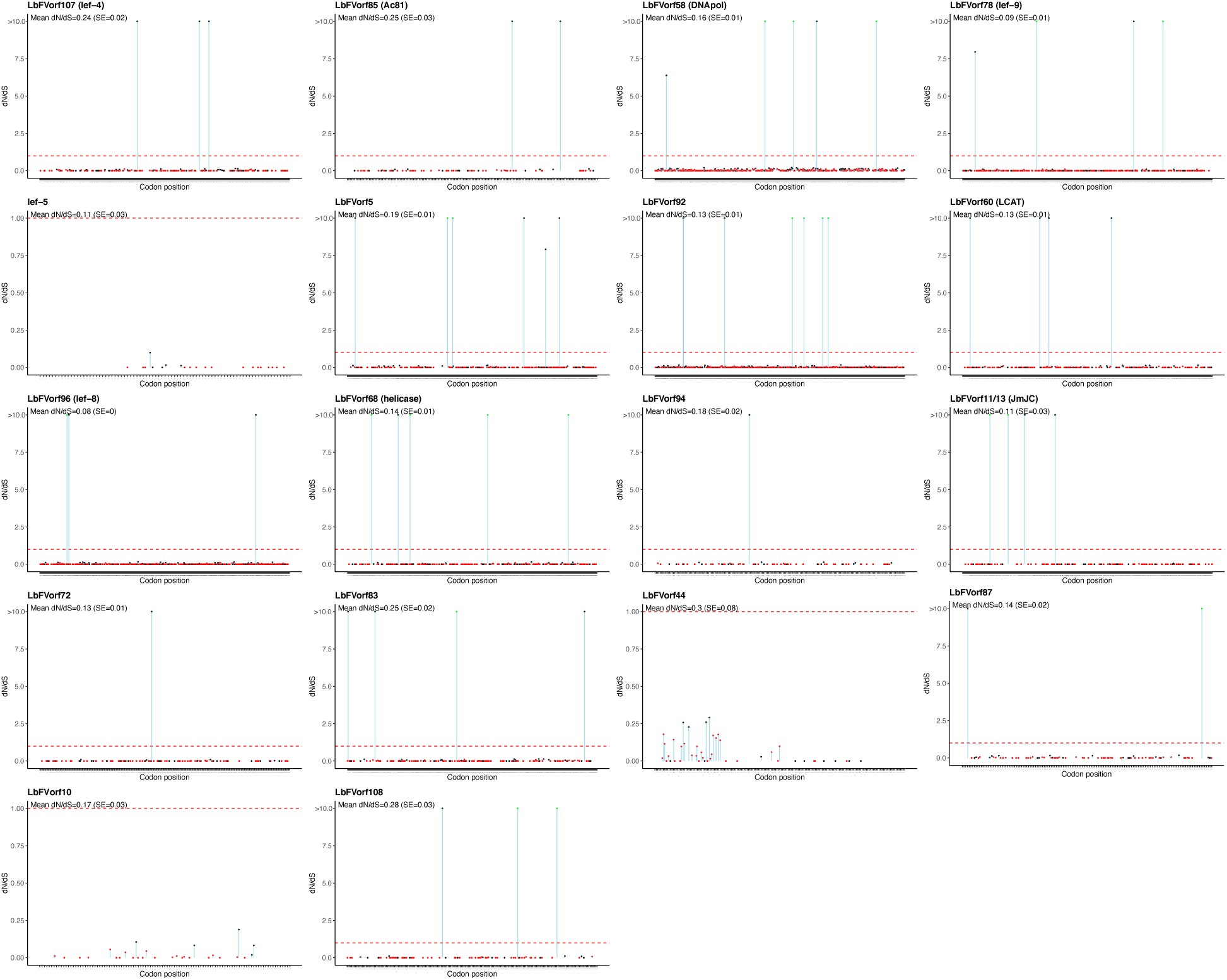
*dN/dS* profiles for the 18 filamentous genes endogenized in Eucoilini wasps. The *dN/dS* values were estimated for each codon position of the endogenous filamentous viral elements using FEL method. Sites inferred to be under positive or negative selection are displayed in green and red respectively (pvalue<0.05), while sites evolving under neutrality are displayed in black.

**Figure S9.**
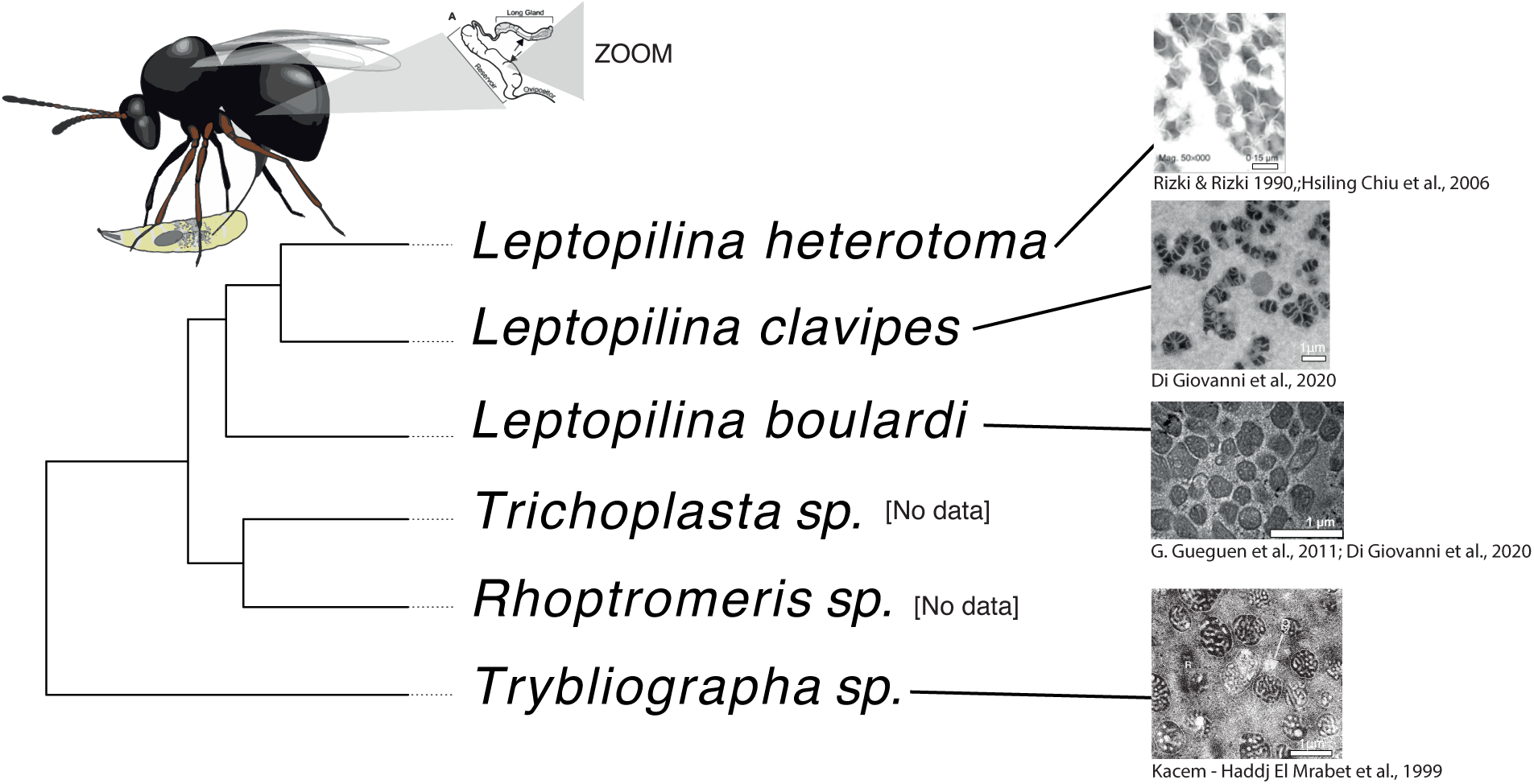
Summary on VLPs production in the venom gland of Eucoilini species. TEM figures are from [19, 22, 34], and K.Haddj E PhD 1999 respectively. To our knowledge, no data are available for *Trichoplasta* and *Rhoptromeris* genus. The cartoon in the upper left corner illustrates the location of the tissue used for the TEM investigations (reservoir of the venom gland). Cladogram extracted from the phylogeny presented in FigureS1

**Figure S10.**
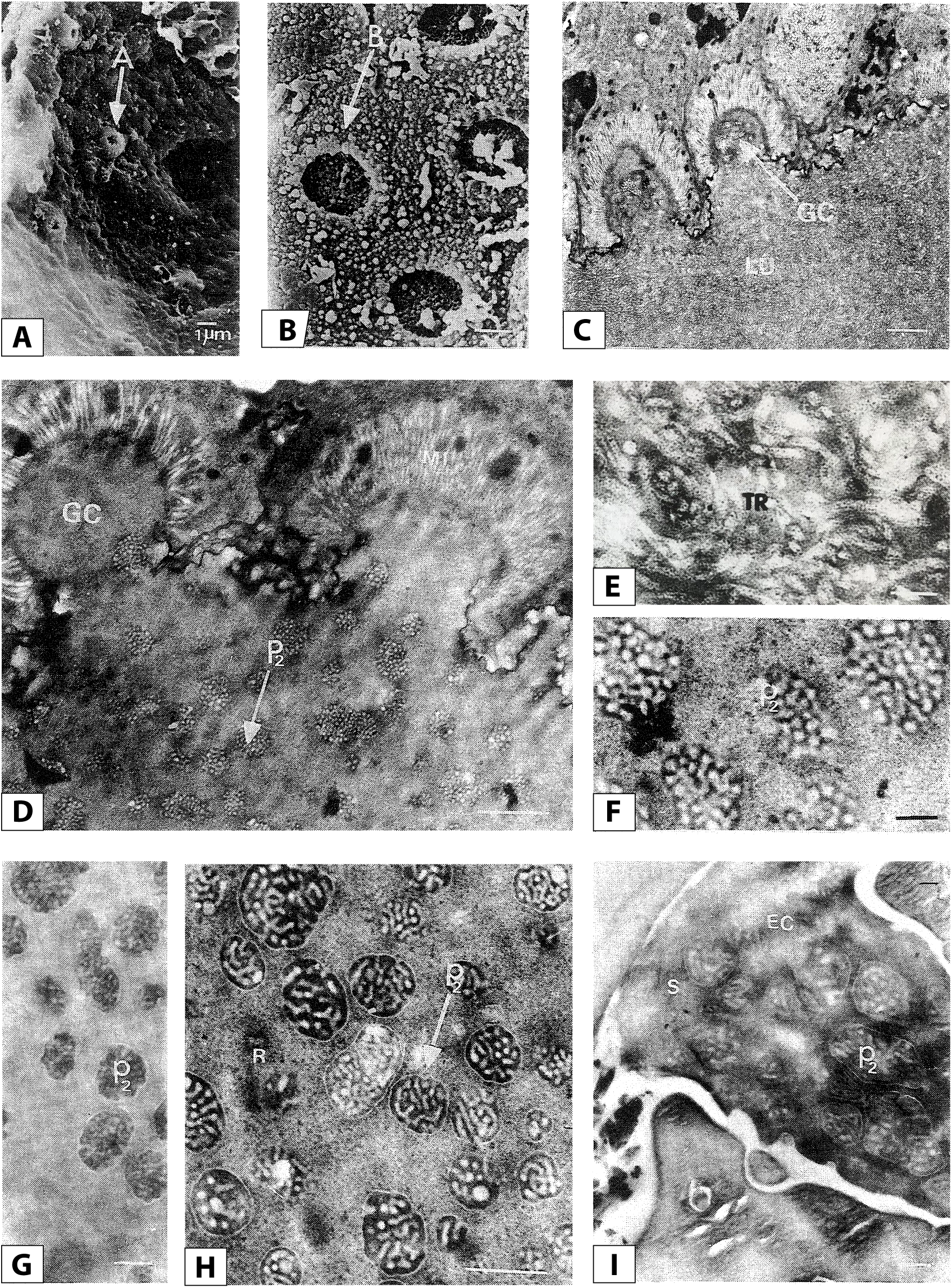
Transmission electron micrographs of sections of *Trybliographa rapae* venom gland. **A** Relief type A pore at the distal end of the inner side of the venom gland, **B** Type B pore at the distal end of the gland, **C** Cross-section of the glandular units at the distal end, the lumen of the gland is filled by a membrane frame, **E** Detail of the membrane frame, **F** Individualization of virus-like particles (VLPs) in the distal part of the odd gland, **G** Presence of virus-like particles in the lumen of the middle part of the venom gland, **H** Appearance of particles in the lumen of the venom gland reservoir, **I** Presence of VLPs in the egg-laying canal. A: type A pore, B: type B pore, EC: oviposition canal, EXO: exochorion, GC: glandular cells, LU: gland lumen, MI: microvilli, p2: type 2 particles, R: reservoir, S: secretions, TR: frame. Pictures from the PhD thesis manuscript of Nabila Kacem-Haddj El Mrabet, 1999

**Figure S11.**
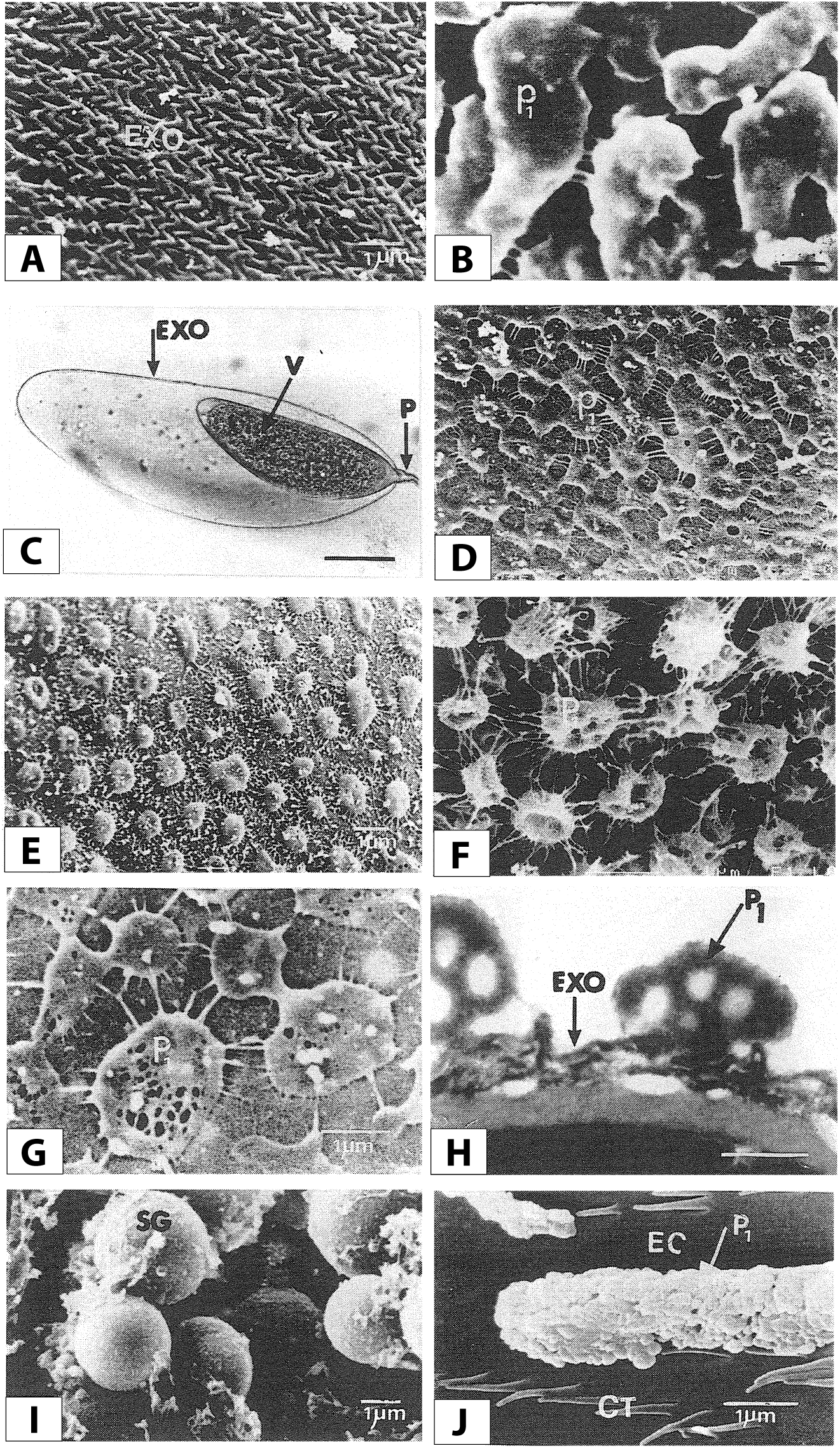
*Transmission electron micrographs of sections of Trybliographa rapae uterin gland*. **A** The exochorion of the ovarian egg, **B** Coating of the egg surface with particles 5 min after oviposition. The chorion has a dense coating of viscous substance, **C** General appearance of the *T.rapae* egg 24 hours after oviposition; **D** Surface of an egg 24 hours after laying from a Breton strain; **E** Surface of an egg 24 hours after laying from a Quebec strain, **F** Surface of an egg 24 hours after laying from a Finnish strain, **G** Egg 48 hours after laying, **H** Egg 48h after oviposition in semi-fine section showing the viral-like particles found on its surface, **I** Secretion grains of the paired gland, **J** Presence of particles in the middle part of the ovipositor at the level of the oviposition canal lined with ctenidia. CT: Ctenidia, EC: oviposition canal, EXO: exochorion, FC : folded chorion, P: pedicel, p1: type 2 particles, p2: type 2 particles, SG: secretion grains. Pictures from the PhD thesis manuscript of Nabila Kacem-Haddj El Mrabet, 1999

**Figure S12.**
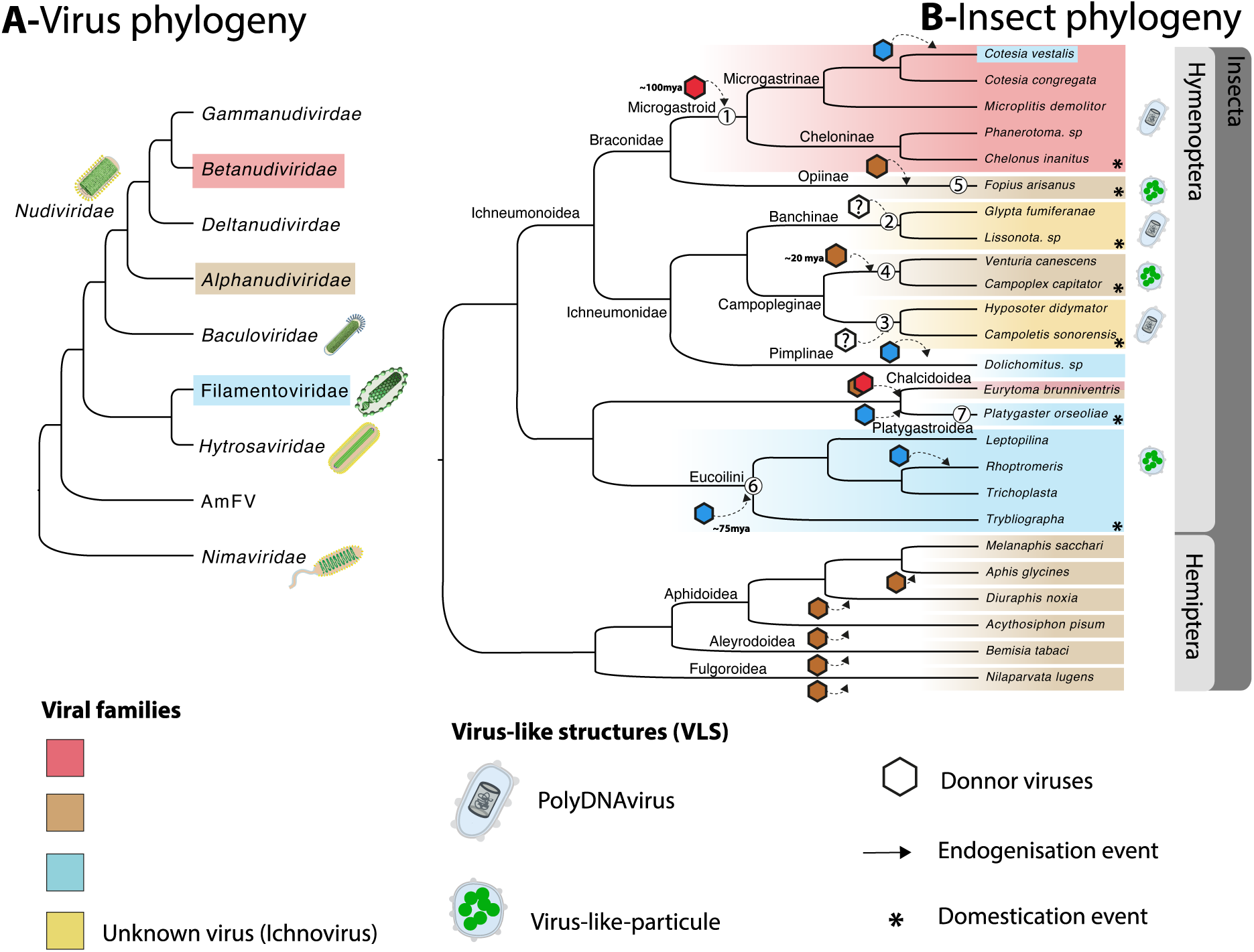
Summary of endogenized and domesticated viral elements of dsDNA viruses in arthropods. **A** - Phylogeny of viruses of the class *Naldaviricetes*, with the virus clades involved in endogenisation phenomena coloured. **B** - Phylogeny of insects including Hymenoptera and Hemiptera having integrated and/or domesticated genetic material from viruses of the class *Naldaviricetes*. Event (1) corresponds to the domestication event of a betanudivirus in the common ancestor of Microgastroids, about 100 million years ago, which allows the production of PDVs [9]. Event (2) corresponds to the domestication of an unknown virus in the ancestor of at least two species of the Banchinae family which is independent of event (3) in which another unknown virus was domesticated in species of the Campopleginae family. Both allow the production of PDVs [8, 15, 48]. Event (4) corresponds to a domestication event of a betanudivirus that took place about 20 million years ago and is found in the genome of *V.canescens* and *C.capitator* and allows the production of VLPs [63] (PhD thesis of Alexandra Cerqeira de Araujo). Event (5) corresponds to a domestication event of a betanudivirus that took place in the genome of *F.arisanus* and allows the production of VLPs [16]. Event (6) corresponds to a domestication event of a Filamentoviridae close to LhFV that occurred around 75 million years ago in part of the Eucoilini species described in the present paper, allowing the production of VLPs at least in *L.boulardi* and *T.rapae*. Another independent event involving a Filamentovirus close to LbFV also occurred within the *Rhoptromeris* genus. Event (7) corresponds to the endogenization and probably domestication of a Filamentoviridae closely related to PoFV that occurred recently within the *P.orseoliae* genome [37]. All the other unnumbered events correspond to independent *a priori* events that show no trace of domestication.

