## Supplementary material for "Dating the origin of a viral domestication event in parasitoid wasps attacking Diptera": FigureS14: FigureS14_ALL_Eucoilini_filamentous_phylogenies.pdf

Figure S8-1: Cluster194 – LbFVorf5

**Phylogeny of type 3**

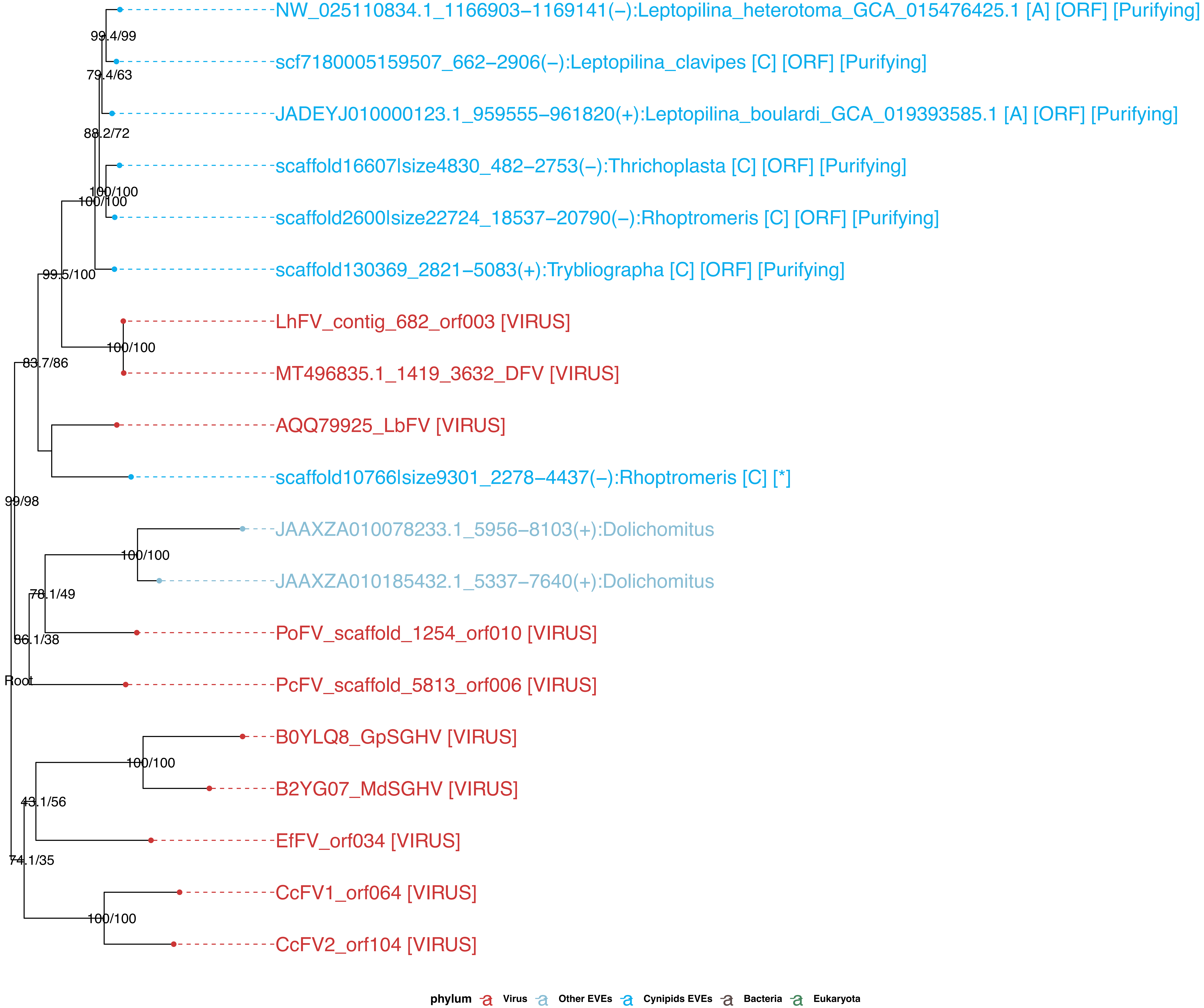

**Figure S8-2: Cluster279 – LbFVorf10**

**Phylogeny of type 3**

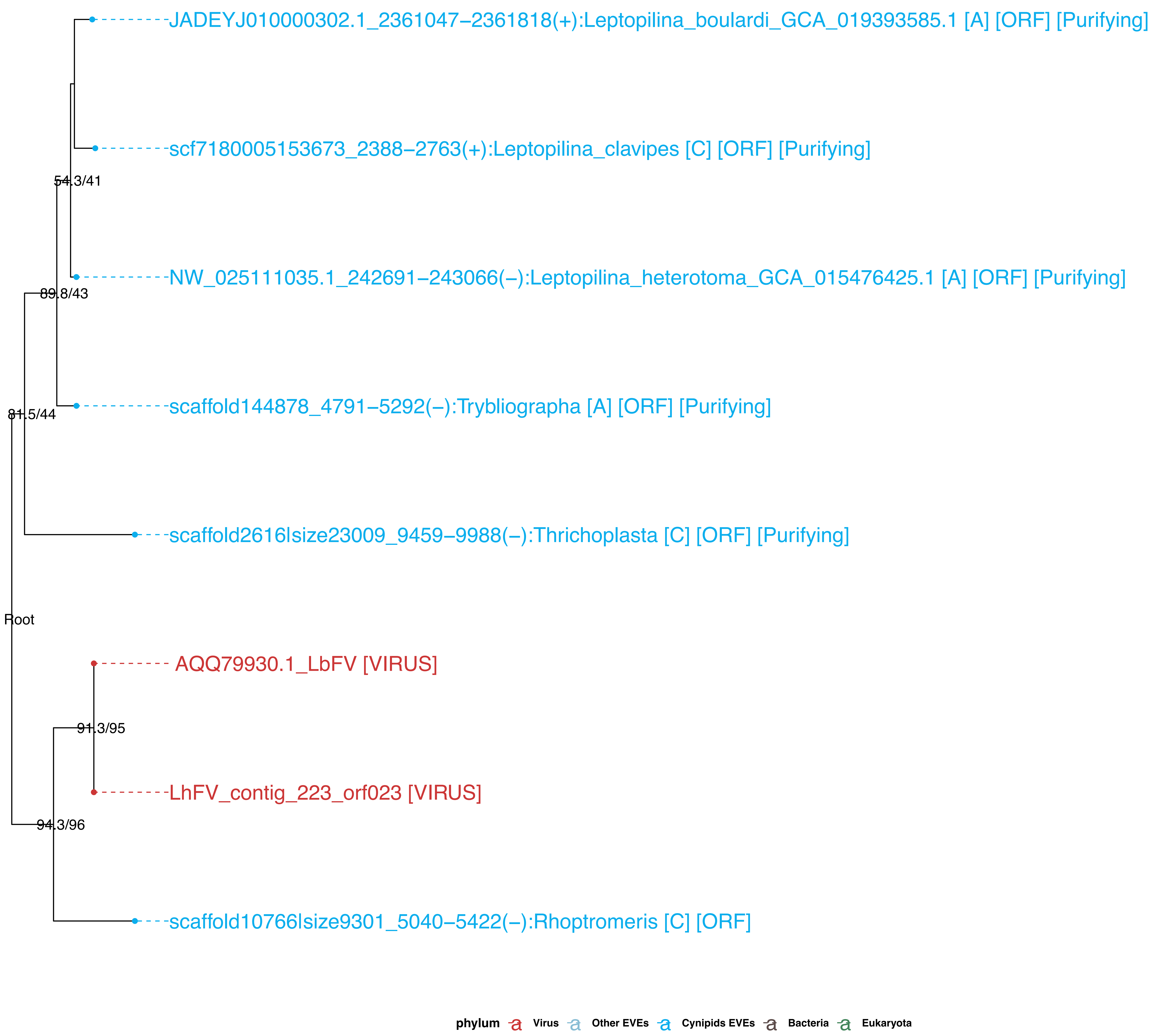

**Figure S8-3: Cluster114_redefined – LbFVorf11 (JmJC)**

### Phylogeny of type 3

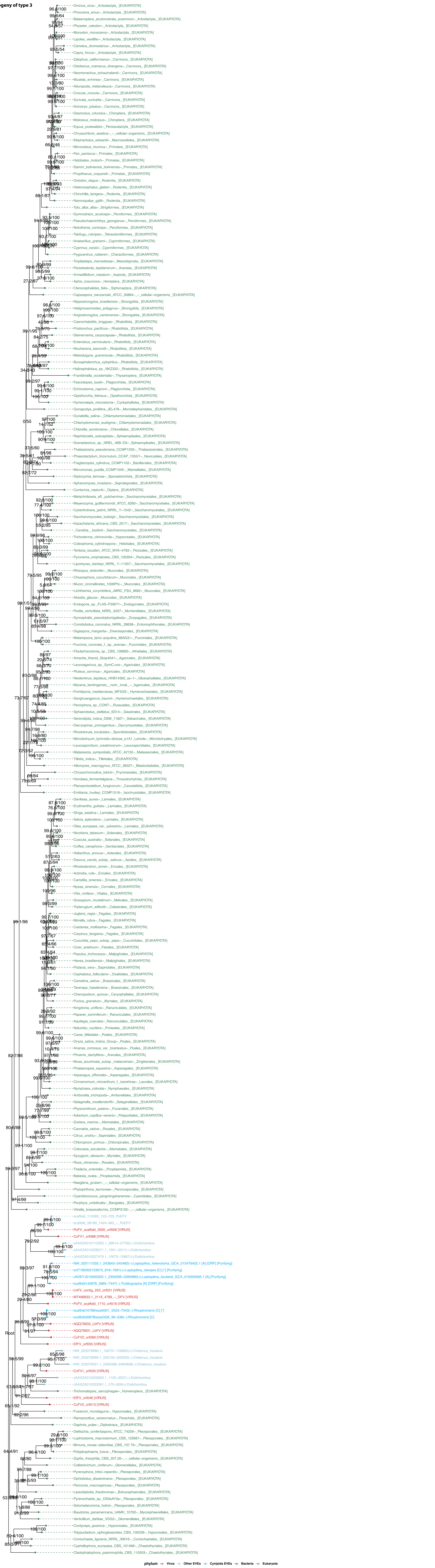

**Figure S8-4: Cluster29_redefined – LbFVorf44**

**Phylogeny of no type**

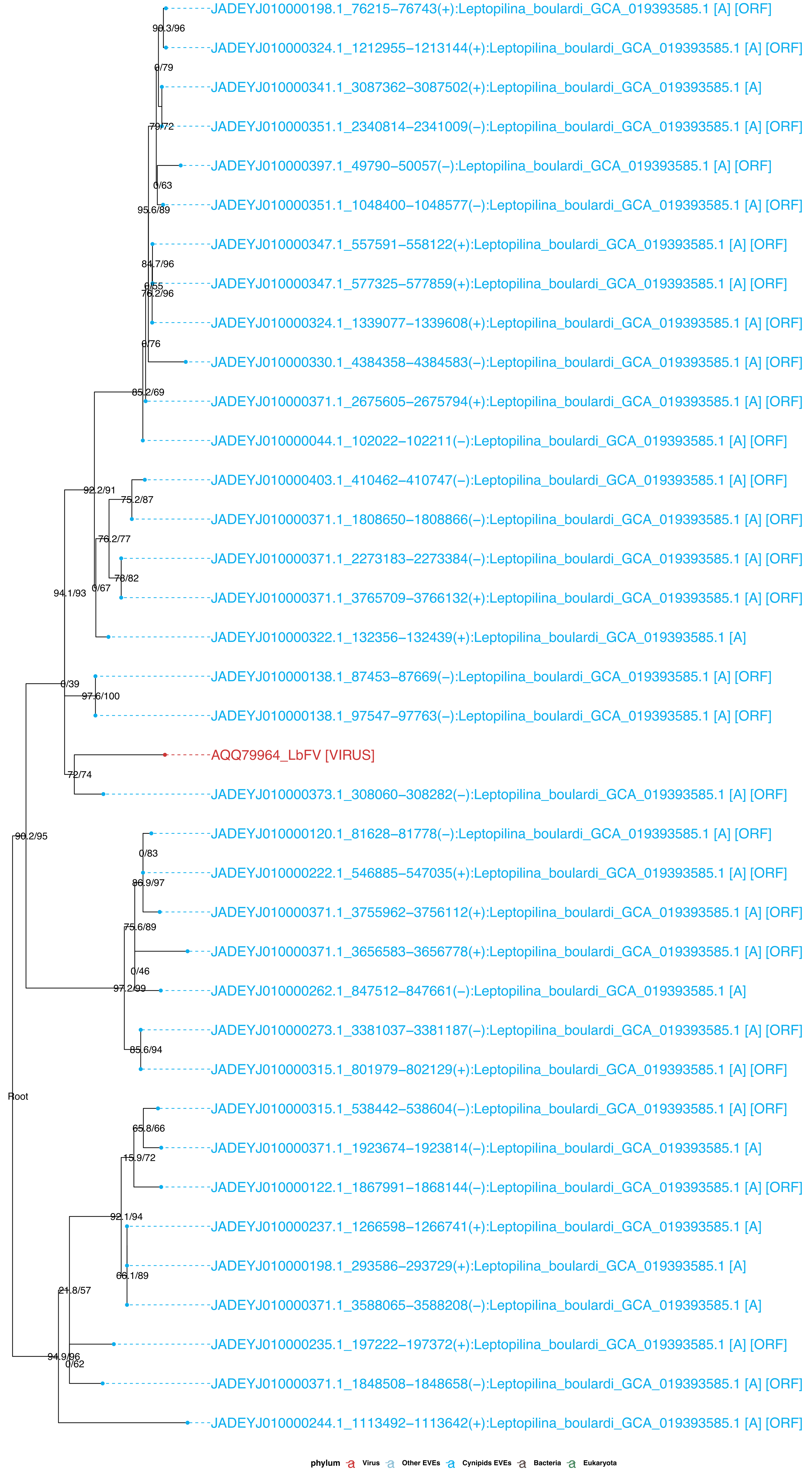

Figure S8-5: Cluster258 – LbFVorf72

**Phylogeny of type 2**

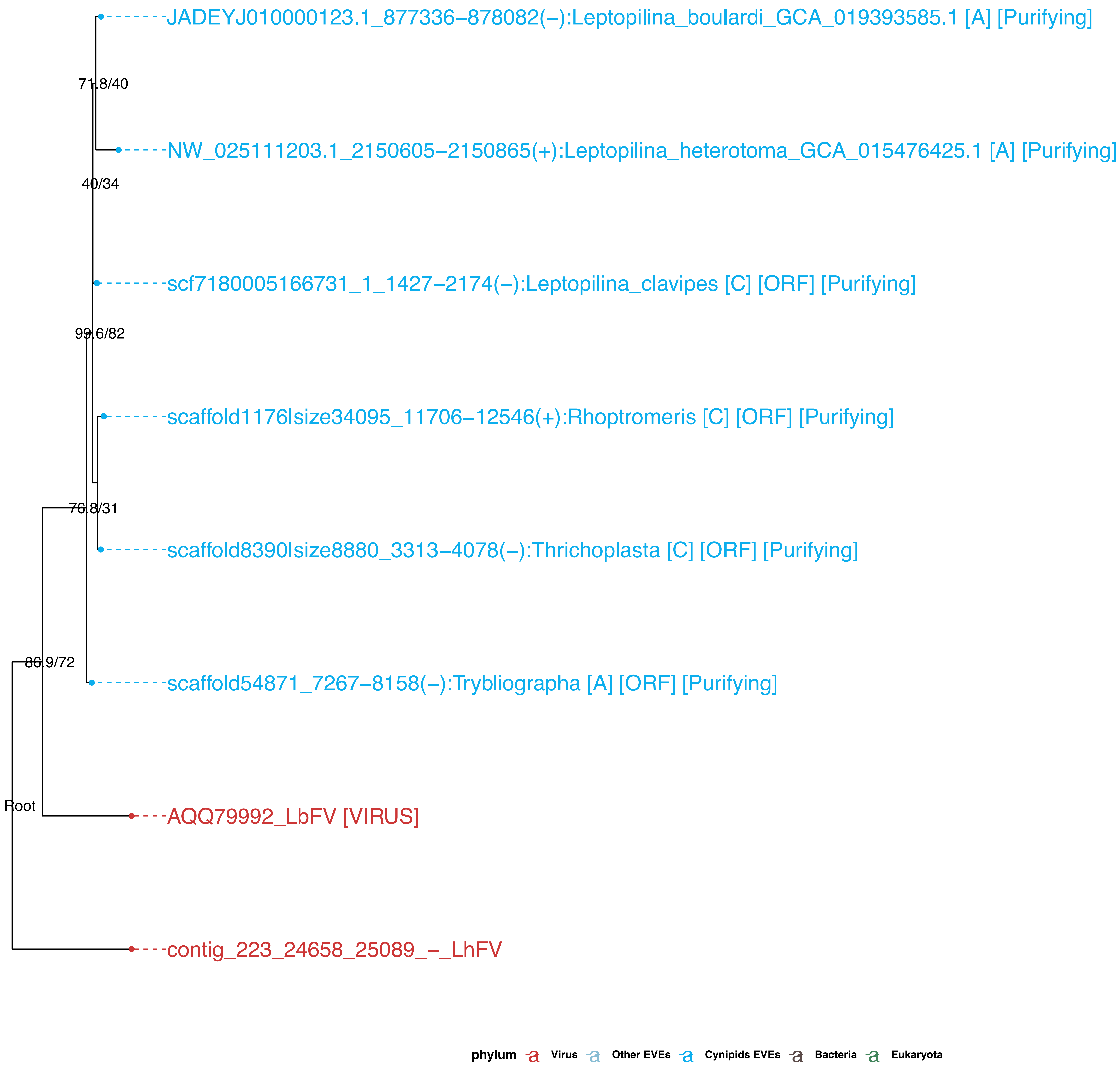

Figure S8-6: Cluster27–Cluster1722 – LbFVorf83

**Phylogeny of type 1**

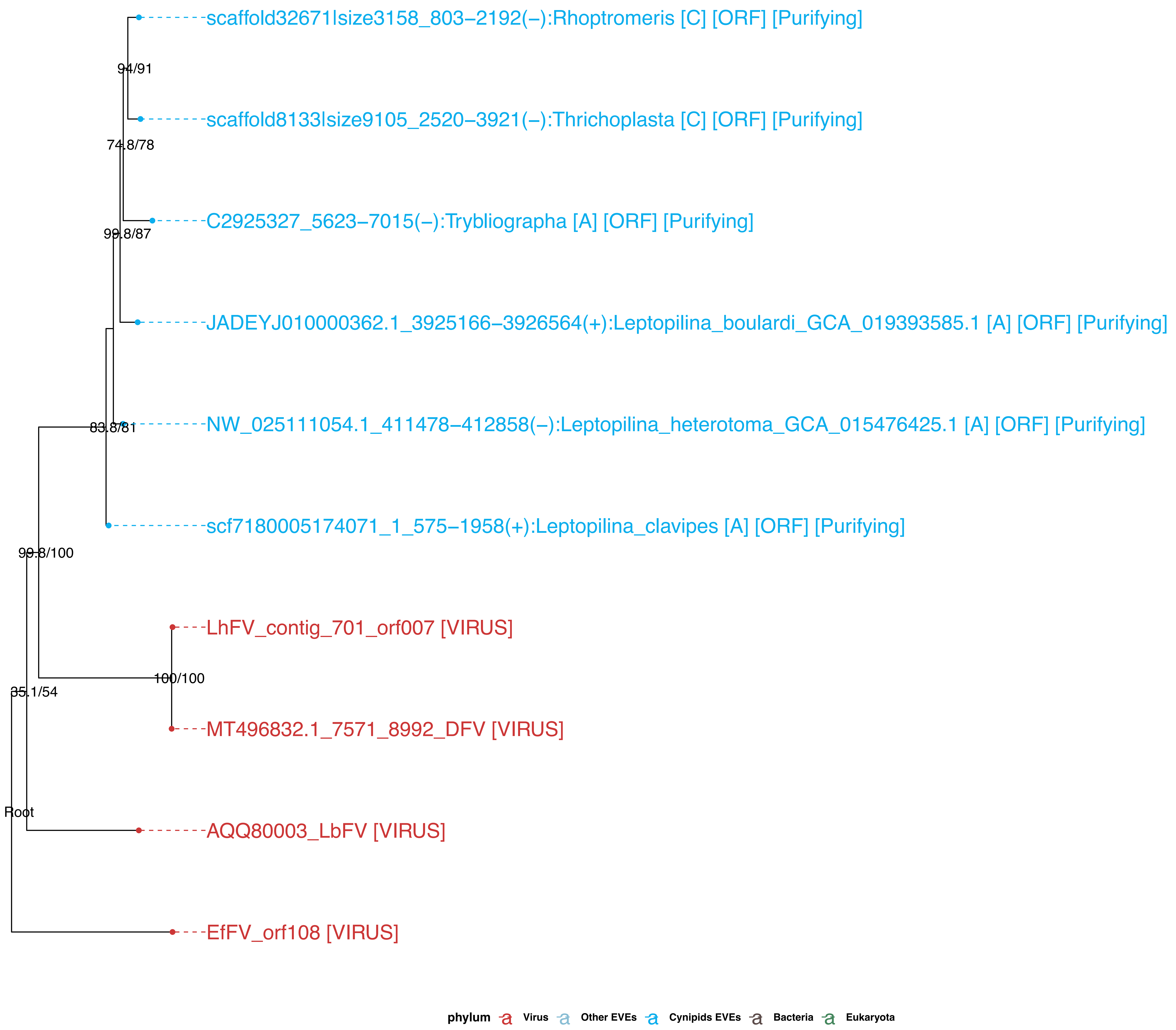

Figure S8-7: Cluster544 – LbFVorf87

**Phylogeny of type 1**

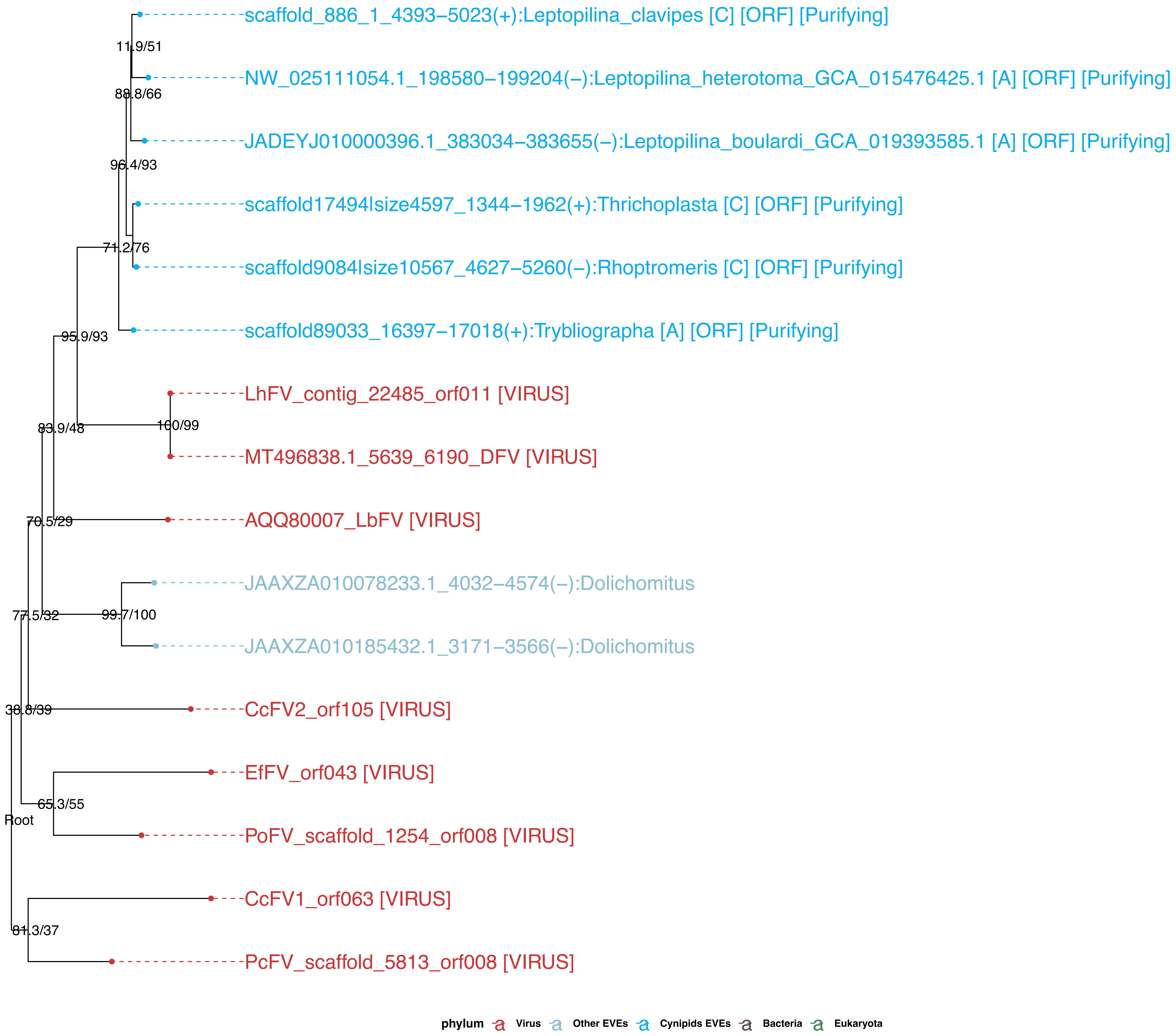

Figure S8-8: Cluster186 – LbFVorf94

**Phylogeny of type 1**

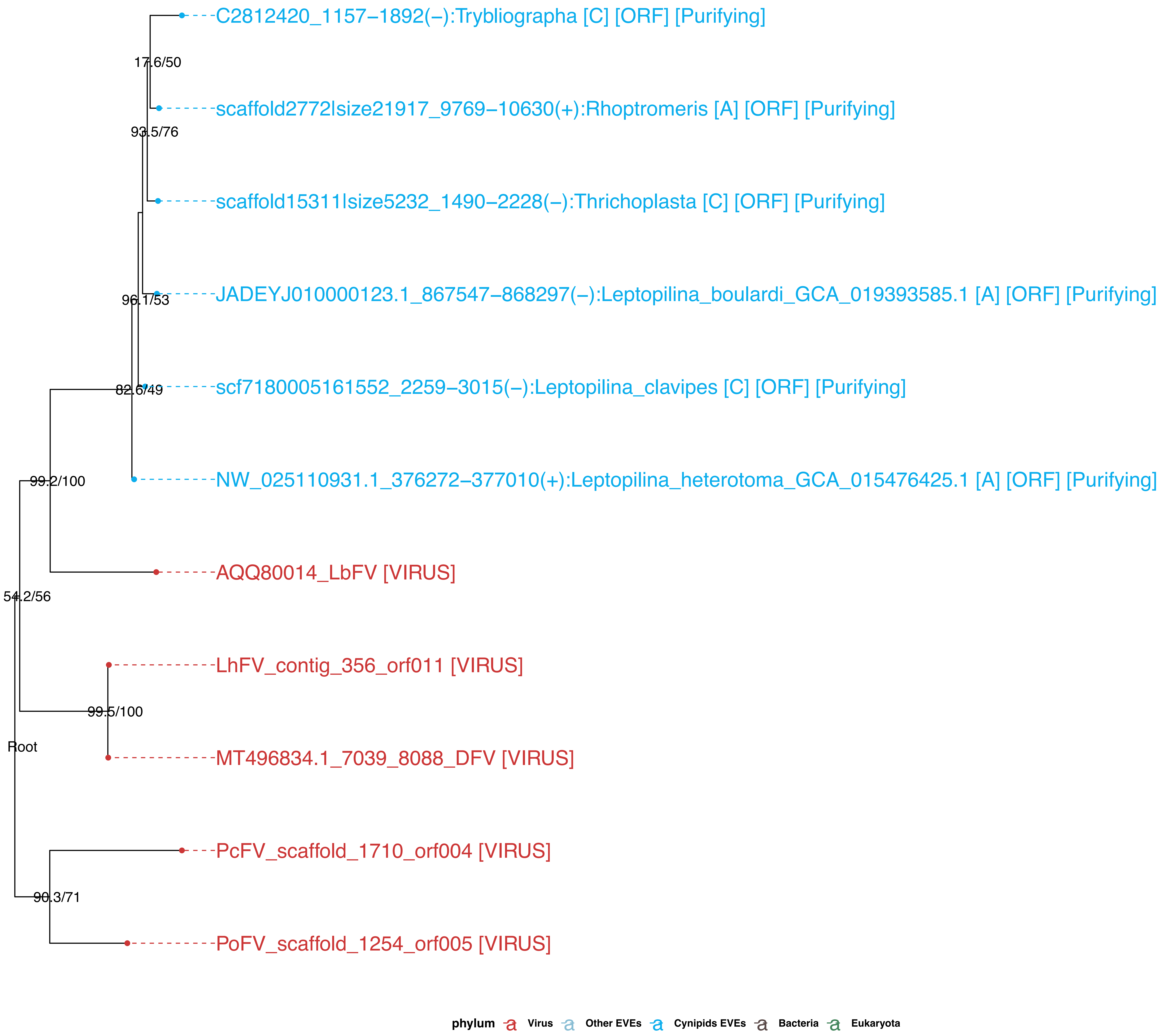

Figure S8-9: Cluster45_redefined – LbFVorf106 (Odv-e66)

**Phylogeny of type 2**

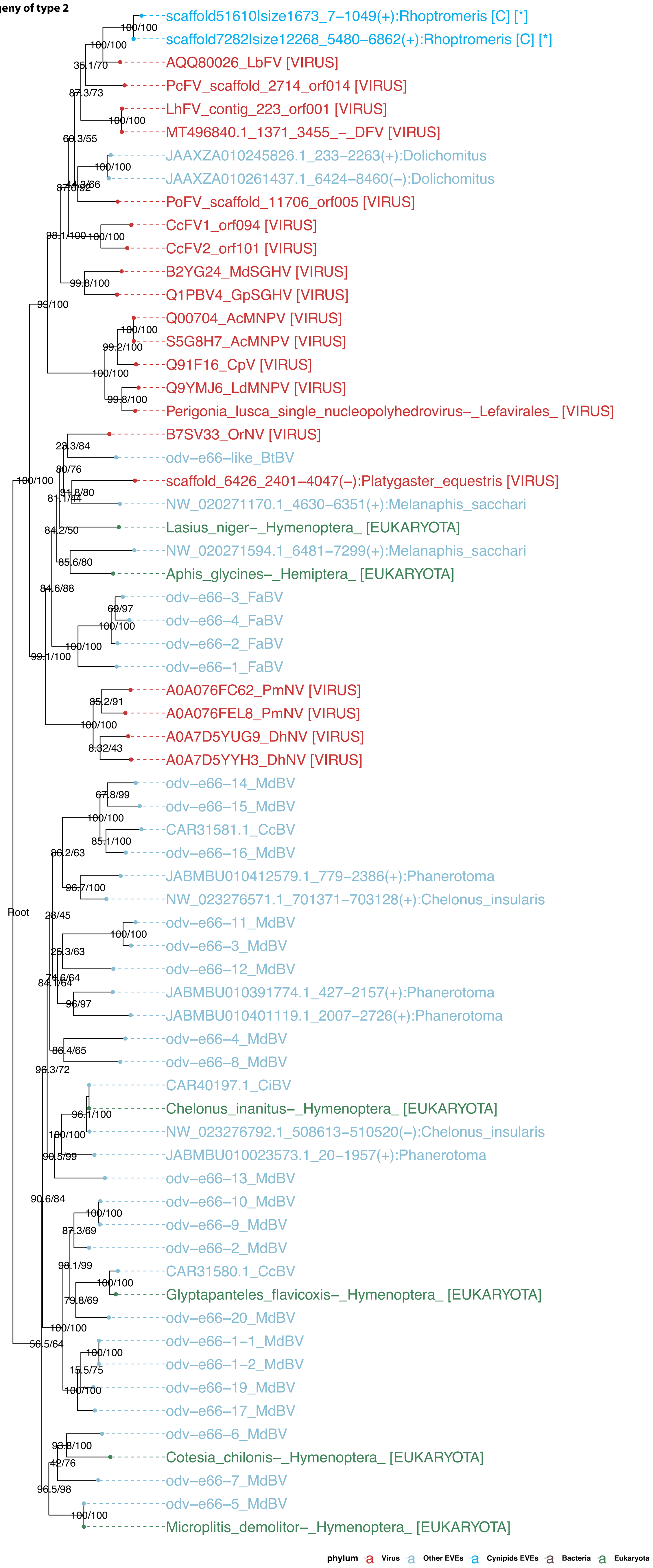

Figure S8-10: Cluster425–Cluster1270–Cluster1978 – LbFVorf108

**Phylogeny of type 3**

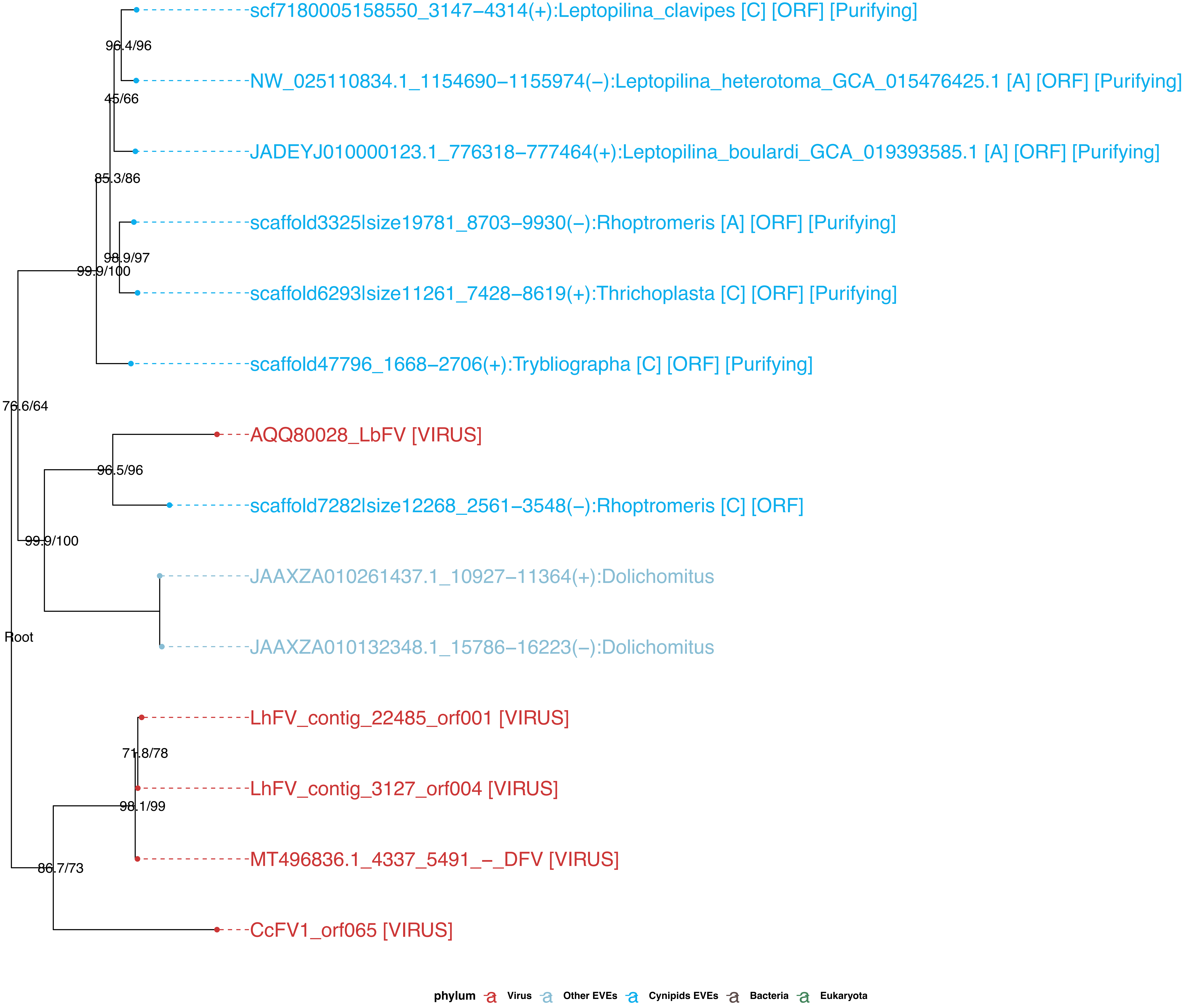

Figure S8-11: Cluster440 – LbFVorf60 (lcat)

**Phylogeny of type 1**

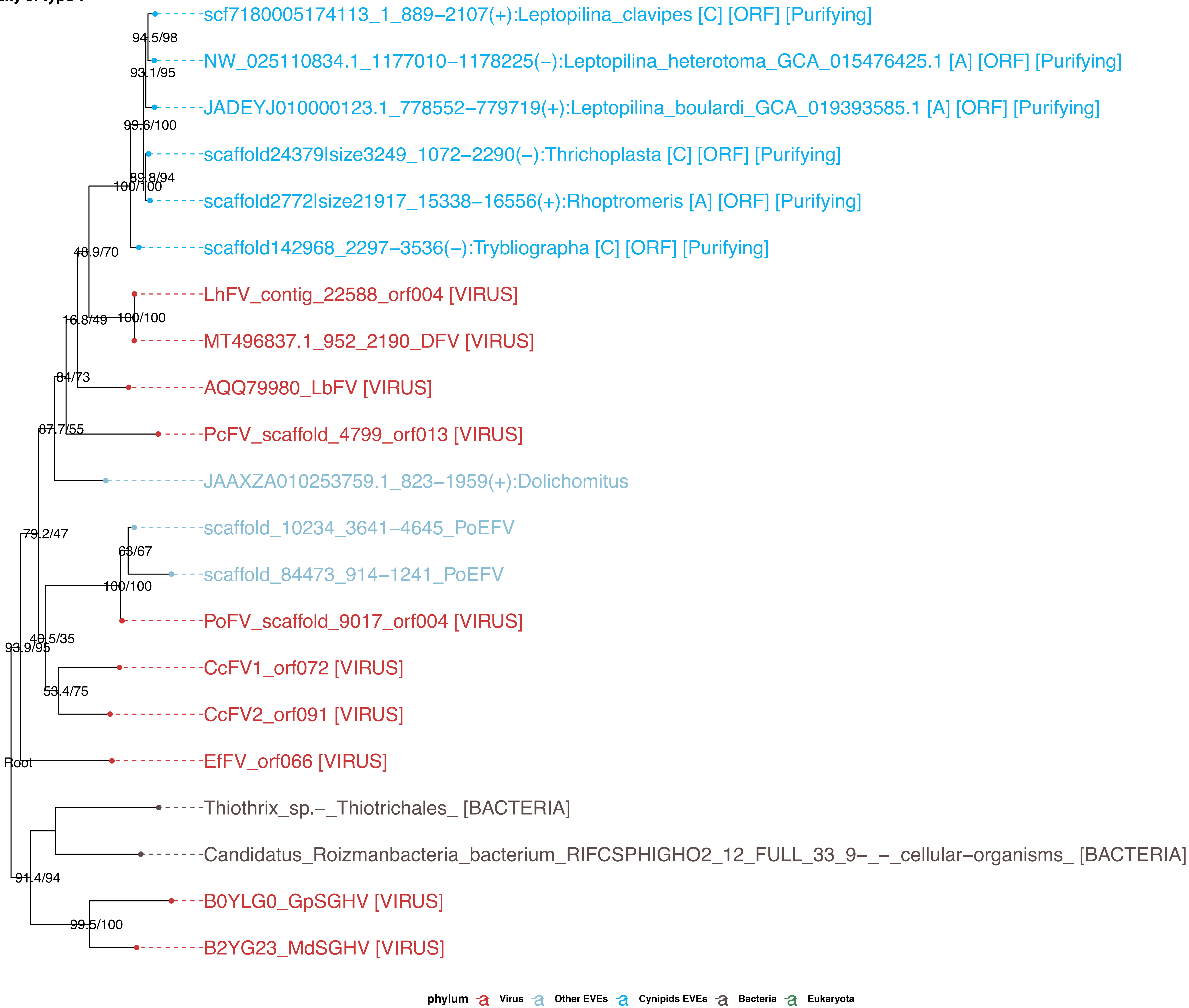

Figure S8-12: Cluster115–Cluster431–Cluster37 – LbFVorf107 (lef-4)

**Phylogeny of type 3**

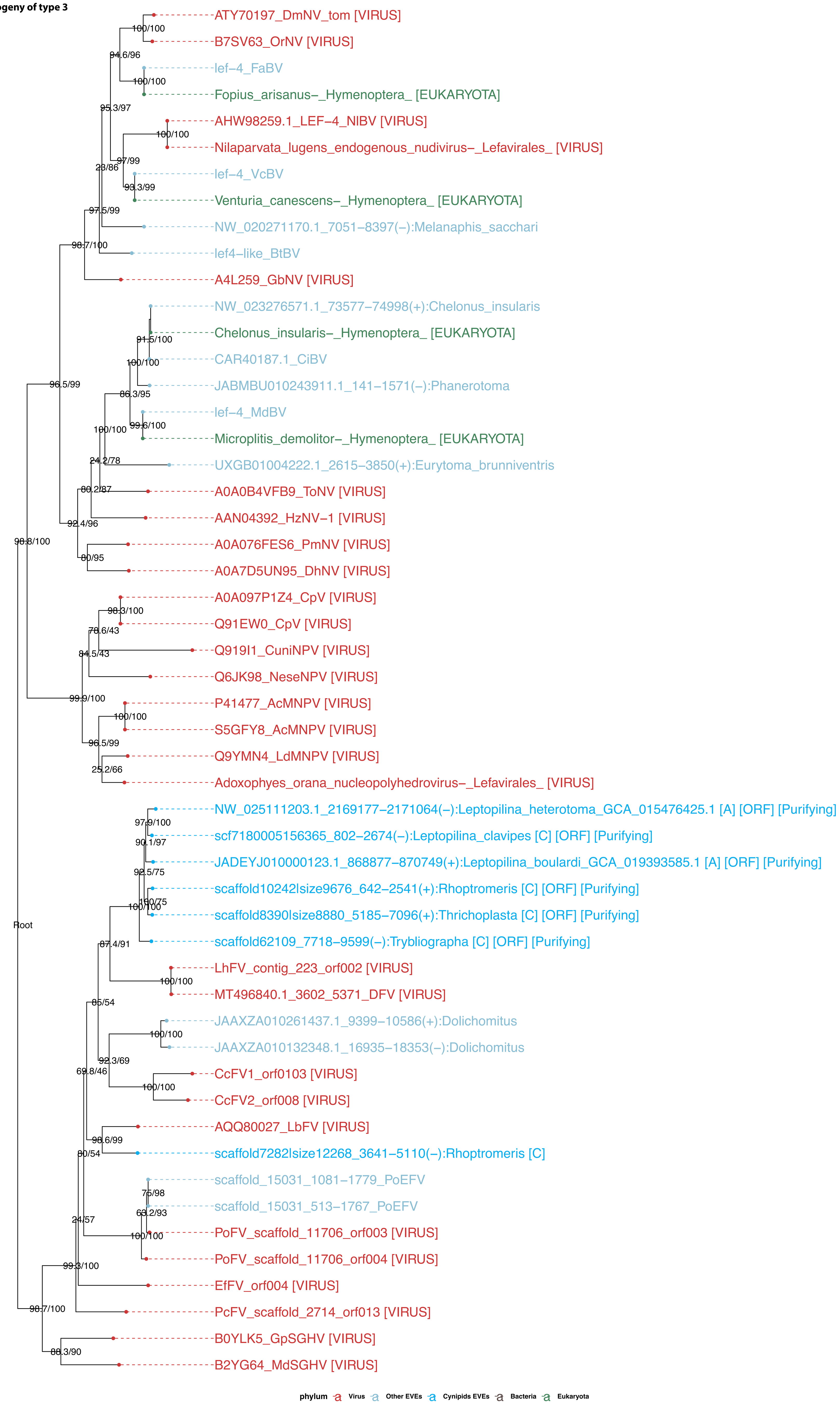

Figure S8-13 Lef–5

**Phylogeny of type 1**

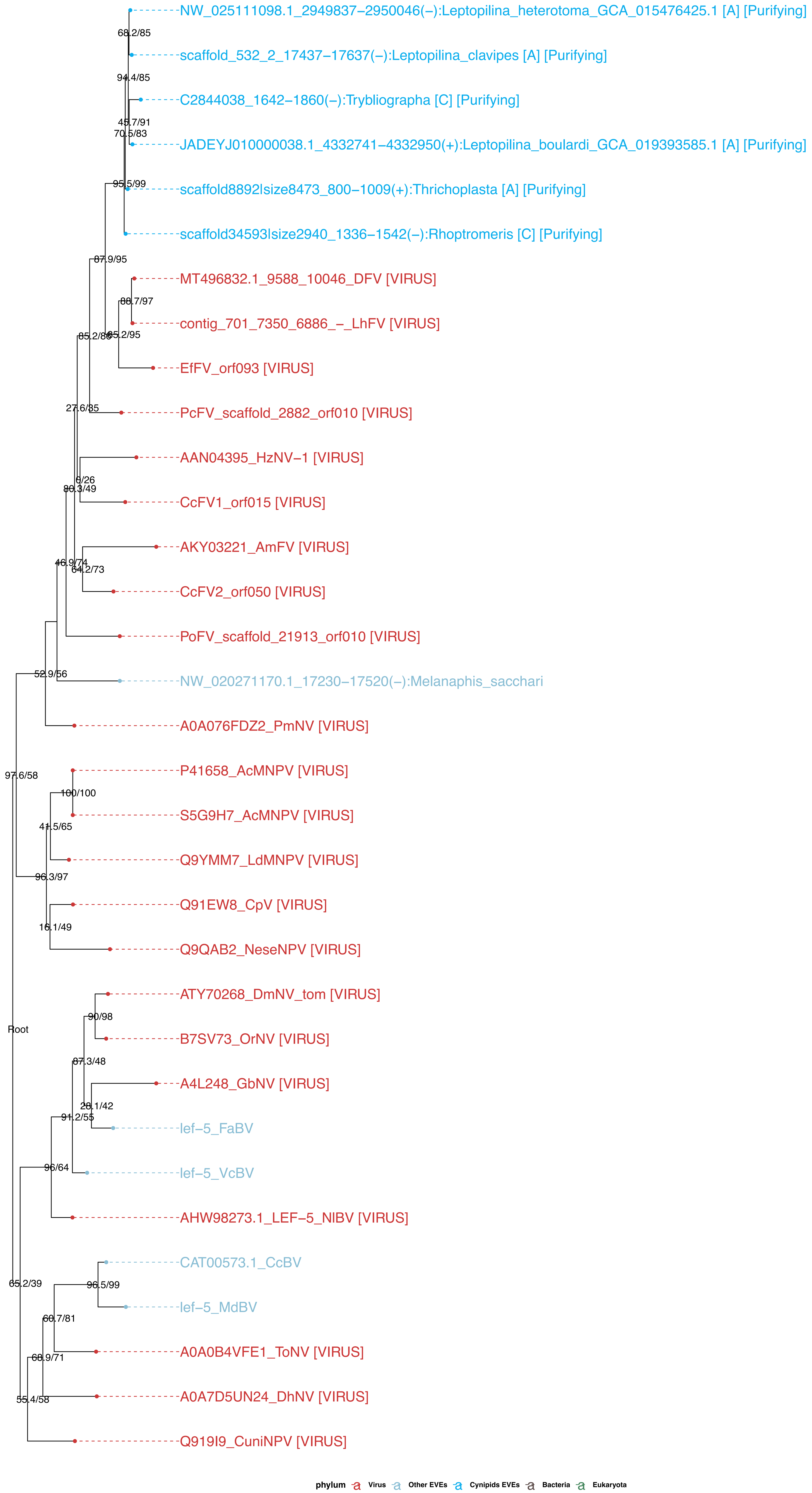

**Figure S8-14: Cluster1614–Cluster91_redefined – LbFVorf96 (lef–8)**

**Phylogeny of type 1**

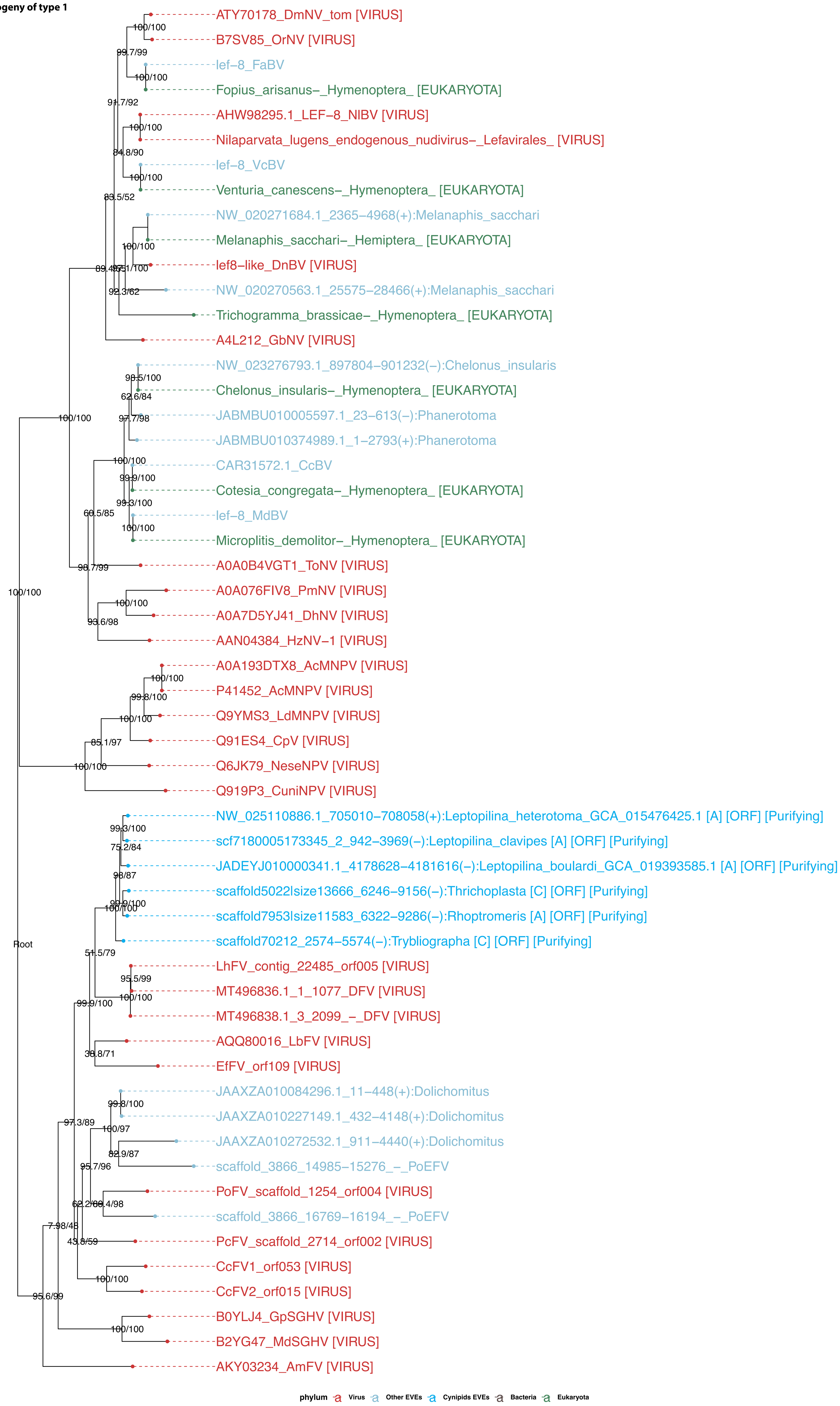

Figure S8-15: Cluster15 – LbFVorf78 (lef–9)

**Phylogeny of type 1**

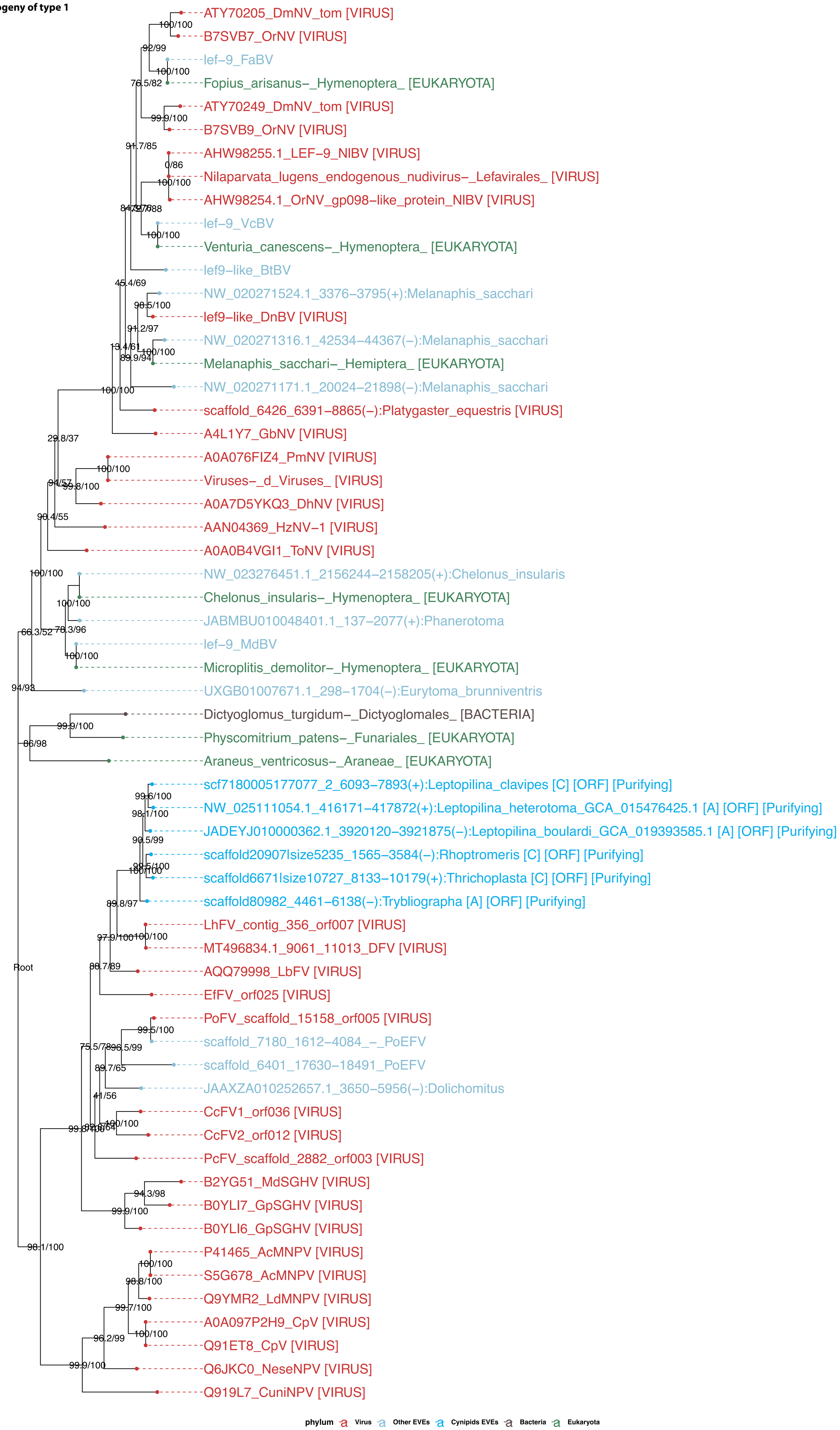

**Figure S8-16: Cluster126_redefined – LbFVorf58 (DNApol)**

**Phylogeny of type 1**

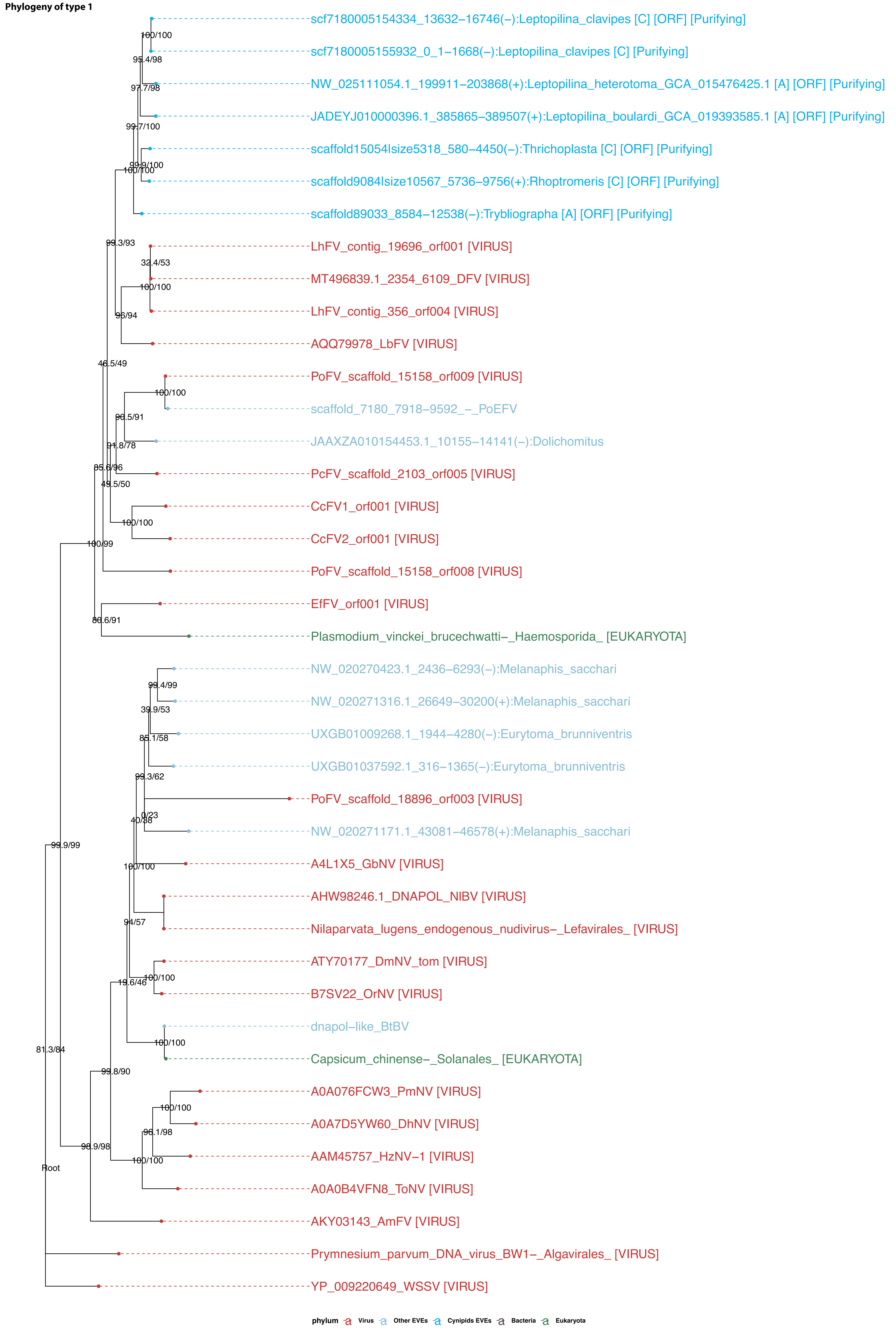

**Figure S8-17: Cluster22 – LbFVorf2 (Integrase)**

**Phylogeny of type 2**

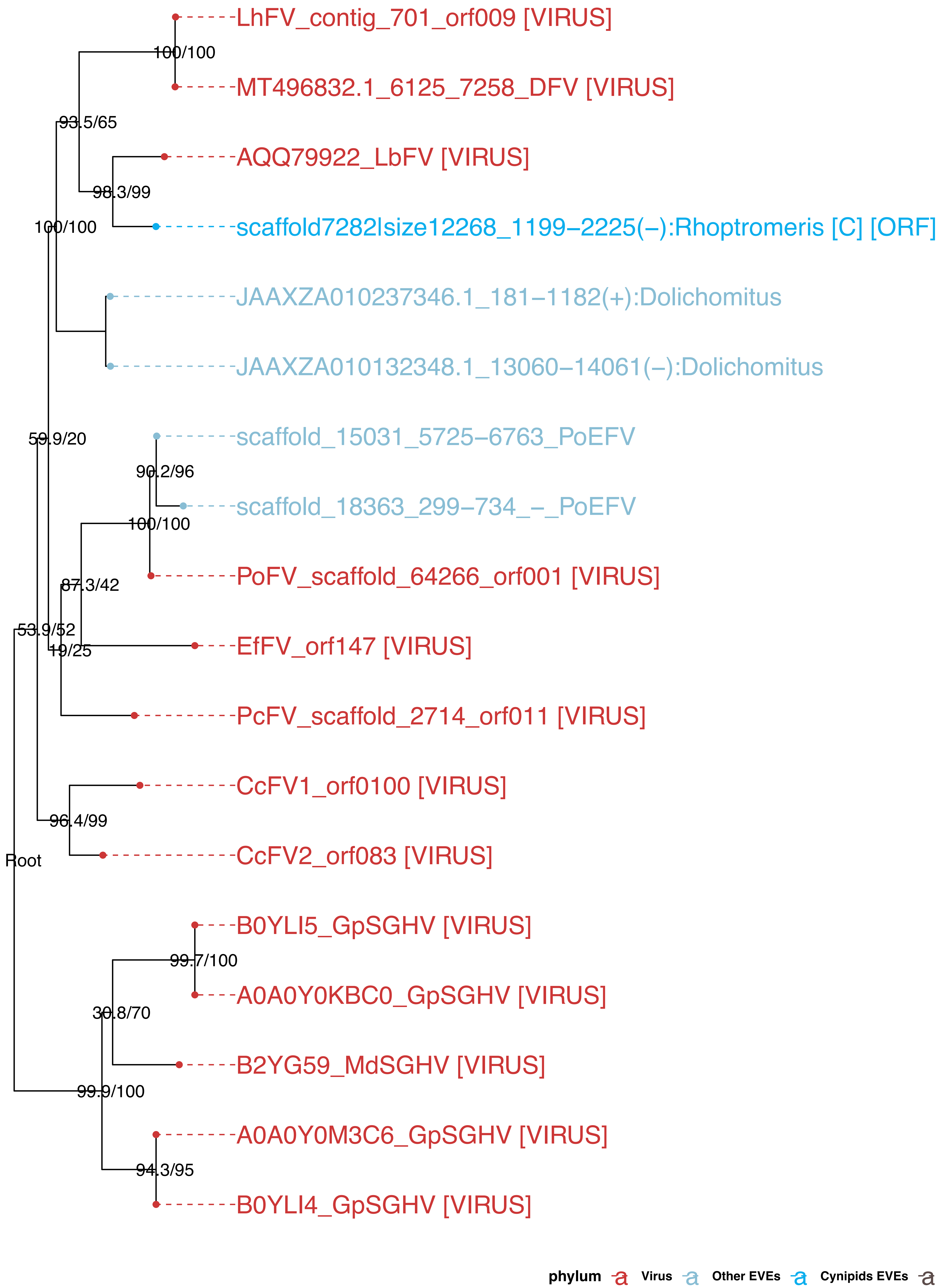

Figure S8-18: Cluster2683–Cluster470–Cluster79–Cluster2592 (Helicase2)

**Phylogeny of type 1**

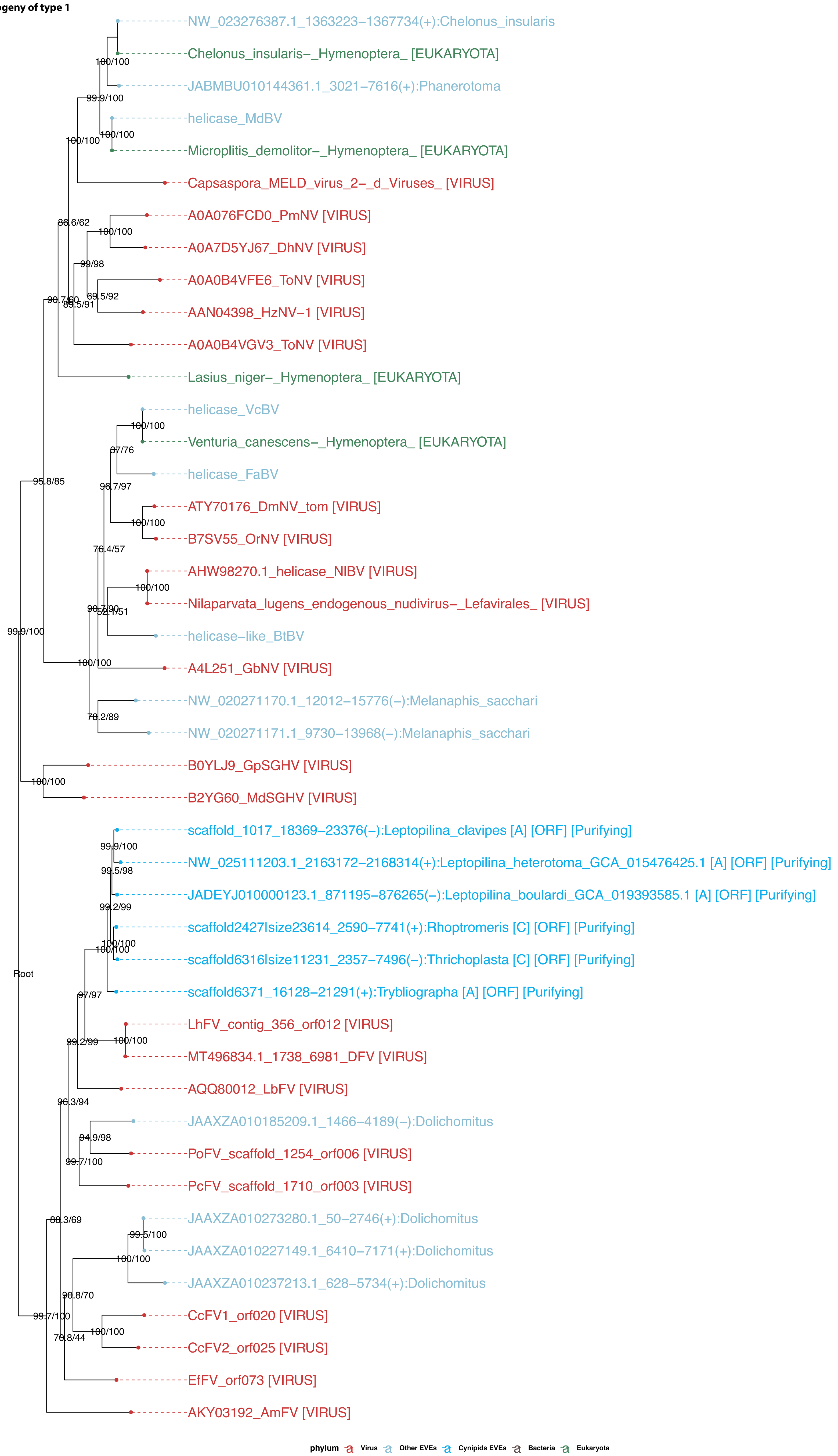

**Figure S8-19: Cluster59 – LbFVorf92 (Putative Lef-3)**

**Phylogeny of type 1**

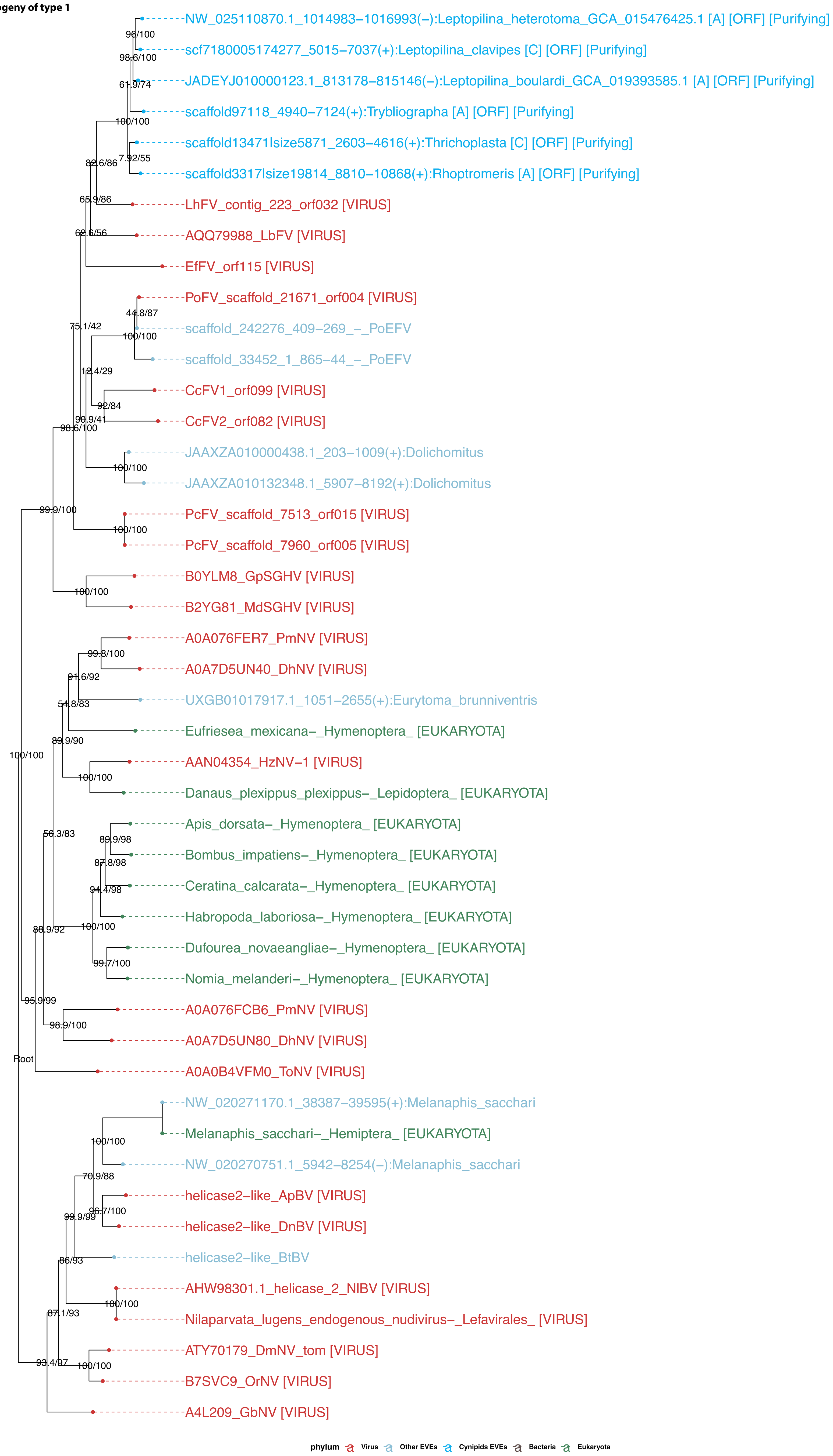

**Figure S8-20: Cluster38 – LbFVorf19 (38k)**

**Phylogeny of type 2**

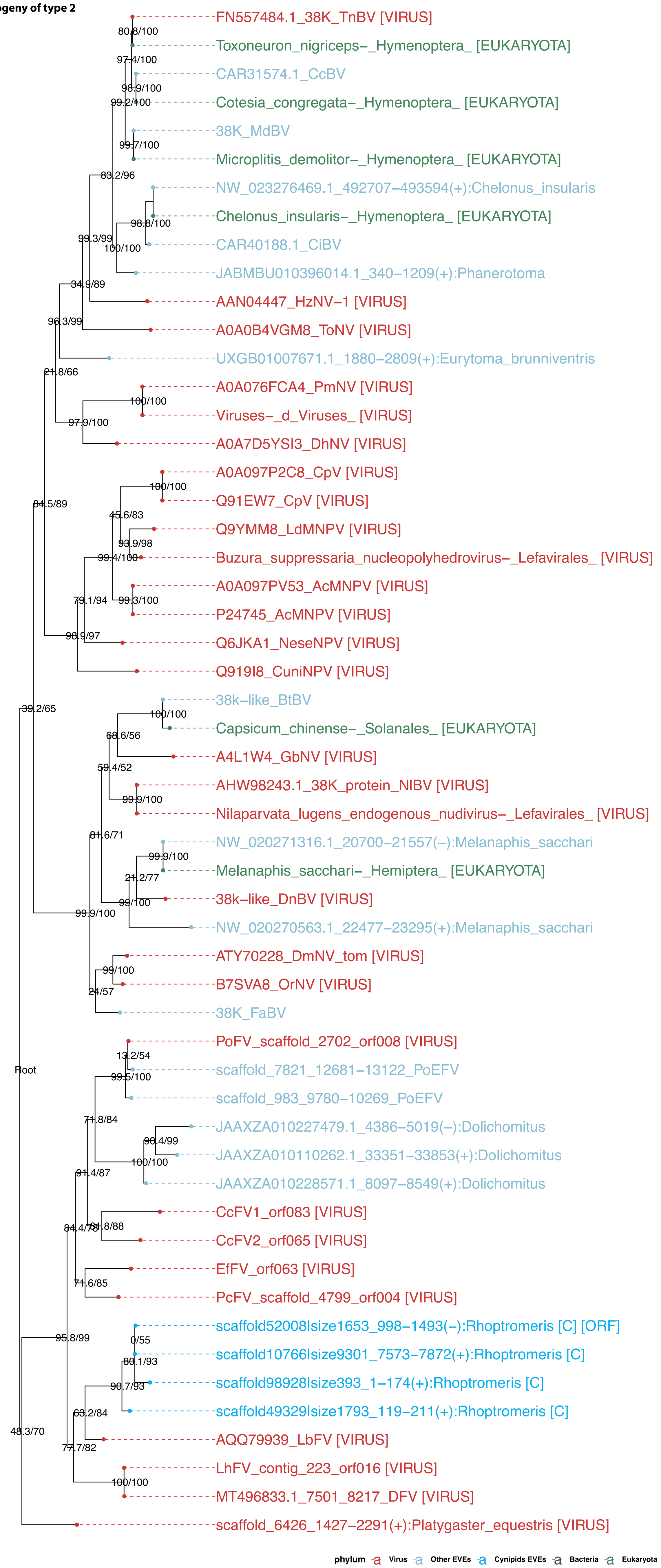

Figure S8-21: Cluster965–Cluster60–Cluster519–Cluster44 – LbFVorf85 (Ac81)

**Phylogeny of type 1**

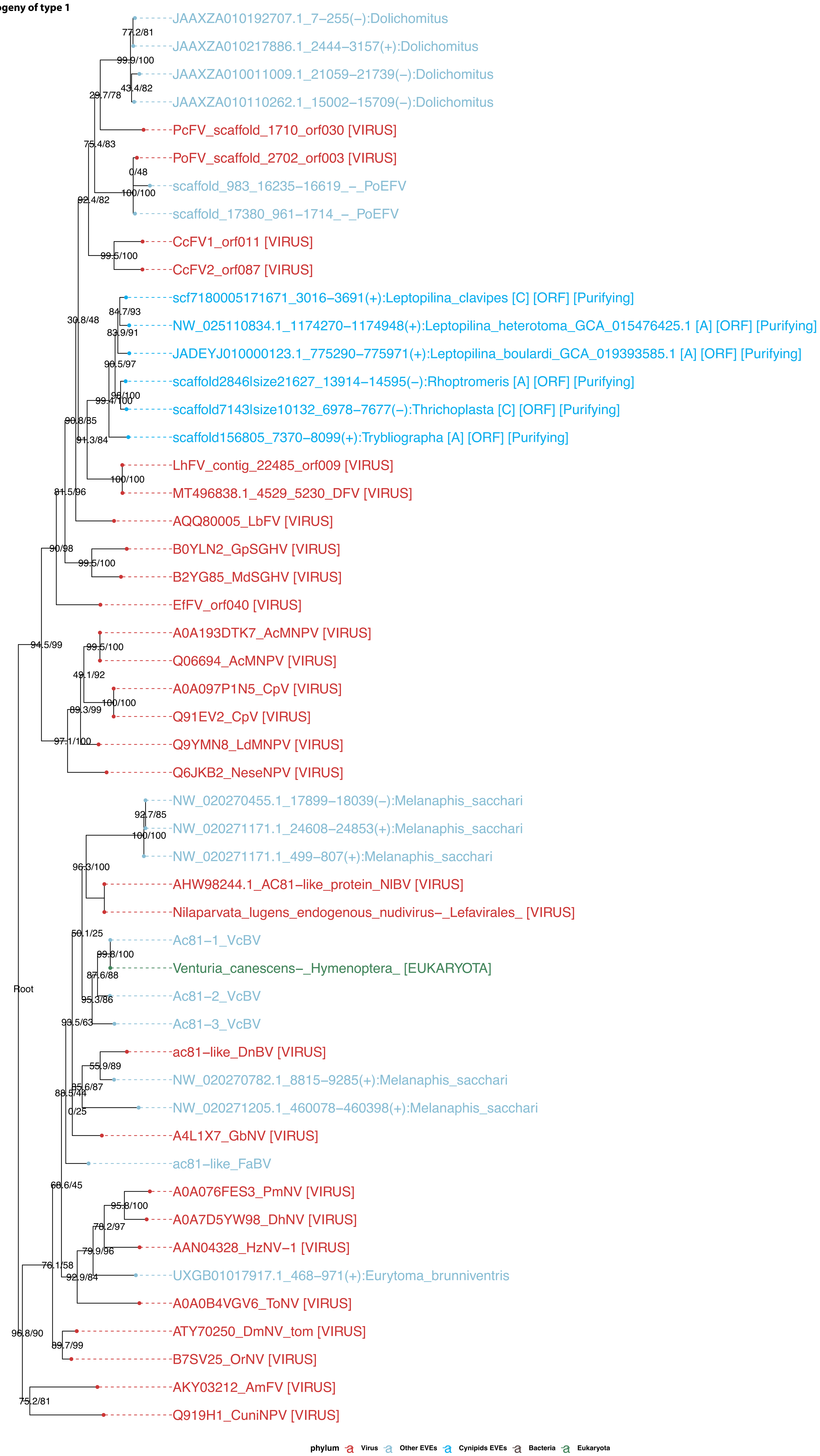
