## Supplementary material for "Dating the origin of a viral domestication event in parasitoid wasps attacking Diptera": FiguresS13: Cluster29_LbFVorf44_MSA.pdf

logo

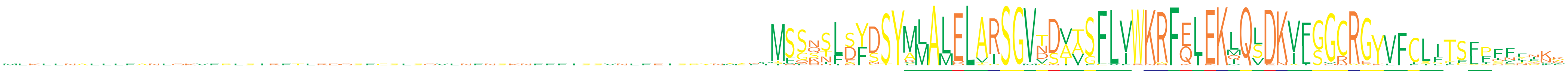

```
LbEFV1   ...........................................................................MMALELVRSGVTDAVSFLVRKRFELEKIQLDKVIGGCRGYVFCLFTSLFDNNPKK
LbEFV2   ................................................................MSSQSFSFDSYVMALELARSGVVDAAGFLVWKRFELEKLQSDKVLSGCRGYVFCLITSFPFFFFFF
LbEFV3   ...........................................................................MLALELARSGMTNVTSFLVWKRFELEKLQLDKVFGGCRGYVFCLFIPFFSCDNKR
LbEFV4   MLKLLNALLLFANLQKVFPLSIRFTLRDQSFCSLSQVLNFNSKNFFFISSVNLFEISPYNYSNIMSSNSLSFDSYMMALELVRSGVTDAVSFLVWKRFELEKIQSDKVLGGCRGYVFCLFTSLLILTQKN
LbEFV5   ................................................................MSGSTLEYDSYMLALELTRSGVNDTTSFLIWKHFQIEKQQVDKIFGGRRG................
LbEFV6   ................................................................MSGSTLDYDSYMLALDLARSGVNDTISFLIWKRFQIEKQQVDKIFGVNAGKLFILLPCLINKIYR.
LbEFV7   ..................................................................SQNFDYSSYALAMELAISGVDSVVSFLVWKRFQLEKLQLDKVFGGCRG................
LbEFV8   ................................................................MSSKSLSFDSYMMALELARSGVTDAASFLVWKRFELEKMQSDKVLSGCRGYVFCLITSFPFFL...
LbEFV9   ................................................................MSSNSLSFDSYMMALELVRSGVTDAVSFLVWKRFELEKIQSDKVLGGCRGYVFCLFTSLLILTQKN
LbEFV10  ..................................................................SHNLDYSSYALAMQLAISGVDSVVSFLFWKRFQLEKLQLDKVFGGCR.................
LbEFV11  ................................................................MSQNSFDYSSYSLAMELAISGVNSVTSFLVWKRFQLKKLQLEKAFGGCRG................
LbEFV12  ................................................................MSSNSLSFDSYMMALELVRSGVTDAVSFLVWKRFELEKIQLDKVLGGCRGYVFCLFTSLFDNNPKK
LbEFV13  ................................................................MSSQSLSFDSYMMALELARSGVVDVAGFLVWKRFELEKLQSDKVLSGCRGYVFCLITSFPFFFFL.
LbEFV14  ...........................................................................MMALELARSGVVDAAGFLVWKRFELEKLQSDKVLSGCRGYVFCLITSFPFFFIKC
LbEFV15  ................................................................MSSKSLSFDSYMMALELARSGVTDAASFLVWKRFELEKMQSDKVLSGCRGYVFCLITFFPFFYNNY
LbEFV16  ................................................................MSSKSLSFDSYMMALELARSGVTDAASFLVWKRFELEKMQSDKVLSGCRGYVFCLITFFPFFYNNY
LbEFV17  ................................................................MSSQSLSFDSYMMALELARSGVVDVAGFLVWKRFELEKLQSDKVLSG...................
LbEFV18  ................................................................MSSKSLNFDSYMMALELARSGVTDAASLLVWKRFELEKMQSDKVLSGCRGYVFCLITSFSPFFFII
LbEFV19  ................................................................MSSKSLSFDSYMMALELARSGVTDAASFLVWKRFELEKMQSDKVLSGCRGYVFCLITSFPFFYNNY
LbEFV20  ................................................................MSSQSLSFDSYKMALELARSGVVDVAGFLVWKRFELEKLQSDKVLSGCRGYVFCSITSFPFFL...
LbEFV21  ................................................................MSSNSLSFDTYMMALELARSGVTDAVSF......................................
LbEFV22  ................................................................MSGSILDYNSYMLALELARSGVNDTTSFLIWKRFQIEKLQVDKIFGGRRG................
LbEFV23  ............................................................MYFLTMFHNLDYSSYALAMELAISGVDSVVSFLVWKRFQLEKLQLDKVFGGCRG................
LbEFV24  ................................................................MSGSILDYNSYMLALELARSGVNDTTSFLIWKRFQIEKLQVDKIFGGRRG................
LbEFV25  ................................................................ISGSTLDYDSYMLALELAPSGVNDTTSFVIWKRFQIEKQQVDKIFGGRRG................
LbEFV26  ................................................................MFQHTFDYSSYALAMKLARSGVDSITFFLIWNWFQLEKLQLDNIFGDCRA................
LbEFV27  ..................................................................SQNFDYSSYALAMELAISGVDSVVSFLVWKRFQLEKLQLDKVFGGCRG................
LbEFV28  ................................................................MSQESFDYCSYLLAMELAISGVNSVTSFLVWKRFKLEKVQLDKVFGGCRG................
LbEFV29  ................................................................MSGSTLEYDSYMLALELARSGVNDTTSFLIWKRFQIEKQQVDKIFGGRRG................
LbEFV30  ................................................................MFSQSLSFDSYKMALELARSGVVDVAGFLVWKRFELEKLQSDKVLSGCRGYVFCSITSFPFFFYKM
LbEFV31  ..................................................................SQNFDYSSYALAMELAISGVDSVVSFLVWKRFQLEKLQLDKVFGGCRG................
LbEFV32  ................................................................MFSNSLAYDSYMLALELARSGVADVTSFLVWKRFELEKLQLDKVFGGCREYVFCLSTSLFCVIIKK
LbEFV33  ................................................................MFSNSLAYDSYMLALELARSGVADVTSFLVWKRFELEKLQLDKVFGGCREYVFCLSTSLFCVIIKK
LbEFV34  ...............................................................MSHNNNLDYSSYALAMELAISGVDSVVSFLVWKRFQLEKLQLDKVFGGCRG................
LbEFV35  ................................................................MSGSTLEYDSYMLALELARSGVNDTTSCLIWKRFQIEKQQVDKIFGGRRG................
LbEFV36  ................................................................MSSKSLSFDSYMMALELARSGVTDAASFLVWKRFELEKMQLDKVLSGCRGYVFCLITSFPFFL...
LbFV     ................................................................MSSNHISYDSYMLALELSRSGVTDVTSFLIWKRFELEKIQLDNIFGGCRGYVFFIIFSYIFN....
```

logo

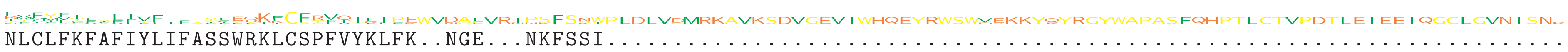

```
LbEFV1   NLCLFKFAFIYLIFASSWRKLCSPFVYKLFK..NGE...NKFSSI........................................................................
LbEFV2   FFFFFFFFFFFF...FFFYIKCFTFI...........................................................................................
LbEFV3   ICPYACLSVFVF.F.TIEFKL................................................................................................
LbEFV4   FYVCFNLLLF.....I.....................................................................................................
LbEFV5   .....................................................................................................................
LbEFV6   .....................................................................................................................
LbEFV7   .....................................................................................................................
LbEFV8   .....................................................................................................................
LbEFV9   F....................................................................................................................
LbEFV10  .....................................................................................................................
LbEFV11  .....................................................................................................................
LbEFV12  IVFMFI...............................................................................................................
LbEFV13  .....................................................................................................................
LbEFV14  FTFI.................................................................................................................
LbEFV15  KMFYVY..LIVF...YLEQKFCFRVQILIPEWVDALVRIPSFSNWPLDLVDMRKAVKSDVGEVIWHQEYRWSWVEKKYQYRGYWAPASFQHPTLCTVPDTLEIEEIQGCLGVNISNK
LbEFV16  KMFYVY..LIVF...YLEQKFCFRVQILIPEWVDALVRIPSFSNWPLDLVDMRKAVKSDVGEVIWHQEYRWSWVEKKYQYRGYWAPASFQHPTLCTVPDTLEIEEIQGCLGVNISN.
LbEFV17  .....................................................................................................................
LbEFV18  ITI.................KCFTFI...........................................................................................
LbEFV19  KMFYVY..LIVF...YLEQKFCFRVQILIPEWVDALVRIPSFSNWPLDLVDMRKAVKSDVGEVIWHQEYRWSWVEKKYQYRGYWAPASFQHPTLCTVPDTLEIEEIQGCLGVNISN.
LbEFV20  .....................................................................................................................
LbEFV21  .....................................................................................................................
LbEFV22  .....................................................................................................................
LbEFV23  .....................................................................................................................
LbEFV24  .....................................................................................................................
LbEFV25  .....................................................................................................................
LbEFV26  .....................................................................................................................
LbEFV27  .....................................................................................................................
LbEFV28  .....................................................................................................................
LbEFV29  .....................................................................................................................
LbEFV30  FYVYLIVILL......KTRKICFRVQIFIPEWVDAPVRNPSFSKWPLDLVGMRKAVKSDVGEVIWHQEYRWSWDAKKYRYRGYWAPASFQHPTLCTVPDTLEIEEIQGCLGVNISN.
LbEFV31  .....................................................................................................................
LbEFV32  YVLMLI...............................................................................................................
LbEFV33  YVLMLI...............................................................................................................
LbEFV34  .....................................................................................................................
LbEFV35  .....................................................................................................................
LbEFV36  .....................................................................................................................
LbFV     .....................................................................................................................
```

X acidic (−)
X basic (+)
X polar uncharged
X hydrophobic nonpolar
