## Supplementary material for "Dating the origin of a viral domestication event in parasitoid wasps attacking Diptera": FiguresS13: Cluster59_LbFVorf68_[helicase2]_MSA.pdf

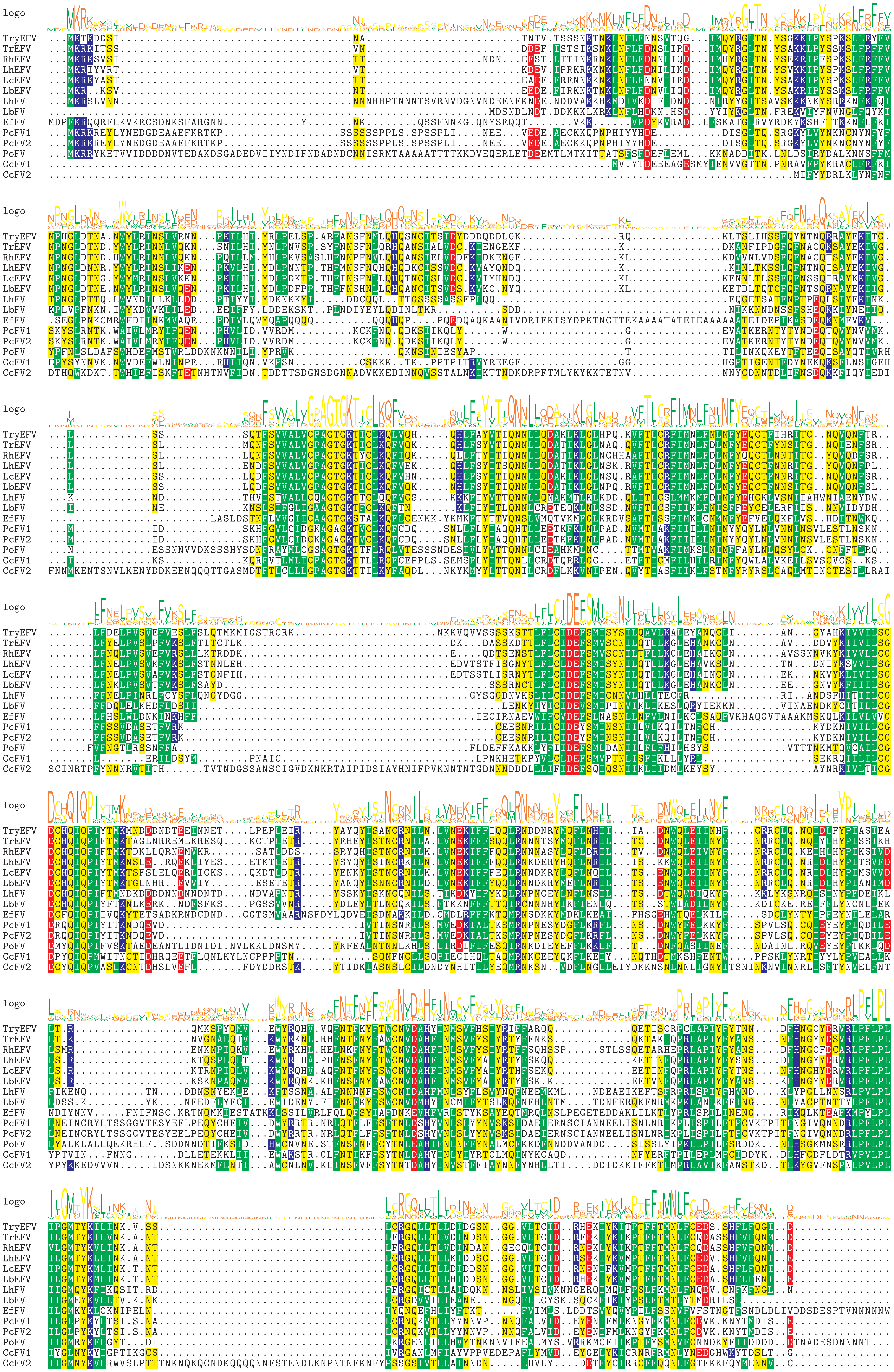
logo
TryEFV
TrEFV
RhEFV
LhEFV
LcEFV
LbEFV
LhFV
LbFV
EfFV
PcFV1
PcFV2
PoFV
CcFV1
CcFV2

logo 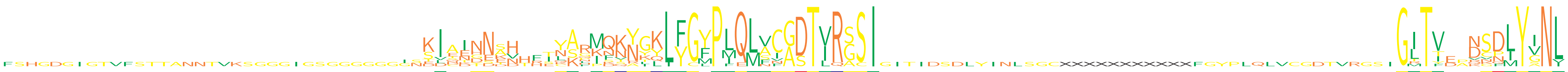

```
TryEFV  ...................................KIATNSTH...YAHMQKYGKLFGYPLQLVCGDTVRGSIGITIDSDLYINLSGCXXXXXXXXXXXFGYPLQLVCGDTVRGSIGITI..DSDLYINL
TrEFV   ...................................KITINNSN...YARMQKYGKLFGYPMQMACGDTVRASI...........................................GITV..DSDLYINL
RhEFV   ...................................KIAINNAH...CARMQKYGKLFGYPLQMVCGDTVRASI...........................................GITV..DGDMYVNL
LhEFV   ...................................KIAINNEH...YARMQKYGKLFGYPMQLVCGDTVRGSI...........................................GITV..DSDLYINL
LcEFV   ...................................KIAIDNAH...YARMQKYGKLFGYPIQLVCGDTVRGSI...........................................GITV..DSDLYVNL
LbEFV   ...................................KIAITNAH...YARMQKYGKLFGYPMQLVCGDTIRGSI...........................................GITV..VSDLYVNL
LhFV    ...................................ELCNGNEN..EKKFIQQRGQLYGFPLQLSFADTIRGSI...........................................GITY..NSDLYINL
LbFV    .....................................SKYNK....NKQKFVNTKIFGFPIQLNCADTIRSSI...........................................GMTVE.RKDIYINL
EfFV    FSHGDGIGTVFSTTANNTVKSGGGIGSGGGGGGGSSDDDDNDDTRTNKLKFKNIQLYGYPFELANASTVRSCI...........................................GLTFEPNVNLYVNL
PcFV1   ...................................SIEEHESVHFILSGINNNYYLYGMPLQMFCASTLRSSI...........................................GLTT..NCPLYGNL
PcFV2   ...................................SIEEHESVHFILSGINNNYYLYGMPLQMFCASTLRSSI...........................................GLTT..NCPLYGNL
PoFV    .................................INDINNNDK..HSNSNKKNNKLLYCLPLQLAVADTIRGSI...........................................GLTV..NSFIYGNL
CcFV1   ...................................TVQEG..........STAKLLFGFPLQLAIGDTIRSSI...........................................GTTVDAKSQIYSNV
CcFV2   .............................................TQAPPDSNLYLYGFPLQLSIGDTVQSSI...........................................GRTI..QNNMYANI
```

logo 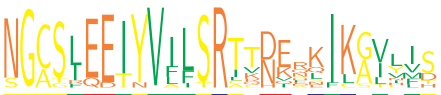

```
TryEFV  SGCSLEEIYVILSRTRDERKIKAVYIS
TrEFV   NGCSLEEIYVILSRTTDEQKIKAVYIS
RhEFV   NGCSLEEIYVILSRTTNQHKIKAVHIS
LhEFV   SGCSLEEIYVILSRTTDERKIKAVLVS
LcEFV   NGCSLEEIYVILSRTTDERKIKAVLIS
LbEFV   NGCSLEEIYVILSRTTDERKIKAVLMS
LhFV    NGCSIEETYVILSRTKHVEKIKSVCL.
LbFV    NGCSLQETYVILSRATNIDQIKGIYIH
EfFV    NGCGREDTNVAISRIKNKSNIKGLVI.
PcFV1   NGASIEEIYVFLSRTVDENLIKGILVD
PcFV2   NGASIEEIYVFLSRTVDENLIKGILVD
PoFV    NGCTIQEIYVFFTRICNKQQILGINF.
CcFV1   NGASLEEIYVLFSRCQNKASFCGTVMT
CcFV2   NGASFEEIYVLFSRLTNEFEILGLIF.
```

- X acidic (−)
- X basic (+)
- X polar uncharged
- X hydrophobic nonpolar
