## Supplementary material for "Dating the origin of a viral domestication event in parasitoid wasps attacking Diptera": FiguresS13: Cluster114_LbFVorf11_[JmJC]_MSA.pdf

logo



```
LbEFV   ..................................................................................................................................
LhEFV   ..................................................................................................................................
LcEFV   ..................................................................................................................................
RhEFV1  ..................................................................................................................................
RhEFV2  ..................................................................................................................................
TryEFV  ..................................................................................................................................
LhFV    ..................................................................................................................................
LbFV1   ..................................................................................................................................
LbFV2   ..................................................................................................................................
EfFV1   ..................................................................................................................................
EfFV2   ..................................................................................................................................
CcFV21  MPINSSQKVTWVPEMNSPHFSSYSPIQLKNYFTGLMSPSLIEECKYSKNTYQNQQHEEITITSNTVISNTTIDFKKLLELENDPVIDPASTLSFSCDFLISEKAVVKTSKRYERRHDYDDDDEEDKNFQR
CcFV22  ..................................................................................................................................
CcFV11  ..................................................................................................................................
CcFV12  ..................................................................................................................................
PoFV    ..................................................................................................................................
PcFV    ..................................................................................................................................
```

logo



```
LbEFV   ..................................................................................................................................
LhEFV   ..................................................................................................................................
LcEFV   ..................................................................................................................................
RhEFV1  ..................................................................................................................................
RhEFV2  ..................................................................................................................................
TryEFV  ..................................................................................................................................
LhFV    ..................................................................................................................................
LbFV1   ..................................................................................................................................
LbFV2   ........................................................................................MQNEEILMENVSSSSFMKEK......................
EfFV1   ..................................................................................................................................
EfFV2   ..................................................................................................................................
CcFV21  LTEYLQRELENVDFDEIPQQHSVENEELIDKQSQEHHEDISDILHVQNNDELSQQKHTIEHMEIEGEIEEQYENLNDVLFTTDQNNNDMRQQQQYDEAFSNNSWNIEDTEIIEVTAQ.......ERITSL
CcFV22  ..................................................................................................................................
CcFV11  ..................................................................................................................................
CcFV12  ..................................................................................................................................
PoFV    ........................................................................................MNVDQSFLSDEATNTYPSNDVPSLNLLSPFSPQWLQQNHHHQ
PcFV    ..................................................................................................................................
```

logo



```
LbEFV   ..................................................................................................................................
LhEFV   ..................................................................................................................................
LcEFV   ..................................................................................................................................
RhEFV1  ..................................................................................................................................
RhEFV2  ..................................................................................................................................
TryEFV  ..................................................................................................................................
LhFV    ..................................................................................................................................
LbFV1   ..................................................................................................................................
LbFV2   ENTVNQKIDTNISTPMS.EVLSIMHNTSM...........YLMN.......TDIGNYSNIEII............PSS.....MELDISSNGTNVT........VIPPPSIIDDYH...NGLLNTELSY.
EfFV1   .........................................................................................................MAVASYLNNYVVSPSDY........
EfFV2   ..................................................................................................................................
CcFV21  EEQDQQRINTSIEGQDQQQINTTLHEQDH...........QQIN.......TSIEGQDQQRINTSIEGQDQQQINTTLHEQDHQQINTSIEGQDHQQIITLTEEQIPQLIILNQEQVPEHQQINTSIEE.
CcFV22  ..................................................................................................................................
CcFV11  .................................................................................................MNTDRELSMAIASVVGDISTNDNGA........
CcFV12  ..................................................................................................................................
PoFV    QQQQQQQQQVLLYSPMPQSVSSTLNTPQPDFSEISENQYKYQHHHQHQHEHHHHHHQTQQELQNQ.......YCASPLHQQQQQQ..........QQQQQQQQQQIPEAIYMTPNIEPTY.LFSPSFS..
PcFV    .........................................................................................................................MTTTCFDIY
```

logo

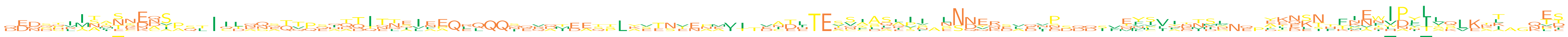

```
LbEFV   ........MSRENS........................................................SF......................LS.........DCEEN.YIEWIDYLLKLKTK...ES
LhEFV   ......MSSNSECS........................................................SYLSLPI.LS......P......ENV.........EEDST.KVEWIDYLLPLKAK..QES
LcEFV   ..................................................................................................................................
RhEFV1  ................................................................................................................KTSIPLT.....I...TD
RhEFV2  ...................................................................................................................IPLT.....I...TD
TryEFV  ......MSTSCKDS........................................................SSSLKSS.SS......S......TYS.........AINST.TIEWIDYLLQLKNT..QET
LhFV    ................................................................................................................MEFWPFLAQ.......RS
LbFV1   ..............................................................................................MSLFKISL...EKE....KNNVAVIV....T...PR
LbFV2   .ENSHLIN.NNDNSPSTIIDKNSTTPSITTTTNIINP.....N.....................NTITESNIASQVN.LNDE.....N.....NFIVIDNIL...YKQKNFFINNIPP......L...TI
EfFV1   .DDDTIIAAEKEQA...ICTAGATT...TTITTAVAA................................AVAASAIP.ENDYR...IP.....EVL.YRMSG...AKNKI.......FIG........ET
EfFV2   ..................................................................................................................................
CcFV21  QEQQQIITLTEEQIPQLIILNQEQVPEHQQINTSIEEQEQQQI.....................ITLTEEQIPQLII.LNQEQ...VP.....QFIIQTNDS...FLQKN..EVRIPNIS....L...L.
CcFV22  ..................................................................................................................................
CcFV11  ......AAATYEAVA............AATIENE..........VAIEETPLKETNEEAAVI.EADHTES..AQEPV.PNNERSVENE..........NTSNNE........PEIHPTIVSVEKAAGNEE
CcFV12  ..................................................................................................................................
PoFV    ....PVTTINNNAS...IQLQQQQQQQQQQQQQQLEEQRQQQQQQQMEESTLAVENVNNEITTVAELTEVCCDKIII.NNNQKELQLPSQQSVEYCVA..KK...YENSNTFNAFV..............
PcFV    DDDGGRINNSNDDT..............DDITLEIFEQLLQQSERDTDAQSLYSTSTEPMVI.DCDETETVAAAATFANNNVNDAD.DDDDIFMYPVIETMVNQPTTKLSHFNNTLNILT...CT..KVS
```

logo

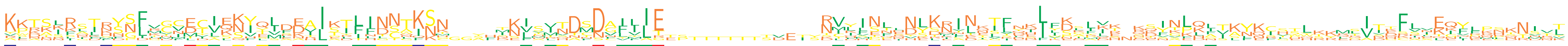

```
LbEFV   KPKSIRSTRSSLVCCETIQSIQLDEAISTFLNCTKSN....MNVSINDMDAFIIE.............TMYINLQDLKRINITLNELFDGIIN.REINLQIIKYKCSSL.....................
LhEFV   KPRSIRSTRSSLVCCETIQSIQLDEAITTFFNCTKNN....INVSMNDMDVFIIE.............RMYINLHDLRRTNATLDELFDAIVT.RQINLQAVKYKCTTL.....................
LcEFV   ..................................................................................................................................
RhEFV1  KKTSLRAIHYGFMGVDCIEKVYVFDVLKTLIDNTISK....TKISVTDSDGILLE.............RVLISTENYKNINEIFNKITSSLLR.QDYKLKF...........LVTEFSVEKFLPHKNI.T
RhEFV2  KKTSLRAIHYGFMGVDCIEKVYVFDVLKTLIDNTISK....NKISVTDSDGILLE.............RVLISTENYKNINEIFNKIISSLLR.QDYKLKF...........LVTEFSVEKFLPHKNI.T
TryEFV  KPRTIRSTRSSLVCCETIQSIQLDEAISIFFNCAKSN....MNIAMNDMDAFIIE.............SLYINFHDLKRINQTVDQLFEAMVT.GKINLQAVKYKCSTL.....................
LhFV    KEKKSSGTFSNIGMCMCASKYSIDDIIAYFINNLKNC....IEFNTTDCNALILE.............NVILGEKHYKIITKTLEHIVKILKK.NNSAFHNIKYEIVEQK..SIITKLLQQ.........
LbFV1   KLTKKINIKYGFLGIECVENHEIENIISLLINNTISN....TSVTVSNSDGILIE.............KVKVNLINSLKIEKSFQQIYNELVK.KSVTKNL...........YISKRVCKQFLQNSTDVY
LbFV2   YKNSKFLLNYSFKGVLVVKNFTLAKTITDIFNNTKSN....SKLVLHDCSAFFLE.............NIHFENNYKDLVLKVFQDIKKLFQS.DTFSLKLTSNKIHPLSRMDVTNCVIEQIEPSQRNVK
EfFV1   RSDNNRLLRYNFRVF..VEKYLMRA.......SAKNT..A........FDFYYIR.............DCFIS....DRFDSTFRILFKKAIK.SSNG................................
EfFV2   ....................................................MI..............RVKSNF...SNNLQGFLHFLDTYDENEKFVPD...........LGAIGFLAKGVFEC.....
CcFV21  KKAKRKSFNDNDTDNDDVKNY...KRIKTTIDNNKNM..YFKKLPIV.TDNLLLTEP...........RTLMEPLNESQALNIIKKIIPKPK..KQYNELITLNNFLHYRKETINQFLEENYRKRNNSNL
CcFV22  ......................MSEAIKILHGTGFSH.................G........SVEIYNVDLTNRNSKNLLYSSYILWKGFEE.KSITP................KFHTLEYNKNDNVRT
CcFV11  VWTRLFKTKHSFRNMTIYKKIEL...........................................................................................DTFINTIYKRSTNITL
CcFV12  ...............................................................................................................MSIVHFLNKGLPMP.....
PoFV    ..TASLIDSFNFKCNEYLLQYQS..HLGRIITSQKKN......IDITANDVVVATTSTTTTTTTDETSNIQHPLLNLTMCKNLTHNLYDLRKH....GKLLSAFQRGWHKKQRVKDFKIEQYFNLKYMYV
PcFV    AKDAKKQKMNPLHTIMCLSNYTILDALEILNLSWQKGGGSHDKLMRQTSNCIVIE.............NCIFNQSVLKEINTLKYSLLSDLI.....NPQSTALQSHSWTYRQITDSAHEIQ.....SQL
```

logo

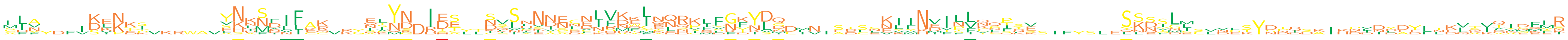

```
LbEFV   ..................YNKMDIF.....QIYN.IDE..NLININ..ENHNFLKQQKQFGKYLQ.............ILLQQTT........SSSSL..............................LN
LhEFV   ..................YNKMDIF.....QTYN.IDT..SVLNNIECENSMWMRQRKLFGKYLQ.............ILFKQIS........SSSSL..............................LR
LcEFV   ..................................................................................................................................
RhEFV1  LLN....SLK................R...ESND.IES..DVLLNNECNIVDKIN..................EIINV.NV.................................................
RhEFV2  LLN....SLK................R...ESND.IES..DVSLNNECNIVDKIN..................EIINV.NVCCP.........SLQIDS.........................A.....
TryEFV  ..................YNKMDIF.....ELYN.IDT..CVLNNAGNETFDWCRQRKRFGTYLQ.............TFLQESSS.......SSSSL..............................LR
LhFV    ..................VKQSRLF.....NEYN.DEE..SSSP.........LQNQSIDNDYVGDIN.SESLKLINEILLKSILK.......LNESMM.........................QKMKMR
LbFV1   LLY....KEKKK......LNNNFLIAK...NLNN.IS...SQSFKDELLLEKQIN..................NLMNV.NLNPT.........QKDILI.........................K.....
LbFV2   MIF....KENES......SSLS..LC..............NNSANDNDGIFKELA..................KIIND.T...F.........SKNVLI.........................Q.....
EfFV1   ..................................................RMELENDCMVGKYSDCLV.SRSEVVYKEFPVVLPKE.......SSDGNL..VGSYDDD....DDDDDD..DVTILYNFMR
EfFV2   ..................ENANVILEQVRGILYRDRDSYLSTNKEDNLNLVRELQLRPLSNYKDG.................................................................
CcFV21  GLV....KKNKS......YHHSFIFKC...HPY...EC..KKSHSNNRNLVKTLT.........M....ADCSKIFHIIPICNFNV.......SKDFGL.........................TIDEYR
CcFV22  CHV...IDIKDI......YKKSK........TYNDITE..NSVINISNDLFNDCQNHI.....DTDMNIETLNKYFNDILFNFQYESIFYSLEYKNYLMSLNEKYNPIDKIHKIIRDYLTEQQIDVGFFS
CcFV11  SFA....KKNYPVKRWAVENLNNVFAK...SVMG..................AFPDR.IFKFNDD........NRVNMVSLTQPRV..........TGECVDLTYSTQSAIKPVNLEELNKLSVV.SMEK
CcFV12  ..................VHNFF.....KYIKQSDR.SLTDGSVVTCSVLVAQIEQNPLLNFNLN.................................................................
PoFV    MIA...IDENKY......ANIS..........................DIVKFINESKFYFFYDI......................................SYDNR....NLDYAYFLKVKKQIDFSI
PcFV    AIAYDFVDVPTI......SKENINIAK...AIYD..HA..IISHNVAESKRKKTHLRVFFGYIDQ........................................................LAVTAEQ..
```

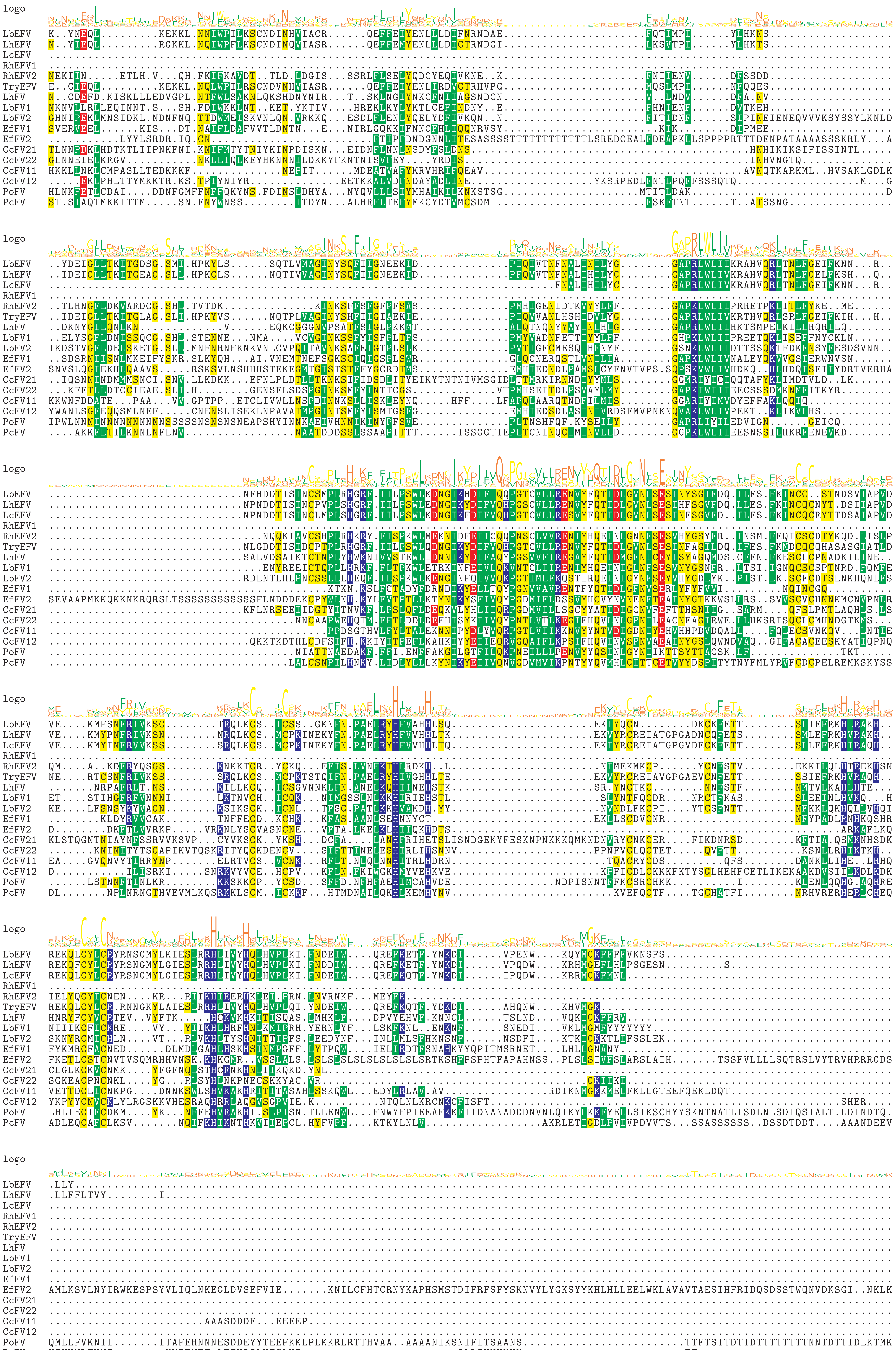
logo
LbEFV
LhEFV
LcEFV
RhEFV1
RhEFV2
TryEFV
LhFV
LbFV1
LbFV2
EfFV1
EfFV2
CcFV21
CcFV22
CcFV11
CcFV12
PoFV
PcFV

logo

```
LbEFV   ..................................................................................................................................
LhEFV   ..................................................................................................................................
LcEFV   ..................................................................................................................................
RhEFV1  ..................................................................................................................................
RhEFV2  ..................................................................................................................................
TryEFV  ..................................................................................................................................
LhFV    ..................................................................................................................................
LbFV1   ..................................................................................................................................
LbFV2   ..................................................................................................................................
EfFV1   ..................................................................................................................................
EfFV2   QRDSFFFKGEKMERPFVEVFE.PKIDTEASFLFYLK...NKNSDIFL.DSLDKLLKARAYKIKRTYRKTYASEFLFNNCRHLDNFKNVHQESY.......SPCGCSTAATRF.............KCYV.
CcFV21  ..................................................................................................................................
CcFV22  ..................................................................................................................................
CcFV11  ......P..TKKKK.................V......INDTVD....DDEDDD.....EEPERAVQIK.....TRHKC..MKCKKNINALGFDRHVFRCVGASCPFCGALFFSEKTCNNH..IPKCKKN
CcFV12  ..................................................................................................................................
PoFV    KLIKNFPSKKKNNKIILQTIECPLNDTISTTTINNTTTSNTTTDATTVNNSDTY.....DKKKLSVTVAFNSDAMKKQC..VKCYKLI..KNFDAHTKRCQGAQCEHCLKFFVNNTTKTFH.MKNSCKKL
PcFV    ........KSKSKK............TTTTTVVYKK..TNDATDTTTDDGDDDN.....NNKNTSPK.......MYLKCA.VNKYQNI..NKLV....INDKKTCKYCGKVFKQNWSKIRHENKQVCRK.
```

logo

```
LbEFV   ..
LhEFV   ..
LcEFV   ..
RhEFV1  ..
RhEFV2  ..
TryEFV  ..
LhFV    ..
LbFV1   ..
LbFV2   ..
EfFV1   ..
EfFV2   ..
CcFV21  ..
CcFV22  ..
CcFV11  V.
CcFV12  ..
PoFV    KK
PcFV    ..
```

X acidic (−)
X basic (+)
X polar uncharged
X hydrophobic nonpolar
