## Supplementary material for "Dating the origin of a viral domestication event in parasitoid wasps attacking Diptera": FiguresS13: Cluster115-Cluster431-Cluster37_LbFVorf107_[lef-4]_MSA.pdf

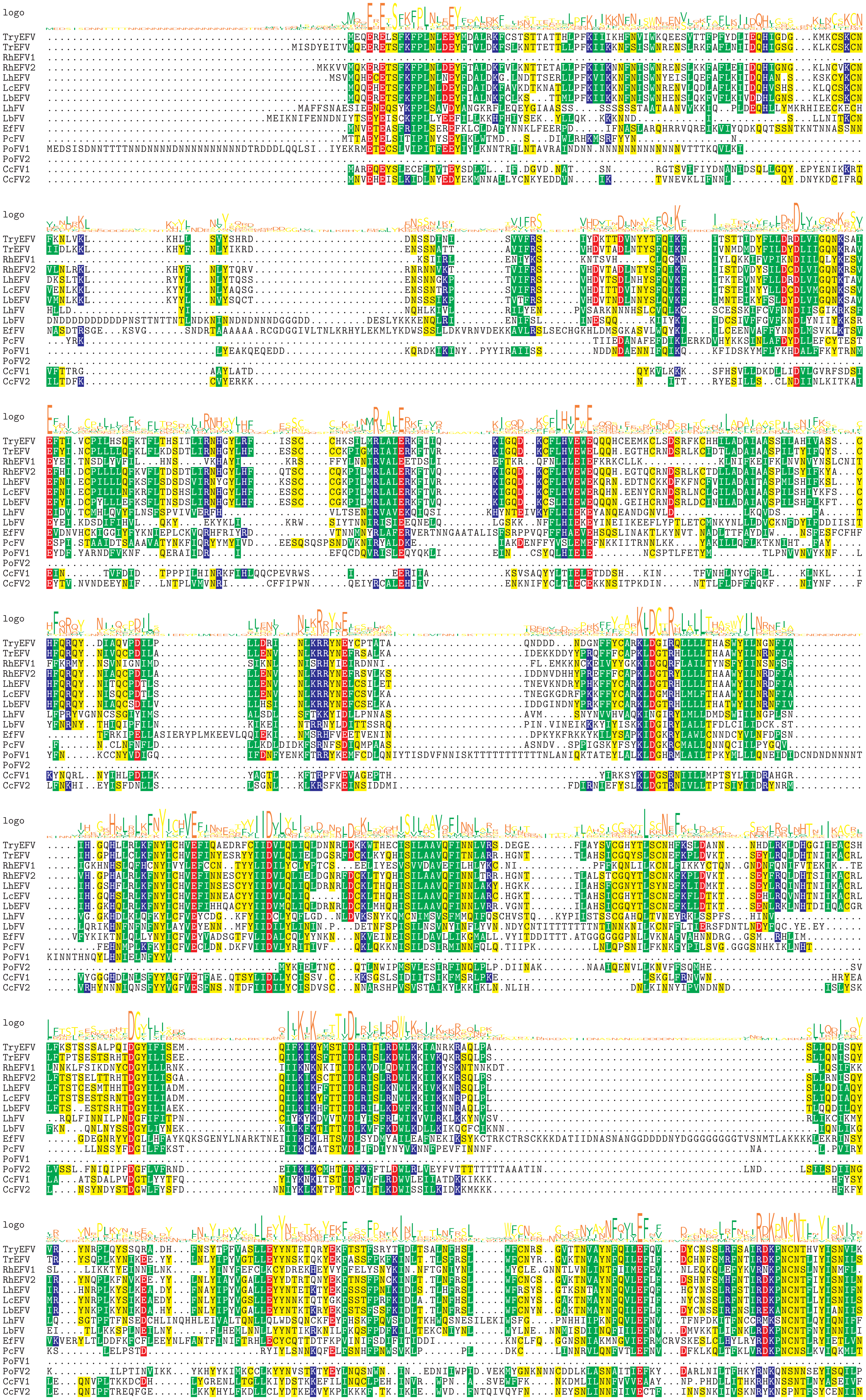
logo
TryEFV
TrEFV
RhEFV1
RhEFV2
LhEFV
LcEFV
LbEFV
LhFV
LbFV
EfFV
PcFV
PoFV1
PoFV2
CcFV1
CcFV2

```
logo
```
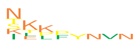

```
TryEFV  K.......
TrEFV   S.......
RhEFV1  ........
RhEFV2  N.......
LhEFV   N.......
LcEFV   N.......
LbEFV   KK......
LhFV    ........
LbFV    TK......
EfFV    TEPFYNVN
PcFV    I.......
PoFV1   ........
PoFV2   IKK.....
CcFV1   STLP....
CcFV2   NLK.....
```

X acidic (−)
X basic (+)
X polar uncharged
X hydrophobic nonpolar
