## Supplementary material for "Dating the origin of a viral domestication event in parasitoid wasps attacking Diptera": FiguresS13: Cluster126_redefined_LbFVorf58_[DNApol]_MSA.pdf

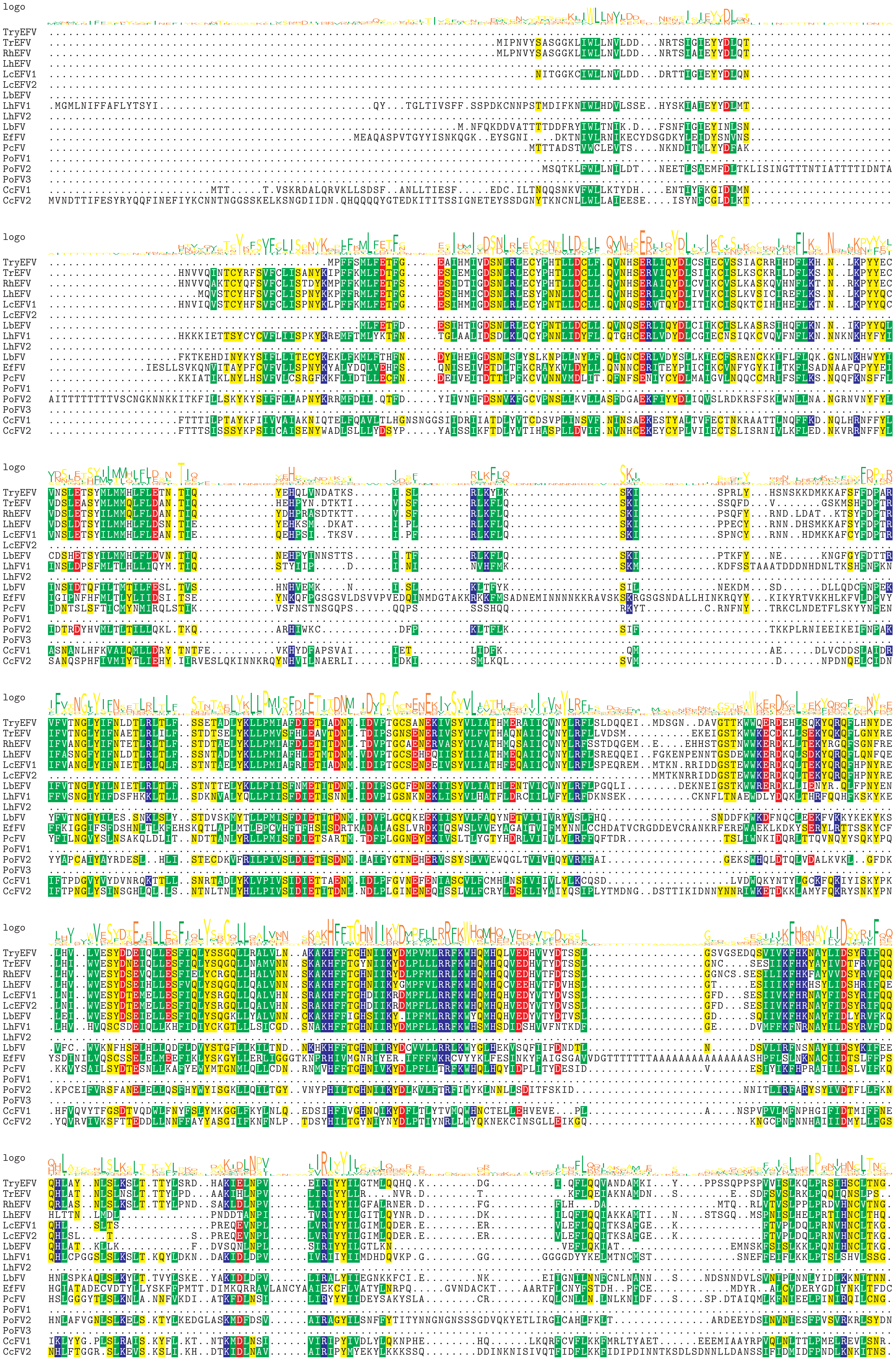
logo
TryEFV
TrEFV
RhEFV
LhEFV
LcEFV1
LcEFV2
LbEFV
LhFV1
LhFV2
LbFV
EfFV
PcFV
PoFV1
PoFV2
PoFV3
CcFV1
CcFV2

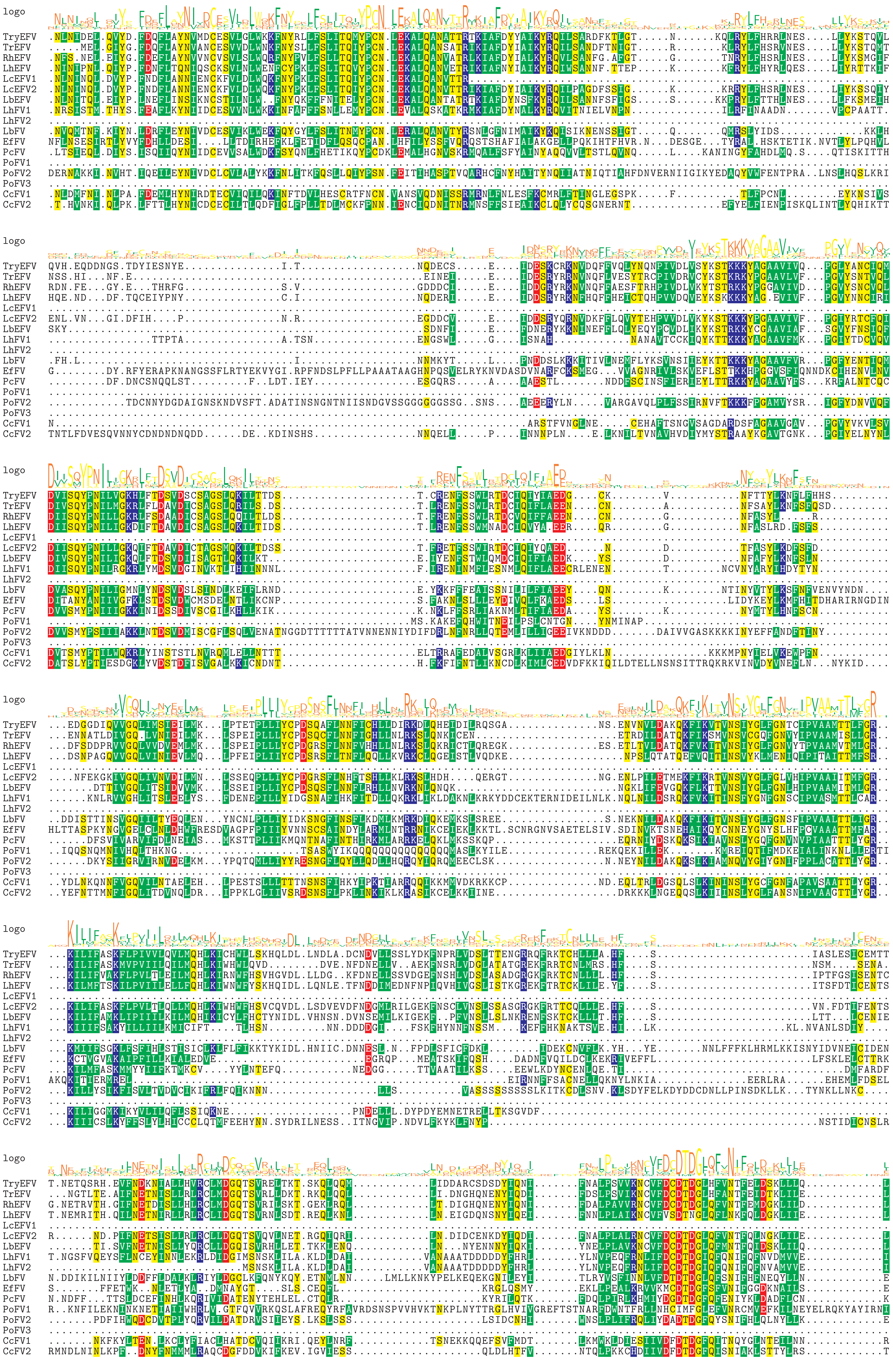
logo
TryEFV
TrEFV
RhEFV
LhEFV
LcEFV1
LcEFV2
LbEFV
LhFV1
LhFV2
LbFV
EfFV
PcFV
PoFV1
PoFV2
PoFV3
CcFV1
CcFV2

```
logo
TryEFV  LN........EEMV....QVFQMDNIIFDDKQSAYIFSLAKKKCTTFKFDPLCNVKFL..........N..EQCVIYIDDCSVRHNGYERNALPCLKYLSNFVA.....VLCLLI..........HQ
TrEFV   LN........QEIV....QAFQLENIIFTDKQFDHVFALAKKKYTTFKCPTGQVNHLE..........D..KEYVIYIDDCAVQHIGYEKNASPIFKYLCNFIA.....VLCMLI..........HQ
RhEFV   LN........LEVA....KVFQLDNIIFTLKQSDHVFSLAKKKYTTFNVSPLCKVKYL..........E..KETLISIDDCTIQHIGYERNALPCFKYVCNFIA.....VLCLMI..........HQ
LhEFV   LN........NEIG....KVFELDNIILSKKECAFVFALSNKKYTIFDLNPLEKVTFL..........K..NQYVIYIDDCKVQHVGYQ.AALPFFKYLCNFIA.....QICLLI..........HE
LcEFV1  .........................................................................................................................
LcEFV2  LN........DEIG....KVFQMDNIIFREKQHAYVFSLAKKKYTTFDFDPICGSTFL..........VKNQQHVIYIDNCCVKHVGYQKNALPCFKYLCNYVA.....IICLLI..........HR
LbEFV   LN........DEIV....NVFQLDNIIFTKKENDIVFSLAKKKYTIFKLNPICQLKFL..........HEQNKNVIYIDDCNLQHVGYQNNELSCFNYLSNFIA.....VICILI..........YQ
LhFV1   LN........DKIK....HVFEMDTIVFESKISKFVMSLTKKKYTTFNHVAKIHPDTY..........KLDNVGDFKTIQTTITHSGYERNALPCFKKLCDYVA.....LYCLCI..........FV
LhFV2   LN........DKIK....HVFEMDTIVFESKISKFVMSLTKKKYTTFNHVAKIHPDTY..........MLDNVGDFKTIQTTITHSGYERNALPCFKKLCDYVA.....LYCLCI..........FV
LbFV    LN........ENVQ....NVFEMNNIVFEATSSSYVVCLAKKKYTIFNYMYIPKVA...............ETTITINTDCKIIHKGYERNALLPFKNMCSFLV.....AVSFYI..........K.
EfFV    IH........LFVQ.....HYANKRIIFRATPLDVLLPLNKKMYAKIRCASVLLEHVRSSNDGKQMLHFDKSQKFW.....KEFEHVGYEKNSLAPIKRLFDIHV.....LFCYLM............
PcFV    LN........ELVS....EYFQVENIIFENTENMATLCIEKKRYTIYNRMPELNEFVESTATA..TTTEGQQQPPQRGVDYCRLKHSGYERNAVPHIKRICKYVA.....ILCYYM..........YV
PoFV1   VETNQSVLKNDYILLH..NVRLYKLINVRDENECVTRCLIKNIWTIFDNLS.......................RYIEIDHLTYLKNDI.....CTKMQ............T....RRNTLKR
PoFV2   LN........KNLCLHINGYRNDNSLSFTVKTMASVCALTKKKYTIYEKVLSRSD...............DIFPFYIKNARIKHSGYEKNSLPCFKRMCTFVA.....ALCYCIKCIYDDNNNNNKQ
PoFV3   ..............................................................................MRVLNIAVMKKILYHVLNVCALLWRLSVIALNVFMTITITTTNNNNNDN
CcFV1   FN........AAMK....EITNMPELNFEKAEVLSLILLTRKKYTTLKKGPVLSNLAV.......Y..KMYEKKWLPLQDFSIKHSGYERNALPHIKHICTVIA.....CYVFSY..........YH
CcFV2   FN........ESIR....KTFNMPNLNFESKHNLILISMKKKKYCQLLSGIVIENNKE......SYDFYKNNHNPVLLSEMKIAHSGYERNAVPIIKYICDVLA.....VKTFLN..........YR

logo
TryEFV  SKT.LRRYKFREIIWSVFNYLQ.KL..PPGQLYIQVKLN.RLKVNSARRRFIDKYTQT..YNGSLRAVYVVTTSPLLDKPSSSVELPLSTSSKHLEQLVMLDDYVCNVKHERNNANNQKMQLHHGFFLRT
TrEFV   GKT.LRLYKFREIIWSVFNYLE.NL..SPDQLYIRVKLN.RLKLDSAR.RFTDKYIRM..NSGVLRAVYVVKKSTFAEQSS.......SSSTKRLEQLVLLDDYIYSVKQQ.KSSKNKKMKLHYGFFLRT
RhEFV   GRT.LRPYKFREIIWSVFNYLD.AF..SPGQLYIRVKLN.RLKIDSARRRFIDKYTQT..YSGVLRAVYVVKTSTFTKQSCSNNSFLPSSSAKHSEHLVLLDDYLCSVKHQQKISKNERMKLHFGFFLRT
LhEFV   GKT.LRVYKFGEIIWSVFNYLE.SL..PPGQIYVRVKLN.RYKIDSGRRRFSEKYKET..NNGILRAVYIVNGSTVTKQSLNNNPFTLSNSKNHLNQIILLDDYIWSVQHQQNVGNSNNLQLHYGFFLRN
LcEFV1  ..........................................................................................................................
LcEFV2  GKA.LRFYKFREIVWSVFNYLE.CL..APGELYVRVKLNNRQRKDSARGRFINKYTQT..YNGVLRAVYVVNGDKFCEKSFNDVPLITSTGRKHLERLVLLDDYVWNVKHQRNNANSDKMQLHYGFFLRN
LbEFV   NKC...HYKFRDIIYSCFNYLE.NL..SPGEIYINIQLN.QLNVNSARLRFINKYTQI..YSGELRAVYIVKESLISNK..NDVSFLEQASLNNLEQLILLDDYIYSKKYQENVENHEIMQLHYGFFLRP
LhFV1   KGS.QMIIPFKELLFGVFNYLE.SV..KPDDLYMLLKMN.DISADSARKRFIDLYTSN..YTGSLKVVYIYNGSNVVTE................TIIPHFDYV.........AAPTTYVRHYAFFLRT
LhFV2   KGS.QMIIPFKELLFGVFNYLE.SV..KPDDLYMLLKMN.DISADSARKRFIDLYTSN..YTGSLKVVYIYNGSNVVTE................TIIPHFDYV.........AAPTTYVRHYAFFLRT
LbFV    KKI.VKPISFKRIIIFFSYLS.SL..KNDDLFVSIKLN.LIKNDCERKRFIDFYSKS..YTGSLKAVYIMENN................DTTLEKIILLTDYVNDI.........DKYELNLGFFLRP
EfFV    AKQYLRPFSLEELFRLYFRYLARHYSCGGGEDFILTDR...Q.....RNDMADHSA.A..VASSLPLTCIVHKHTLEKVLLKN.....DEPSEIQNVFVSLEYYKNCA.....SHRQPIFINYLYFLDK
PcFV    DNI.ASHITFRDLILSVYSYLKEHC..QPNELYINVVFK.KLKNNCARRRFIDSLTTS..YTGNVNAVFLYNYNTQEE................HYCTLQDYM.........QNSTKYTLNYPHFLRT
PoFV1   EQ...QNYEM.EMAQRFFNEIN.NQQCTPYEIAQLARLQ...........TISHFETH......................NGNPLLISVQQ....QIM...............................
PoFV2   QQQ.RQRQQFHKL....YNI......................................................................................................
PoFV3   DNSFINFITFDELILSCYAYMRLYLV.AGTDYYINIRLN.KQTNDSERKRFIDSRTLDVNTGGILAAMPIFRSEYAQ.............RFDEERLMSRDAYL.........INANHCWPHYSFFFRN
CcFV1   QPEKNPLGSLEDVMFDIYTYLLTVC..KKEDLYFNIRLQ.KIKPNCARKRFIEKYATN..YQGSLNAVYVVNEANDQEE................TLLLFDDYC.........INPSKYKLHYGYFLRN
CcFV2   YKY..FNVPFSYILEGIYTYIYMRK..DPYDLFINIALK.KIKPNSSRKRFIDRHATN..YTGTLKSVYIVNNEDFEDE................FLMLHDDYL.........RSKDNLKLHYAYFFRQ

logo
TryEFV  FGKIWFQQYCLATTVTTNDNDDTL.....Q....L.KNKSIMSIINQFFGDWLSRSEESR...RVDDLDHFLLSRFVIINKPS............
TrEFV   FGKIWYQQYCLSTTTDNNCD...K.....E....L.KNSSTMSIIHQYFDDWFSRSPQSR...LVDDLDNFLSNRFIVINRPLIF..........
RhEFV   FGKIWYQQYCLSVVNVNSSSNNAK.....E....FK.KSSTMSIINQYFDDWFSRSPQSR...RVDDLDGFLANRFIAINQQSL...........
LhEFV   FGKIWYKQYCQATNNMDDEIKSGS.....N....LKELGGTASIIDQYFDDWLSQSTQSR...SVDDLDSFLLNRFIVVNESSLSESLSSSVEMDT
LcEFV1  ...............................................................................................
LcEFV2  FGNIWYKQYCLSTSGKESNC.........G....SQELRSTVSIINQYFDDWLSRTSELR...RVDDLDDFLSNRFIVIKESSLS..........
LbEFV   FGKIWYKQYCLSTNNKVHNCNNAN.....N....FN.EMNTMLMINQYFDDWFSKSPQLR...CVDELDDFLLNRFIVIKQS.............
LhFV1   FAKIWHSQYTLVVQRLNINDDPSL.....QDQMHATKSFNIMLYVIECYNEWKKCENKTN...KFYNF...IKC.IEKIKK..............
LhFV2   FAKIWHSQYTLVVQRLNINDDPSL.....QDQMHATKSFNIMLYVIECYNEWKKCENKTN...KFYNF...IKC.IEKIKK..............
LbFV    FVKIWYQQYHIINNLEYN..................VKNNIVNQVLLYFSEWREKYNNPT...NVNIN.EIICNNL...................
EfFV    YIAYFLSLYNNTFVAAKTSVNESATENGGGGAAA..VSNHLS.....................................................
PcFV    YSQVWYNQFYIAKRQLEKITT...............TSLPIPTIMMNFFQEWLLIESGQR...SMDYVKLDIANCKLEL................
PoFV1   ...............................................................................................
PoFV2   ...............................................................................................
PoFV3   FGIVWYKQYHLAQKKNLTQIN...............SQKIMTILHANFQNFHQNLTTVLKDKTIQKLCKAMSS..EIIDYDTIVKK.........
CcFV1   YIKVWSVQLRHMMSSSQE..................YDFTPKHLHIQSFSSWLTGAEAEEGRRERVFLES.LLNNKIFIA...............
CcFV2   YIKVWFTQLQHAGVYLNNNTSNNSVRNSSNNNTNI.NETNLQKYQLQAFKQWLHLINNNN...NIFLENV........IYKDLISKSVY....IH.
```

X acidic (−)
X basic (+)
X polar uncharged
X hydrophobic nonpolar
