## Supplementary material for "Dating the origin of a viral domestication event in parasitoid wasps attacking Diptera": FiguresS13: Cluster194_LbFVorf5_MSA.pdf

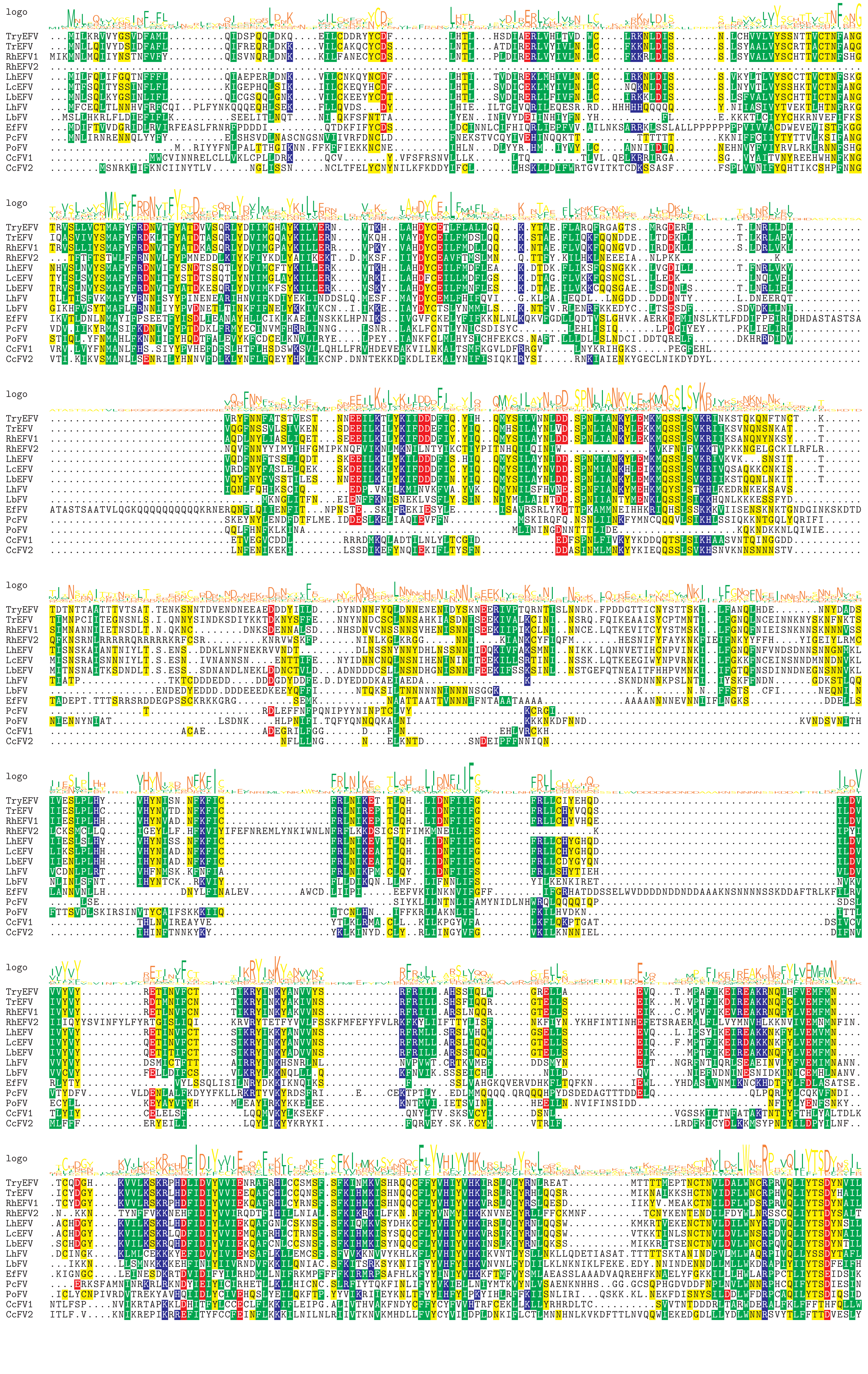

logo

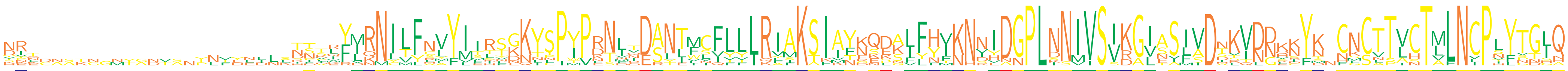

```
TryEFV  DR......................KSTRFMRNILFNVYIIRSGKYSPYPRNLTDANTMCFLLLRIAKSIAYNQDALFHVKNDIDGPLNNIVSIKGIASIVDDKVDNRNYK.CNCTTVCTMLNCPLYTGIQ
TrEFV   NR......................TTTRFMNNILFNVYVIRSGKYSPYPRNLDDANTVCFLLLRIAKSIAYDQDALFHIKNNIDGPLNNMVSIKGVASIVDAKVDNQKYK.CNCTTVCTVLNCPLYTGLQ
RhEFV1  NR......................TTPRFMRNILFNVYIIRSGKYSPYPRNLNDANTICFLLLRIAKSIAYKQEALFHIKNNIDGPLNNMVSIKGVASIVDTKVDDHKYK.CNCTTVCTMLNCPLYTGLQ
RhEFV2  DV......................DILEYLRNILFSMYITITDNYSPIPKSIQDSNTMLFYLLQIAKSIAYRKDKLFHVNHVIDGPLNNIVSVKGISSFVDNKINDKKYS.CNCTTLCTLLNCPVYTGI.
LhEFV   KR......................TSSMFMRNILFNVYIIRSGKYSPYPRNLVDANTMCFLLLRIAKSIAYKQDALFHVKNNIDGPLNNIVSIKGVASFVDTKVDNHNYK.CNCTTVCTMLNCPLYTGLQ
LcEFV   NR......................SNVQFLRNILFNVYIIRSGKYSPYPRNLIDANTMCFLLLRIAKSIAYKQDALFHVKNNIDGPLNNIVSIKGVASIVDIKVDDLNYK.CNCTTVCTMLNCPLYTGLQ
LbEFV   NR......................TTAQFMRNILFNVYIIRSGKYSPYPRNLTDANTMCFLLLRIAKSIAYKQEGLFHVKNDIDGPLNNIVSIKGITSIVDTKVDDQNYK.CNCTTVCTMLNCPLYTGIQ
LhFV    HIQDAAKG.MTADVA.YNFYNDNSISDLYVRNILFHLFIMKVDKYSPIPDNMVDSNLLCFLLLRIAKSIKFEQKSFFYIKNSVDGPLNNIVSIKGISAIADNSINNVDFK.CKCTTLCTMLNCPSFTGLQ
LbFV    YIK.....................DDIKYLKNITFLFYILHTENNNPIPKNIRESNIFIFYLLRLAKTIAFEKEKIFYINNVIDGPLNNIISVKGLASLSDSKVDDKIYD.CGCVTLCTLLNCPIYNGIQ
EfFV    DEL........................IRDRMIMYRYFMESNKKTCPYPFTVQERLEFVQLTIRYFKAKMFKEPKVFFTKNIHDGPLSNVVSSRAISSIVDHLVDDKKYM.CNSVPISTLLNVPSFTDHH
PcFV    STT.....................HPKVYIQNCVLYLYLSLYKQYPIMPDNITDANTFLVFFIRFMKSFITNEKNIYYIKNIHDGPLKNIVSVRAIESLIDSKILGKKYN.NKCTTVCTLLNCPNYSGLQ
PoFV    VKN.....................NFKLYIRNYVFCAYITLFNKITPWPNNNEDATIFSVVYIKAMKTIVTNNLKFYYMNNREDGPLINIVSVRGIESFADDRVNNKCYTNFSCATLNTVLNCPTLVNSQ
CcFV1   TPE.............PLTDELLLNNTIYIQNLTFALYMYTTKKTLPIVETFNDSLLWSILVLRVSKSIINGSDSILNF..NYANPLDNIVSVDGLNVMS.NVTDGERFN.NLSLTAHTALNLPHYGTGA
CcFV2   NNTCNSTNGNYGNTSNTNVELILKNNSLYIKNLTYCVYMFTLKKLSPTVDTDIESYLWNITYLRVIKSIINNSSSILNF..NYNNPLDSIVSVDGLNILA.NTQNGSSYN.HMSNSAHTAFNIPNYPTIN
```

logo

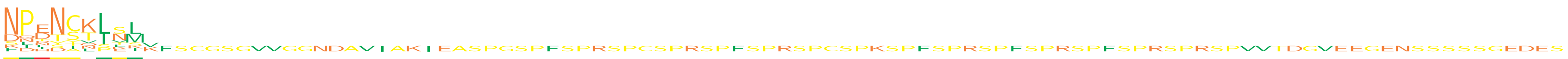

```
TryEFV  NPENCQLTL............................................................................................
TrEFV   NPDNCKISL............................................................................................
RhEFV1  NPDNCKLSL............................................................................................
RhEFV2  .....................................................................................................
LhEFV   NPENCKLNM............................................................................................
LcEFV   NPETCKLSM............................................................................................
LbEFV   NPENCKLNL............................................................................................
LhFV    DPINATIKM............................................................................................
LbFV    DPTTSVLNI............................................................................................
EfFV    KIHDSNSLKV...........................................................................................
PcFV    NLNNTTIV.............................................................................................
PoFV    SNDNYL...............................................................................................
CcFV1   FQRSSTPESKFSCGSGVVGGNDAVIAKIEASPGSPFSPRSPCSPRSPFSPRSPCSPKSPFSPRSPFSPRSPFSPRSPRSPVVTDGVEEGENSSSSSGEDES
CcFV2   NDLYI................................................................................................
```

X acidic (−)
X basic (+)
X polar uncharged
X hydrophobic nonpolar
