## Supplementary material for "Dating the origin of a viral domestication event in parasitoid wasps attacking Diptera": FiguresS13: Cluster279_LbFVorf10_MSA.pdf

```
logo
```

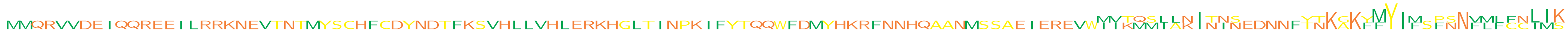

```
LcEFV   ..................................................................................................................................
TryEFV  ............................................................................................MVKMMLLKITIS....TTKAKYMYIFSPNNVLLFCIIK
LhEFV   ..................................................................................................................................
LbEFV   MMQRVVDEIQQREEILRRKNEVTNTMYSCHFCDYNDTFKSVHLLVHLERKHGLTINPKIFYTQQWFDMYHKRFNNHQAANMSSAEIEREVWIITQSIANINNNEDNNFYNKGKFFY...FSNFMFCNLIS
RhEFV1  ..................................................................................................................MYIM.........LMK
LbFV    ..................................................................................................................................
LhFV    ..................................................................................................................................
```

```
logo
```

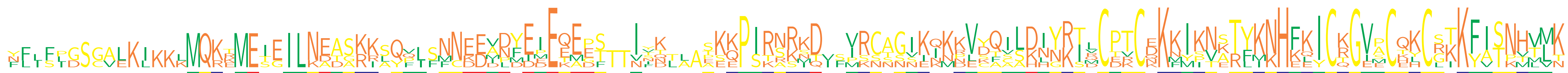

```
LcEFV   ...............MQKEMEIGILNEASKKLQPISNNEEVDYEMEQEES..IFK...SEKPIRNRKD..VRCAGVKKMKVCQILDIYRTQCPTCNKKIKNSTYKNHFKICKGVPCQKCSTKFISNHVVT
TryEFV  NLIFIGSGALKIKKKMQKKMEIEILKATARRSYQLSNCKDADYEEEKEGT..VTK...PKKPIRAKKD..IRCAGIKNKRVDQVLHLYKSRCPTCQKNIKNATYKNHFKICKGISCQKCGIKFISNHVMK
LhEFV   ...............MQRRMETEILNEASRKAQVLFNNEEANFEIEQEET..IIK...SKEPIRNRKD..VRCAGIKQKKVYQILDIYRTLCPTCKKKIKNSTYKNHFKICKGVPCQKCGTKFISNHLMK
LbEFV   FFLSPGSGALKIKKLMQKTMESEILNEASKKSQF..FNEEVNYEIEEE....NYQ...TKQPIRNRND..VRCSGIKQKKVYQILDIYKTLCPTCEKKIKNATYKNHFKICRGVACEKCSTKFISNHVMK
RhEFV1  YF.FLDSCVEKL............NDDGKI..........ILMEIETAPSTTIND..AESQPSTSRIQISISSTSSEINNVSAASKHNKIMCERCDKMYIVERLMKHKEYCQGVECHLCRKKYVTPVTLK
LbFV    ........................................MPFIPEEMPF..ISNLL.......SSSDVFPKNNNNIIKDLDFSSDNGRIVCDKCDKFMPVKRFKKHELICTGEMCKYCRKKFATKTTLK
LhFV    ....................................MNDDDDDDDEQEDSTTIVTTAAAKKPSKRKMT..M........PEKTCAKNNRIDCTVCEKKISTKTFINHKQICRGVACTHCLKKYATNMMLN
```

```
logo
```

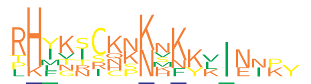

```
LcEFV   PKIGIIN.KHKYLIEIKY
TryEFV  RHYNSCQNKN........
LhEFV   RHFVTCKNKNKKVINNP.
LbEFV   RHYKSCKKNKN.......
RhEFV1  LHMKNRHSNSKKKIN...
LbFV    THITRSCPKMF.......
LhFV    RHLKICKKNVK.......
```

- X acidic (−)
- X basic (+)
- X polar uncharged
- X hydrophobic nonpolar
