## Supplementary material for "Dating the origin of a viral domestication event in parasitoid wasps attacking Diptera": FiguresS13: Cluster425-Cluster1270-Cluster1978_LbFVorf108_MSA.pdf

logo 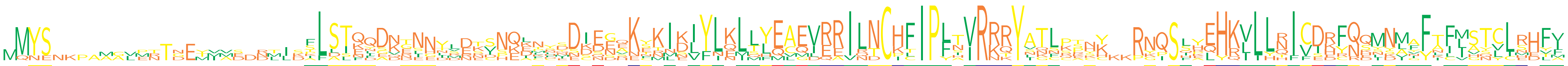

```
TryEFV  MQYS.....................ILSTQQADTNND.....QETHYDDEESKYNIKVYLNLLYEAEVRRILNCHFIPLAVRRRYATLPRNK..PYPSLYEHKILLQICDRFQEMNMIFAFVGTCLRDLV
TrEFV   .MYS.....................AFSTQYGKGNNYRDKSHQEMNNDIDGEKYKIEIYLQLLHEAEVRRILNCHFIPFTVRRRYVNLSSSY..RGQSSHEHKVLLHICDRFQQINMAFTFMSTCLHHFV
RhEFV1  ...........MTTNEYIVE.NMLSPLSINLENNL.Y...DSEILFMNKEDANNKLKIFLELTLQCQIEAIQTCKTIPYHIRKQYSTCRT........LELHKILYTIVEKRNSNDVFFKLIANLIE.YI
RhEFV2  .MYS.....................SLSTENDKRNNYSDRSNQKNSDDISGEKSKIDIYLKLLYEAEVRRILNCHFIPFSVRRRYATLPPYC..RNQSFPEHKVLLQICDHFQHMNMAFIFMSTCLRHFI
LhEFV   .MYS.....................FLSTKQDNENH.VETVNQDNDSDIEGQKCKIDIYLKLLYEAEVRRILNCHFIPFTVRRRYATL.SHY..RNQSQYEHRVLLRICDRFQQMNAAFTFMSSVLSHFI
LcEFV   .MYS.....................FLSTQQDKD.....KCNKDNYEDIDEHKFKIDVYLKLMYEAEVRRILNCHFIPFTVRRRYATLCPHY..RNQSHYEHKVLLRICDRFQQMNAGFTFISTCLRHFV
LbEFV   .MYS.....................FLSTKQDNENNNMDTSNKKKCDDSEEQKSKINIYLKLLYEAEVRRILNCHFIPLTVRRRYATLPACH..RNQSIHEHRVLLRICDHFQQMNMVFTFLSTYLRHFF
LhFV1   MMNENKPAAAVPNTDEMVADDDLIKSLDIEIS..SGLLES...LCLQECENVEETISLYIQMSFEVGLERVRNCKFIPLNIRNKYIQSQKNK.......RQHKLLLNIVSDCNDYYLYFLIIHYV.MEFF
LhFV2   ..................................................................................................................................
LbFV    ..................MS.RTIDKLLLQFDQTI.TDNNVPELFRIDKQSAKERMKLFLEITLQCQIQEINDCICIPLNVRKTYNKNKT........SALHKLLHNIVIKYNSSKIFYNCLATCCN.TM
CcFV1   ........MGLERTGETM.......TLASFDDVGEEILREIN.APYSSDNNRKTMNKLYNKLQLMLMDEEILTCPLIPLIIRKQYLGFKEKKKKRKISLKQYQLILNFFIEYRKLSLNYAILVQLLS.VA
```

logo 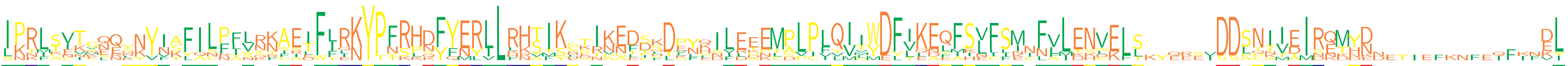

```
TryEFV  IPRIAVTVERQNYVTFILRTVRNAETFLRKYPFRPDFYERILRHSIKCKIAADDHDGFKSLEGEMPLPLQIMWDFLKEQFSVLSI.FLLDNVEFT......DDLCITDMRETYD..............KL
TrEFV   IPRLCIIGLQKKYIAFILPIVRNAEIFLRKYPFRHDFYERLLRHTIKYAIKGDTGDLNQILEEEMPLPLQIIWDFTKEQFSVFSM.FVLENCELL......DDSNIIDIRKMHN..............KL
RhEFV1  INNLKCVEN........NFLNQRCILFFQNYTNRPQYFNYLLLKTISEPRNNFSSLE.HDIRTNCPLPIVVLYDLVLDLMKVF.INN.L...............NNIAINETDDDETIEFKNFEQFKFEY
RhEFV2  IPRLSITG.QYKYIAFILPIVRNAETFLRKYPFRHDFYERLLRHTIKNTINEDSKDVNHIFEKEMPLPLQIIWDFIKEQFSLFSM.FVLENVELL......DDSNIIEIRQMHD..............DL
LhEFV   IPRLSVTGQQSYYISFILPFLRKAEIFIRKYPFRHDFYERLLRHSIKCTIKEESKDPYSNLEEEMALPFQITWDFIKEQFSVFSM.FVLENVELS......DDSNIVEIRQMYK..............DL
LcEFV   IPRLSVTCQELNYFSFILPFLYKAEIFIRKYPFRHDFYERLLRHSIKHTIKEDPKDPYKILEDEMALPLQITWDFIKEQFSLFSM.FVLENVELS......DDSNIVEIRQMYN..............DL
LbEFV   IPRLTVTSQQNNYIAFILPFLSKAEIFLRKYPFRHNFYERLLRHSIKRSIKEDIQDPYQILDEEMSLPLQIIWDFVKEEFSLFSM.FLLENVELS......DDSNTIEIRQMYD..............EL
LhFV1   LKKMTENKSN...NKLQCNLLNKISEWIIKYPYSLNFYNYVLHNAFKGDRKELDFDKLLIV........YVGMDFVKAFISNIIECLAIESDKLSKYQREVDDATTMTISND.D..............VI
LhFV2   ..................................................................................................................................
LbFV    LK...NKNE........SFFWKTSRDFFLNYPNSKKYFERLLFRVTYGKNVNFDLTE.E.L.EKLPPCFLVIYDYLENYFHLCFKNN.LTHL.....EESISSLPKIEIDEKNNN.........IFINDL
CcFV1   SKTFSEKMNEYNVNKYALFASNFFATFTENYPTKHSYQMLILKKTMSLETPETCS......LAPTDVIVQMVYEFIQELAIEFLNIFFMKKWK........DDPQAV......E..............EL
```

logo 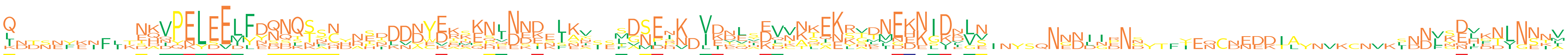

```
TryEFV  Q.........SQVPELEELYYQQQSSNDATD..DEEEMQNVDDE...K..MANEPNDVDDEAEVVK......NEKKIDELS.......................................D........
TrEFV   K.........TKVPELEELFDQNQTAQ..HIDDNYERTENLNSK.LKN..LGSEIK.VEKLIDVNNLEKYVDVEKN.VK....................................INNVNNIDTLNNNA
RhEFV1  LNTNNFKNFIK...........NNKKKC........KT..SQEEDKIAA...D..............SQE.................................EDKIA............S.........
RhEFV2  Q.........NKVPELEEMFDQNQTSH..PTKDAWDKAENINKE.LKI..SDSDIK.VEELSEVKNQAKRVDNEKNIIK....................................LNNVSEFKNVNNTL
LhEFV   Q.........NHVPELEELFDNNQSSCYNENDNNCERKKHLNNDESKV..LGNENV.VKNSYEQCNKEKKHLNNDELKVLS.....NENIIRNS...YEQCNEEDTAYNVKCNVKSDDLYQLKNLNNNV
LcEFV   Q.........NHIPELEELFYRHQTF..NSSDDDYEKKKNPNNK.LEV..FDSEKK.IKTLHKGNKKENRAENEKKIDILN....................................NFKEHENLNSII
LbEFV   Q.........NQVPELEELFEQQQSYNHKE...YYEAEEKLNND.LKV..SDSERNDLINESKLTKKENLGKIDDNGKNVN.....NNDIIPNN...T...............................
LhFV1   I.........EDDPN..VIFKDDL..C...KLVNYVKSKQINNISINEIEMVNLNK.IDVQKDDVASEKSTTMEHKIDDVV.......................................DV......V
LhFV2   .....................................................MEHK.IDDVVDVVAPEKSTTMEHKIDDVV.......................................DV......V
LbFV    IDDSEYEIFLTEKQQEYSEMLNNNTNNN........DDNKNIDETNIAY...D..............TIKKSMNEENSDNLNINYSQNNNDNDNDYTFIENCDNDDIL............KDLFTINSKM
CcFV1   L.........DANYKLDEDYDSSSS...HSDDDDYDSSSPEDDDEFSPSST..PS..V.......TTGERGDLGENGDLGE.....NGDLGENG.......................DLGEHGDLGE..
```

logo 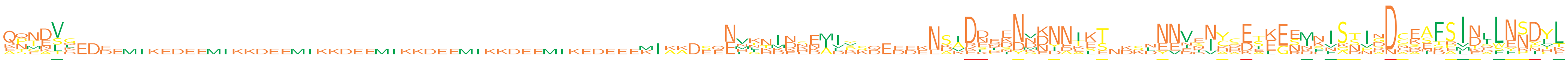

```
TryEFV  ....V...........................................................................DDENNVDNDKSS...NYVNSIYCEAKGRMSFSSNDDGKNSSVNQLYSDHL
TrEFV   QTNEVE......................................................NVKHIK.DMIP......NSIKEKDNMDINIKS....NNVENCSETEENMFNSNVNNCEAFSINLLESDYL
RhEFV1  ...DSQEDD............................................KIAADSQE...DDKNAVDSQEDDKNAADSQEDDKNAA...DSLEEDKNAADSLEEDKIA..ADGQEDEIEIFNNYLK
RhEFV2  QPNDVG......................................................NFKNINNEIIS......NSVRDRKNVENNIKT....NNIDICCEIEGNMKVSSIHDCEAFSMNSLESDCL
LhEFV   QQTDVD......................................................NVN.......Y......NSIDNNSNNNYNTKT....NNVIIHQETKESINISNIGDNESFSIDLLNNDYL
LcEFV   QQNNVL......................................................NN..................DKIIYNNNNTLT....NNVYNYDKSKESINISTINDSKAFSIDILNNDYL
LbEFV   ................................................................................ENCKNGKNNIET....NNVNVYHE...EKEISTISDCEAFPIDLLNSDYL
LhFV1   APEKS.........................................................TTMEHKID............................................................
LhFV2   ESMAL.........................................................LNINSEALV......EDADDDDDDD...........................DDDASTSINELNRNV.
LbFV    KNTETKEDEEMIKEDEEMIKKDEEMIKKDEEMIKKDEEMIKKDEEMIKEDEEEMIKKDEEEMIKKDEEMIKKDEEEENKKDEEENKKDEEEENKKDEEEENKKDEEEEN..K..KDEDEKRLKDWNNILE
CcFV1   .NGDLG......................................................EHG.................DLGEN..........................GDPGDASIFAITASFPPTT
```

logo 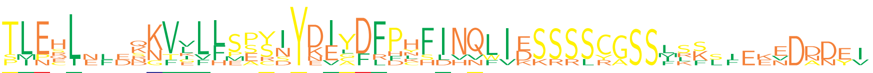

```
TryEFV  TMEHL...RKVVLHPPSNYNVGDLHLSLVN........................
TrEFV   TLEHL...QNVLLL.SSIYDIYDFPHFINQLVESSSSYGSSLSF..........
RhEFV1  ...................KEAFRFK............................
RhEFV2  TLESL...QNVLVFSSSIYDIYDFPHFINQWIDSSSSCASSMLSLIEENDANEV
LhEFV   TLERL...KKFVLLSPYIYEIYDFPCFINQLIESSSSCGSSLRK....EDDDYI
LcEFV   TLEHL...NKVIFLSPYIYDIYDFPYFINQLIDSSSSCGSSFKK....EDNDEI
LbEFV   TLECL...KKVGLLSACSYDILDFPHFIHQLIESSSSCGSSISSSFEKVDDD..
LhFV1   ......................................................
LhFV2   .............FMEYDYDE.................................
LbFV    SINEIELEDKTKLFEERNYKECFFDNHDNVFIKKRRLR................
CcFV1   PLPQLNFDSGVELL........................................
```

X acidic (−)
X basic (+)
X polar uncharged
X hydrophobic nonpolar
