## Supplementary material for "Dating the origin of a viral domestication event in parasitoid wasps attacking Diptera": FiguresS13: Cluster559-Cluster482-Cluster457-Cluster353-Cluster2746-Cluster2234-Cluster1338-Cluster1713-Cluster1176-Cluster1383-Cluster2222-Cluster2543-Cluster735-Cluster167_[lef5]_MSA.pdf

```
logo
LhFV   ..................................................................................................................................
PoFV   ..................................................................................................................................
PcFV   ..................................................................................................................................
LhEFV  ..................................................................................................................................
EfFV   ..................................................................................................................................
CcFV2  MAPSATTPKTAAITNNVSVESNTKNSMELNFFHDLCSFYFDNDPNKCVLFFSKFIKYIFERLAIDQAQNVYTLKDTPNPESYNGYIGNYQHDIIKNFSNIQELMSIFLKDSNEGGGLKIVHELNILTETK
CcFV1  ..................................................................................................................................
LbEFV  ..................................................................................................................................
TrEFV  ..................................................................................................................................
LcEFV  ..................................................................................................................................
RhEFV  ..................................................................................................................................
TryEFV ..................................................................................................................................

logo
LhFV   .....................................................LSRVLITEH.............N.............DDDDDP...........NPSNNSQSYIVDADST........
PoFV   ..................................................................................................................................
PcFV   ..................................................MEKISRILE...........................................................NLKI........
LhEFV  ..................................................................................................................................
EfFV   ..................................................................................................................................
CcFV2  LTGSFLNDTEELLVLQDIDDMEKRKKSLKCETKKTINSENQNTTTPTNSTLSQMLNIYLNDDIFNDDNSSSIKNTYKINDHMLEQKVIEASTAYNAANTNTNNTNINISDVSEEEIASSSTTTTSSSITT
CcFV1  ...........................................MERNEDILSDLINLFFDGE.............YKKFIQFITA..AAEIHKYD...........ALCGKTAEEIFSKLSTTY......
LbEFV  .................................................LLFHLSDLKI......................................................NFTVDDCNL........
TrEFV  ....................................................MIQQILL......................................................NFIVPNDNL........
LcEFV  ......................................................LKIFF......................................................NFAVGSCNL........
RhEFV  .........................................................LF......................................................NFIVQNINL........
TryEFV ..................................................................................................................................

logo
LhFV   ...APPQP....PSPPHQHTLANTTTPTPT............................................................TTT....................................T
PoFV   ...........................................................................................................MENEVEACMMTGNDNNK......
PcFV   ...IKTHL....KT........NNRSSGSG............................................................S.......................................
LhEFV  ..................................................................................................................................
EfFV   ....................MAATTAATTT............................................................TTT....................................T
CcFV2  KQKQEHQPQLSFDSEPHEEMTRNTKK..RKLNETIITKSTKRQKISSSEDIINVDDDNKLDTFLPPSALNTSTTEVRKKRKYQRKEKKDLKSSDNSSIEKKNNKKSNIEDILEEKEEKRNSTNNNNIVKR
CcFV1  ..NRPAQPS...AAPPKAHAKEVTEDEEPQLDQMTLTDLEKRK..............NEL.........KKKLTTEKKKSSLDKKRKKDMVTT....................................T
LbEFV  ...PSTHQ....Q.....................................................................................................................
TrEFV  ...PLSHQ....Q.....................................................................................................................
LcEFV  ...PELKT....S.....................................................................................................................
RhEFV  ...PSSHQ....Q.....................................................................................................................
TryEFV ...LTSQP....P.....................................................................................................................

logo
LhFV   TTTTTTPTPTTTTTTTTTTTQTTCSKRG.RKKNKISME..................DIHIISNDLFNLLNFKLPKDTDCKHQYVINNQ...QTRVGDEIETIMYVCVLCGQFKNYNR
PoFV   ........N.........AMMNVTNTKFNNDNTNVNIN..........NDNATFNIPSTTYEQLIRDTCEIPIPIDDECKHQYDGDKQKSVQFKSGDEGDKKYYDCKNCELRQMQ..
PcFV   GGSGSSGKSSSGGGGINNSRHTTGN..G.DGRNKINKS...............AKLELEFNSILAEELMNFPIPIEQPCKHKFVLLET...QTRRGDEIENETAVCILCGYSKNS..
LhEFV  .........................KKC.NKKMKINDS..................EAREIDKIIENLSSYKIPVESNCRHIFASSDA...QLRSGDEMESILFVCVLCGESKDYYR
EfFV   TTTAAAA.AAATTTTTTTTTKTSSNNLN.SKKYKHNIE..................T.DRKWKILKTQLCESLPI.DLCSHVYKISNI...VNRSGDEMESVLKICILCNDRKMG..
CcFV2  HRPKKLPKN.....QHPQQEENINNGEL.QKQDKIDLNNINDILTCLNSSDTNTVVDKKLLKLIVKKI.MNQTCTDVECDHKWNIERT..KQSRNGDEIENIVWVCELCKSVKSNT.
CcFV1  KKPGKKRRS..SSEESSSGCDGVGGGSG.DEKE.....................AIKAANIKEIARELLESPAPIDVGCAHKFIALTS..DQKRSGDELESVYYMCAHCKYVRP...
LbEFV  ........V..TVEQGHNVEDVKNNKKY.KTKNKIIDS..................ESLEIDKIIENLSSYKIPVESNCRHIFASSDA...QLRSGDEMESIVFVCVLCGESKDYYR
TrEFV  ........D..VSDQAGYVKNVKIKKKC.KRKTKINNS..................EALEIDKIIEDLSSYKIPVESNCQHIFSSSDE...QLRSGDEMESIVFVCVLCGESKDYYR
LcEFV  .............DQICNVKNVKKTRKC.KKKLKMNDS..................EAREIDKIIENLSSYKIPVESNCRHIFTSSDA...QLRSGDEMESIIFVCVLCGESKDYYR
RhEFV  ........T..TSDQADHVKNVKVKQKS.KKKMKIDNS..................AAQEIDKIIENLSSYKIPVESNCQHIFSSSDE...QLRSGDEMESIVFVCVLCGESKDYYR
TryEFV ........P..PIVHAEVFEDIKSKRKR.TKRMETVDA..................NAHEIDEIVENLSLYKIPVESSCRHIFASSDA..QQMRSGDEMESIVFVCVLCGESKDYYR
```

- X acidic (−)
- X basic (+)
- X polar uncharged
- X hydrophobic nonpolar
