## Supplementary material for "Dating the origin of a viral domestication event in parasitoid wasps attacking Diptera": FiguresS13: Cluster653_LbFVorf35_MSA.pdf

logo 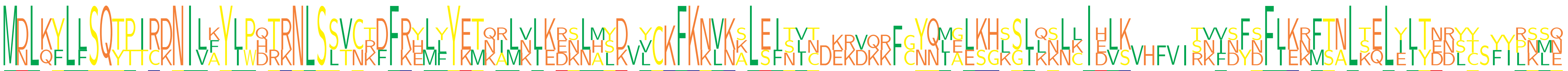

```
TryEFV MDLQYLFSQYPTRKNILKYLWDTRNLSSTNTFFKELLYETKILNIEDSLASDVLCKFKNVKKLEISVC........YNMELKHLGLLNNLIDLKVHFVIRNIDYSFLKNFTALKQLEITDRLC...LSMQ
LbFV1  MDLKFLLSQTTICDNIVAYLPQTRNLSLVCKDFRHLFYKTQRLKLKRNLHYK.VCKFKKLKSLEFNLT.KRVKKFCYQLGESHSSIQKLCIEVK.....SVVNFDFIKKFTNLTELYLYNNSTSFIPKLN
LbFV2  MNLKYLLSQTPIRDNILFYIPHRKNLSSVCRDFRYMYYEMNAMVLKEKNMLD.Y..FKNVNALSITINDEKDQRFGNQIALKGKSLKSLKIHLS.....TKFSFNFLERMSNLSELILTEDYY.YYRNSE
```

logo 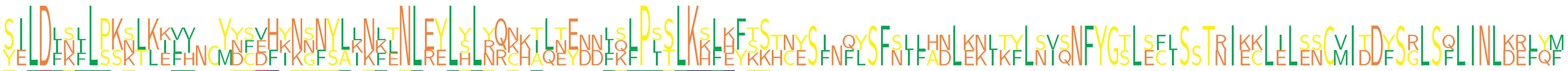

```
TryEFV SELDFSILPSNTLEVINCMYCDHKNSNYLKNFTNLRELYLYRC.AQEYDNIQFPISLKKLKYSS..................................................................
LbFV1  YILDIKLLSKKLKILV..YDFVFINGNYLINLENLEYLSLNQNHTLTE..LKLPLSLKSFEFKKTCESFNFYSFNIFADLKKIKFLNISNFYGTLECISSTNIECLILESCVITDFSRLSFLINLKEFYM
LbFV2  SILDLNFLPKSLKKFH..YNSEHYKNFSALKKLNLEYLHLRQKKILNENDFSLPSTLKHLHFTSHNYSI.QLSFSLLHNLENLTYLSVQNFYGSLSFLS.TRIKKLELSNCMIDDYSGLSQLINLDRLQF
```

logo 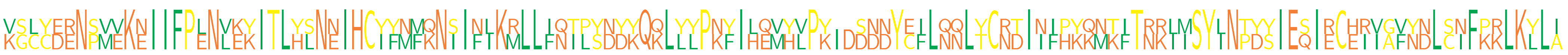

```
TryEFV ..................................................................................................................................
LbFV1  KSCYDRNPMVKNIIFPENLEKITLYSNNIHCIFMMQNIIFLKMLLINTLYNDKQQLYLPKYILQMYVPK..SNDYCFLNQLTCRDIIFPKKMKITRKIISYLNTYSIESIECHIVAFNNLCIFPRLKYLI
LbFV2  VGLCEENSVEKEIIFPLNVKYITLHLNEIHCYYNFKNSINIKRLLFQIPSDYYQKLLYPNFIHEVHLPYIDDDNYEILQNLYCNTINIHYQNTFTNRLMSYINPDYIEQIRCERIGVYDLSNFKKLKLLA
```

logo 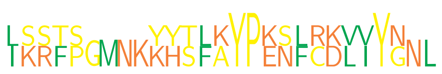

```
TryEFV ...........................
LbFV1  LSSTSGMNKKHSFAYPENFRKLIYNNL
LbFV2  IKRFP....YYTLKYPKSLCDVVYG..
```

- X acidic (−)
- X basic (+)
- X polar uncharged
- X hydrophobic nonpolar
