## Supplementary material for "Dating the origin of a viral domestication event in parasitoid wasps attacking Diptera": FiguresS13: Cluster1567_LhFV_contig_22588_orf001_MSA.pdf

logo 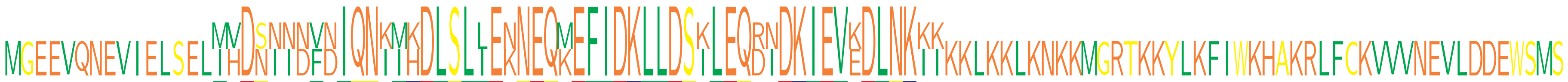

```
LhEFV   MGEEVQNEVIELSELIVDNNNNVNIQNIMKDLSLLEKNEQMEFIDKLLDSILEQRNDKIEVKDLNKIIKKLKKLKNKKMGRTKKYLKFIWKHAKRLFCKVVVNEVLDDEWSMS
LhFV    ...............MHDSIIDFDIQNKIHDLSLIENNEQKEFIDKLLDSKLEQDIDKIEVEDLNKKK.............................................
```

- X acidic (−)
- X basic (+)
- X polar uncharged
- X hydrophobic nonpolar
