## Supplementary material for "Dating the origin of a viral domestication event in parasitoid wasps attacking Diptera": FiguresS13: Cluster2164_LbFVorf105_MSA.pdf

```
logo
```

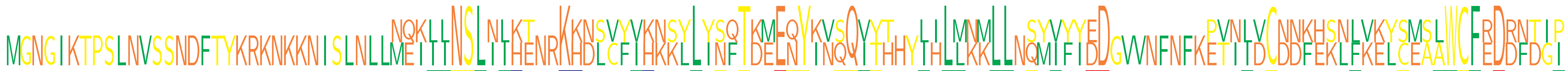

```
RhEFV  ................................MQKIINSLILKT..KHNSVYIHNSLLYSQTKMEQYKVQQVTTHYLHLLNMLLNQMVFYDD........EVILVCDDKHKLFKEYCMALWCFEDRFTGP
LbFV   MGNGIKTPSLNVSSNDFTYKRKNKKNISLNLLNEILLNSLNIHENRKKDLCFVKKKYLINFTDEENYINSQIYH..IILMKKLL.SYIYIEDGVVNFNFKPTNIDCNNFESNLVKLSESAWCFRDDNDII
```

```
logo
```

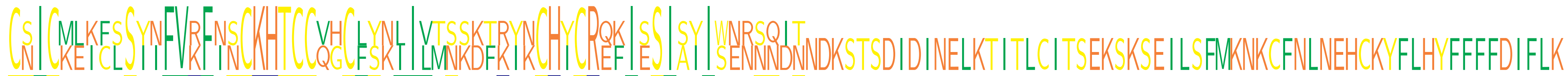

```
RhEFV  CSICMEICSSYIFVKFISCKHTCCVHCLSKLILMNSDTKYKCHYCRQKISSIAYIWERSQIT...........................................................
LbFV   CNICKLKFLSINFVRFNNCKHTCCQGCFYNIIVTSKKFRINCHICREFIESISIISNNNNDNNDKSTSDIDINELKTITLCITSEKSKSEILSFMKNKCFNLNEHCKYFLHYFFFFDIFLK
```

X acidic (−)
X basic (+)
X polar uncharged
X hydrophobic nonpolar
