## Supplementary figures and images for "Dating the origin of a viral domestication event in parasitoid wasps attacking Diptera"

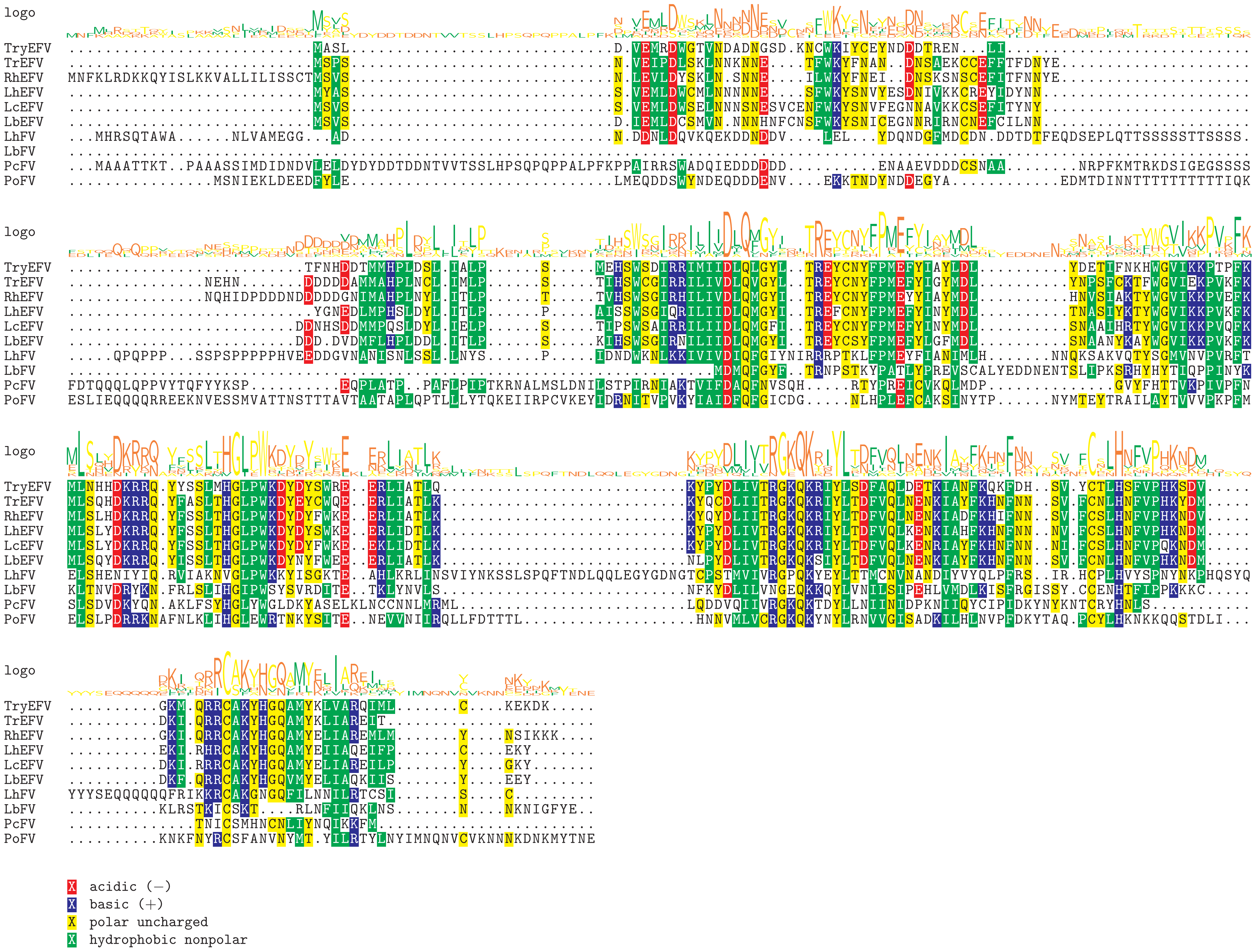

Supplement: FiguresS13 [file 595704_file03.zip › FiguresS13_All_Eucoilini_filamentous_alignments/Cluster186_LbFVorf94_MSA.pdf]

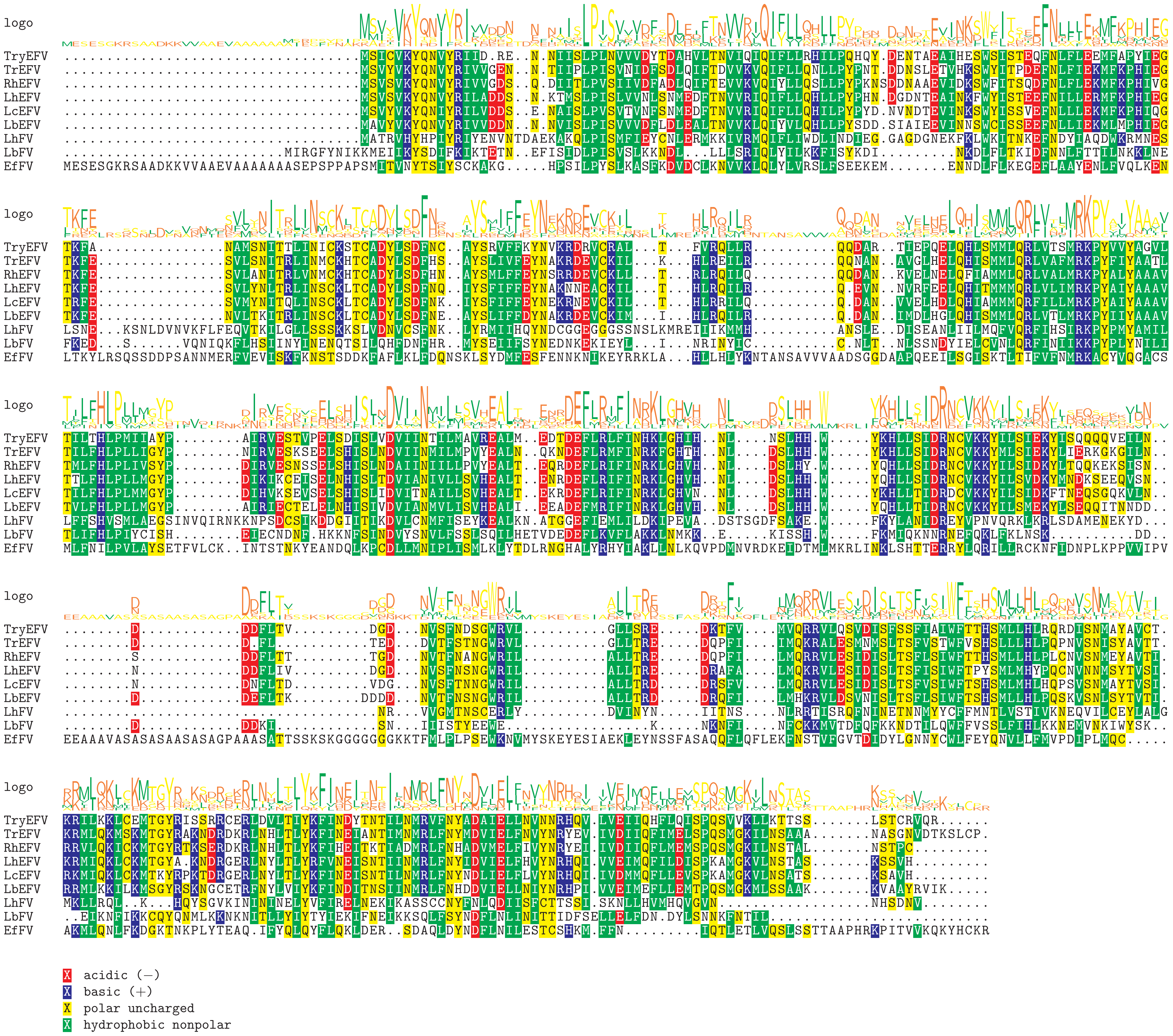

Supplement: FiguresS13 [file 595704_file03.zip › FiguresS13_All_Eucoilini_filamentous_alignments/Cluster27-Cluster1722_LbFVorf83_MSA.pdf]

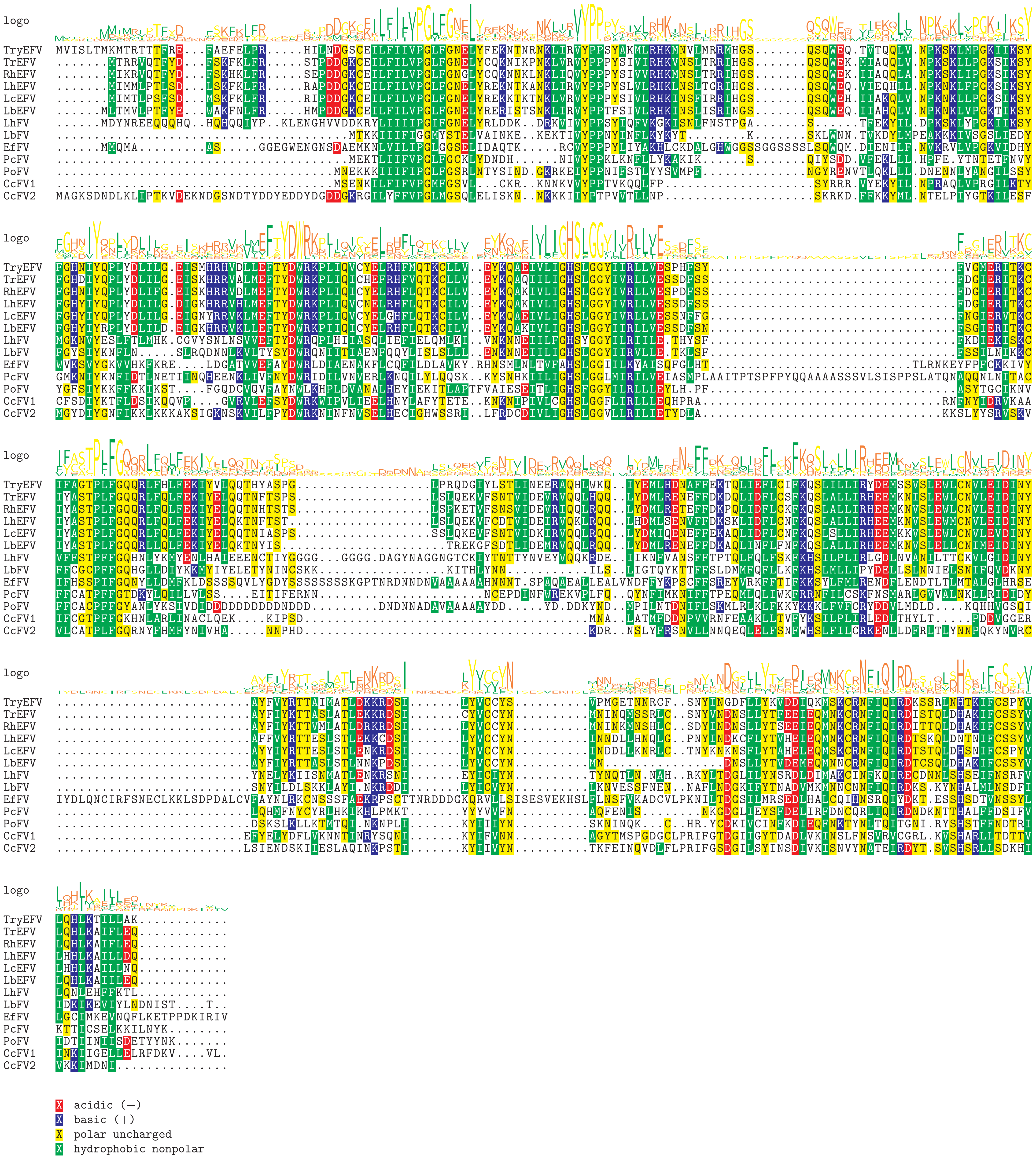

Supplement: FiguresS13 [file 595704_file03.zip › FiguresS13_All_Eucoilini_filamentous_alignments/Cluster440_LbFVorf60_[lcat]_MSA.pdf]

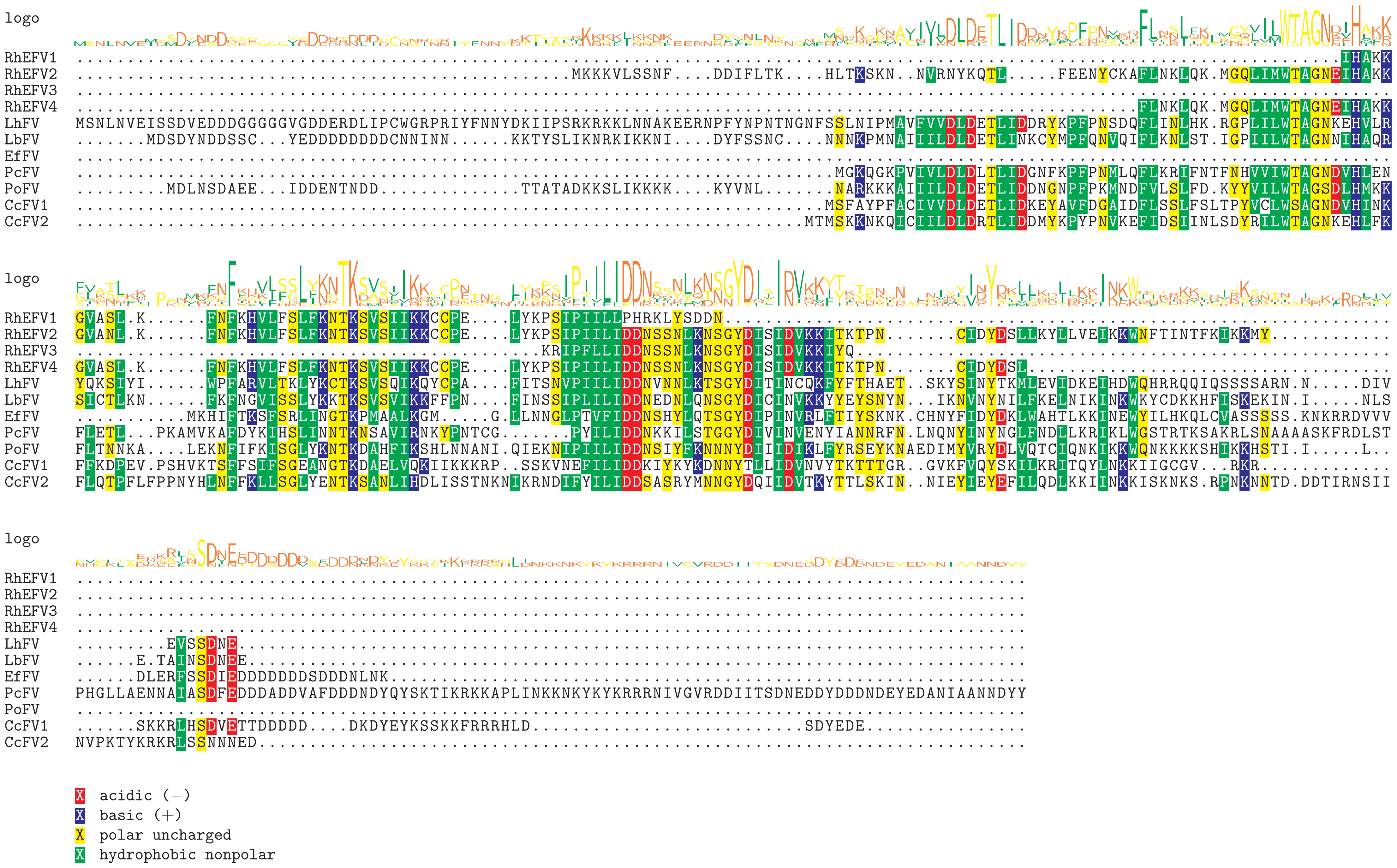

Supplement: FiguresS13 [file 595704_file03.zip › FiguresS13_All_Eucoilini_filamentous_alignments/Cluster38_LbFVorf19_[38k]_MSA.pdf]

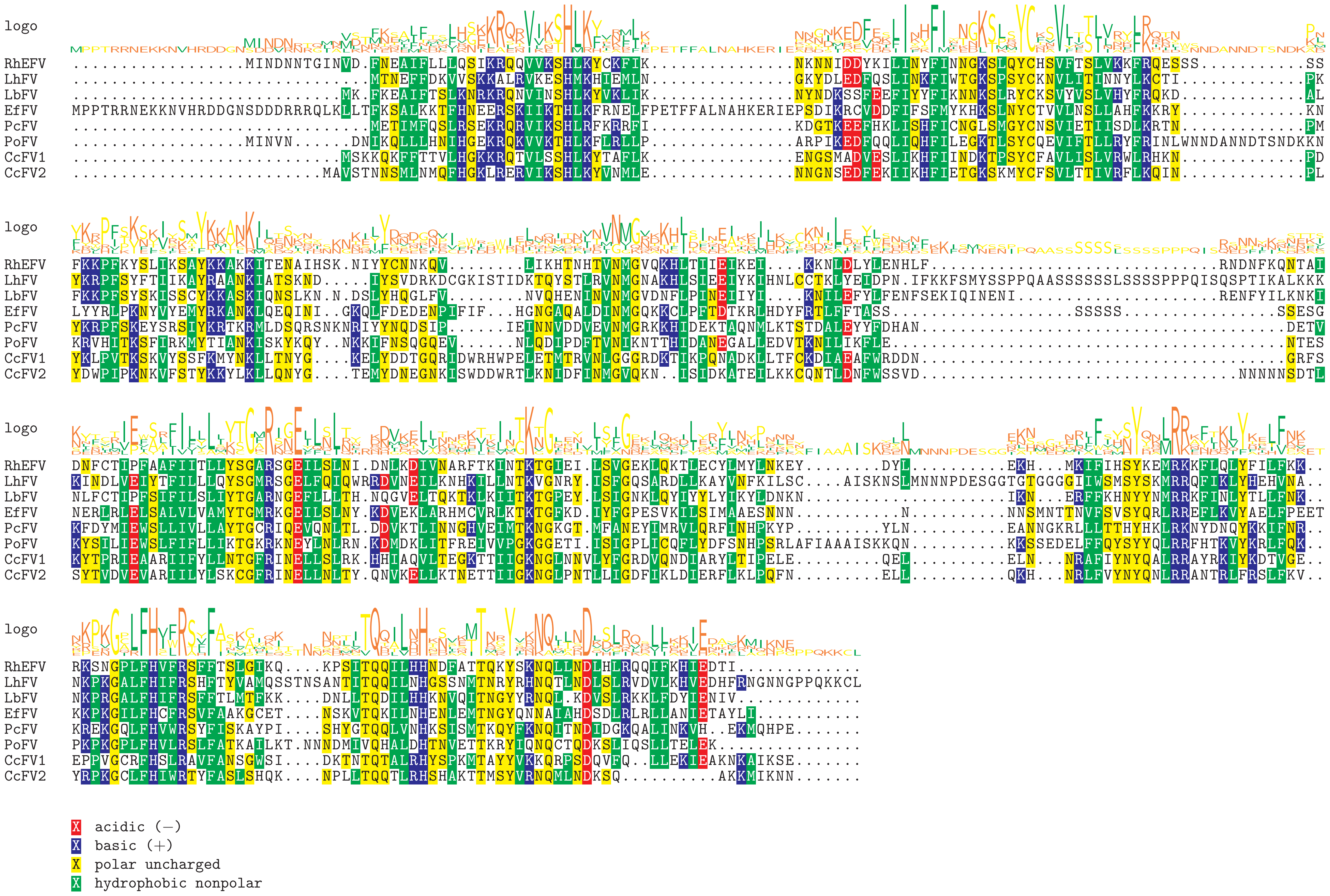

Supplement: FiguresS13 [file 595704_file03.zip › FiguresS13_All_Eucoilini_filamentous_alignments/Cluster22_LbFVorf2_[Integrase]_MSA.pdf]

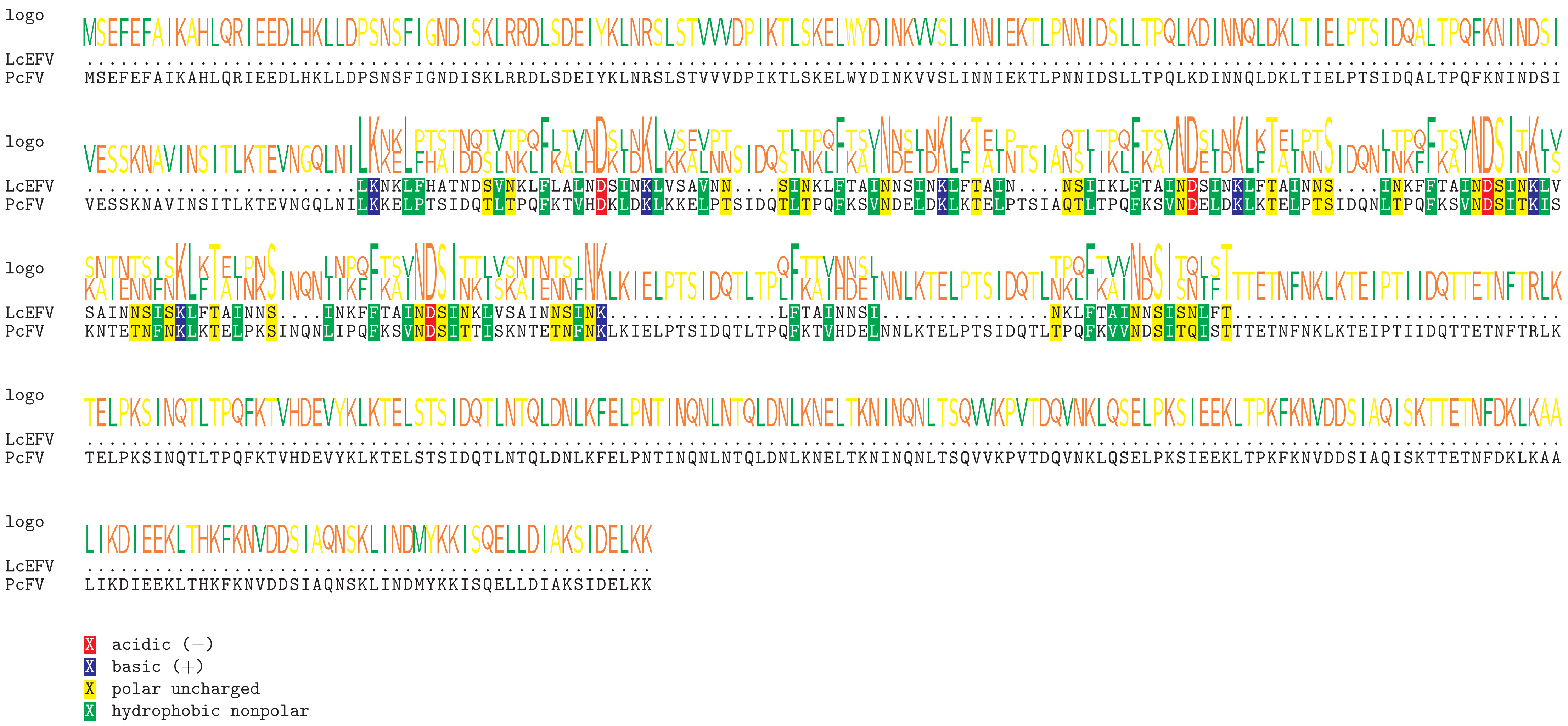

Supplement: FiguresS13 [file 595704_file03.zip › FiguresS13_All_Eucoilini_filamentous_alignments/Cluster1472_PcFV_scaffold_2882_orf004_MSA.pdf]

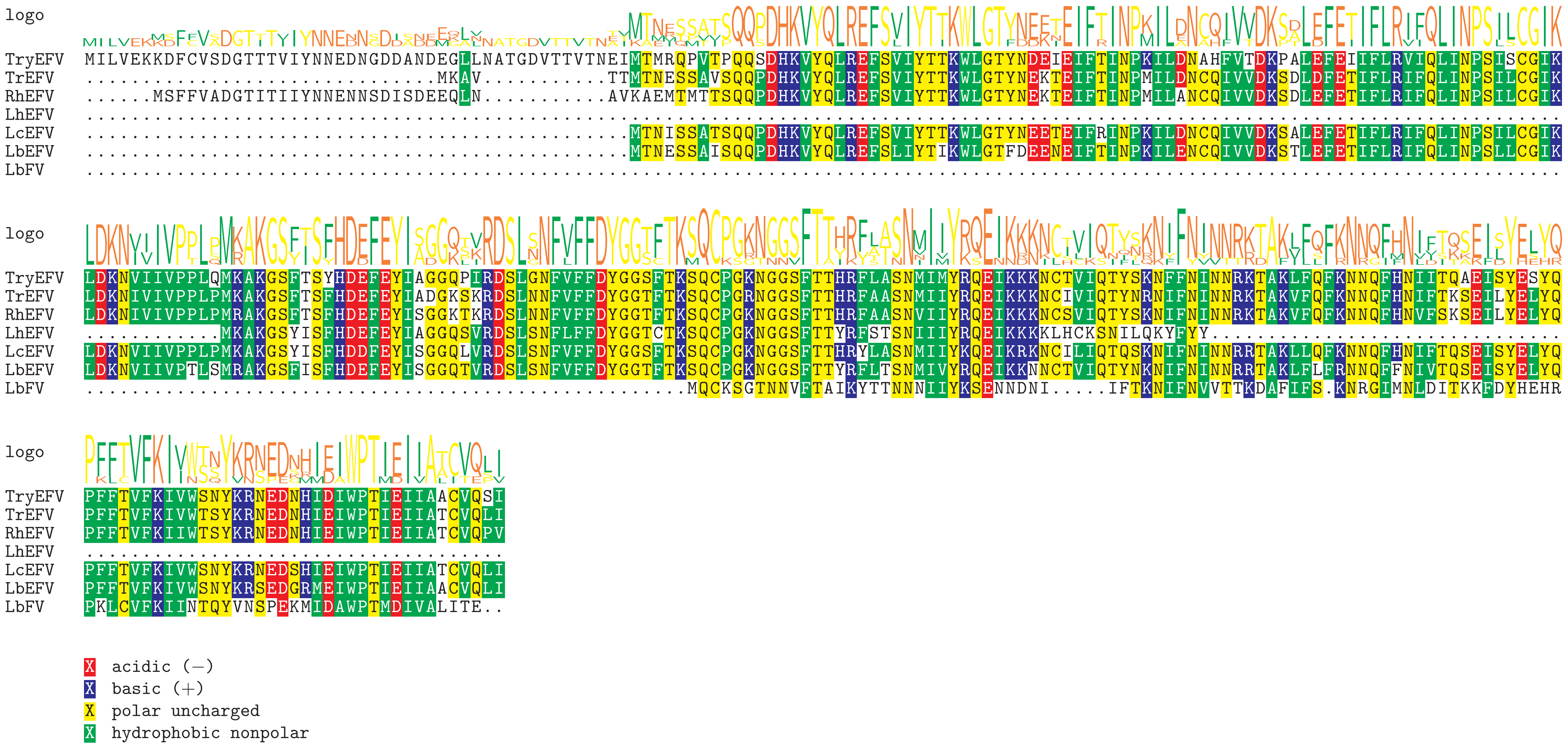

Supplement: FiguresS13 [file 595704_file03.zip › FiguresS13_All_Eucoilini_filamentous_alignments/Cluster258_LbFVorf72_MSA.pdf]

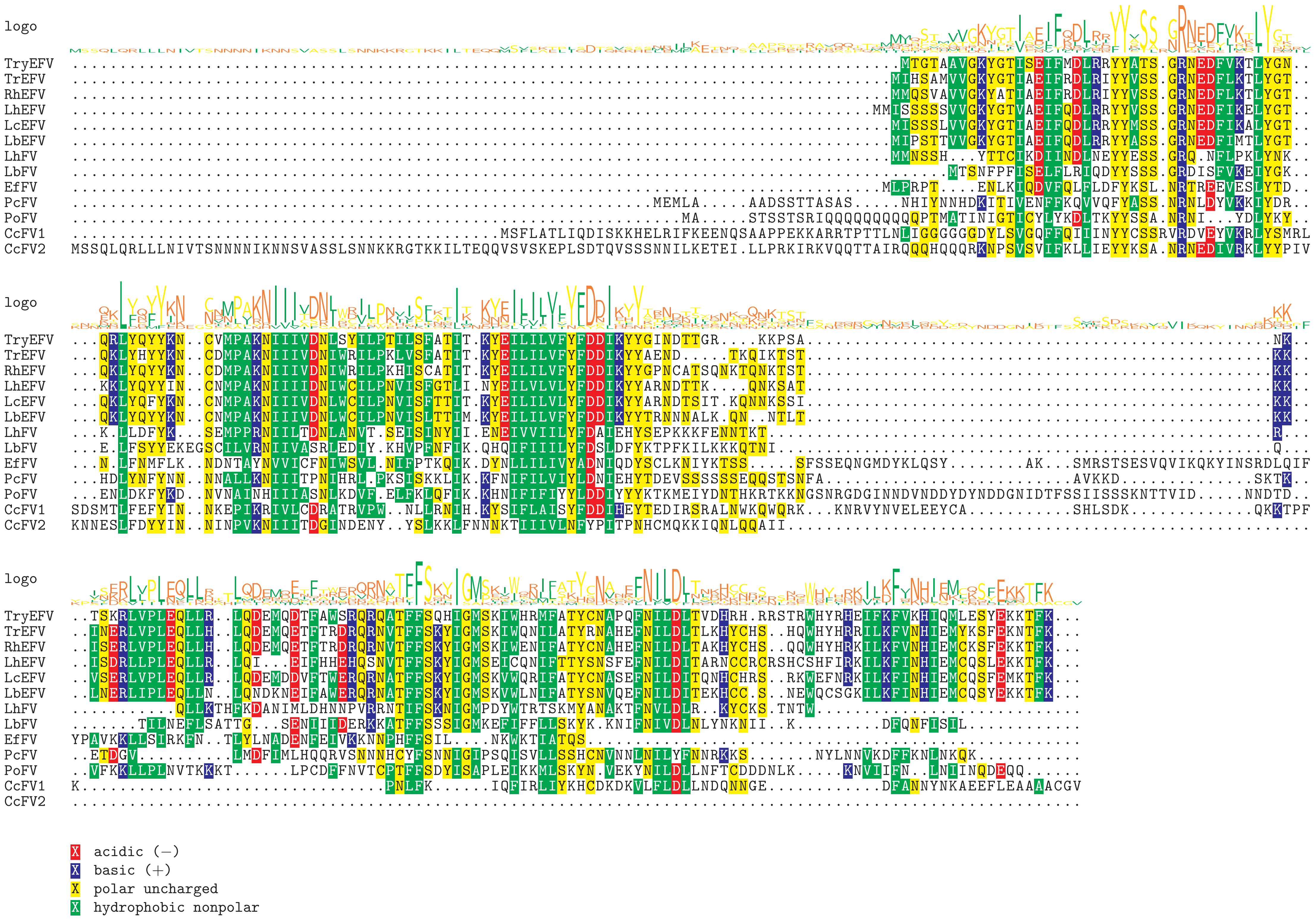

Supplement: FiguresS13 [file 595704_file03.zip › FiguresS13_All_Eucoilini_filamentous_alignments/Cluster544_LbFVorf87_MSA.pdf]

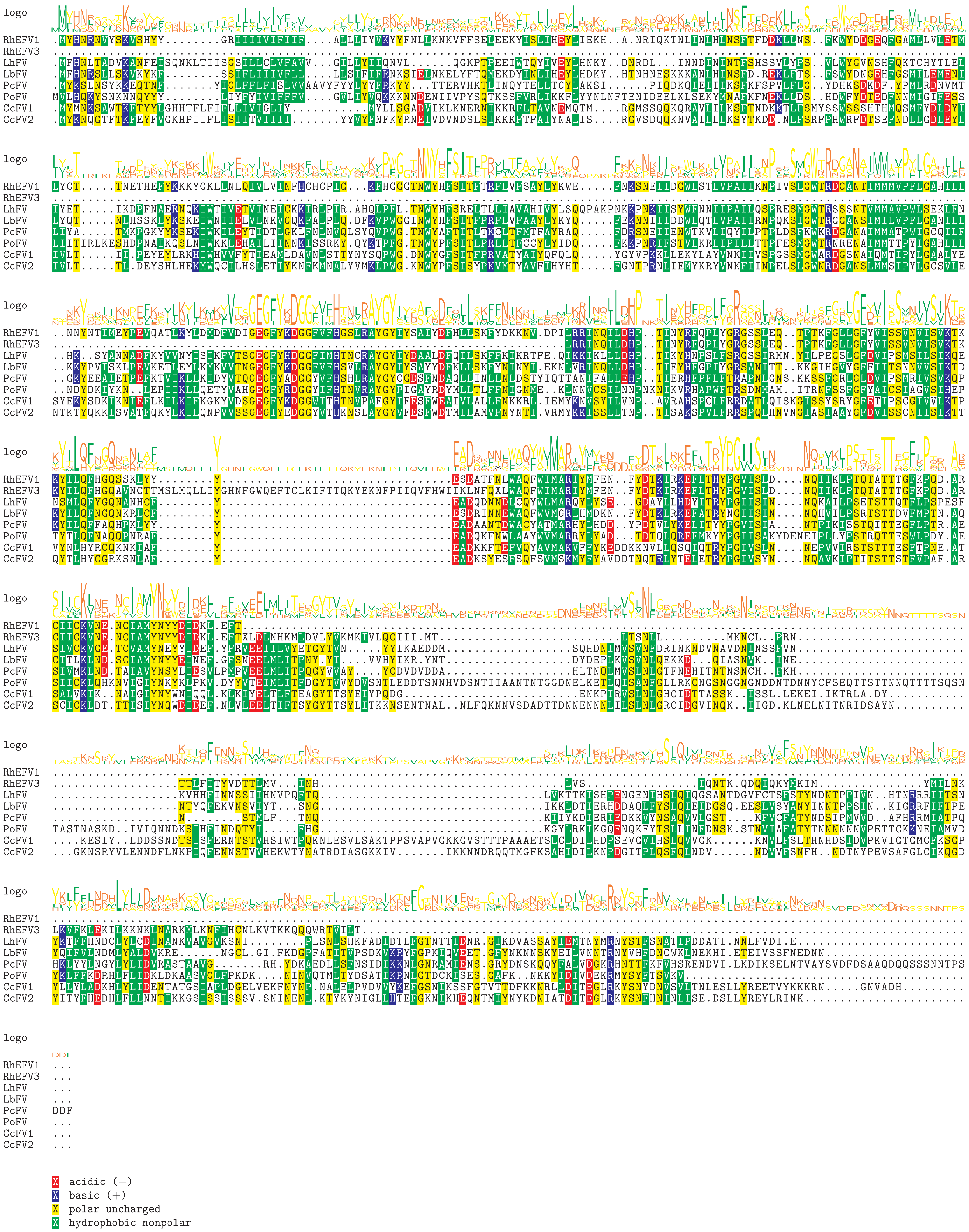

Supplement: FiguresS13 [file 595704_file03.zip › FiguresS13_All_Eucoilini_filamentous_alignments/Cluster45_LbFVorf106_[odve66]_MSA.pdf]

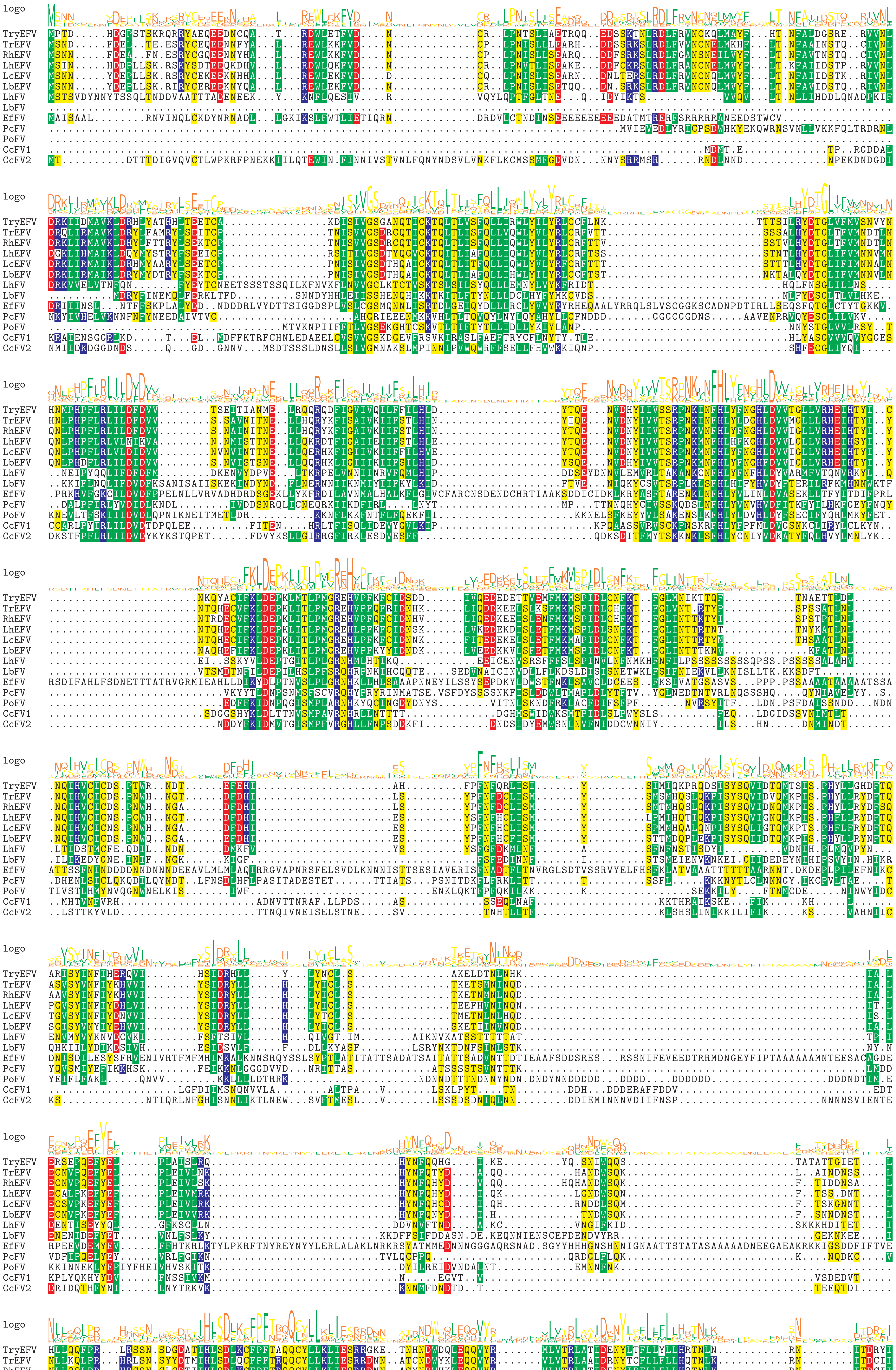

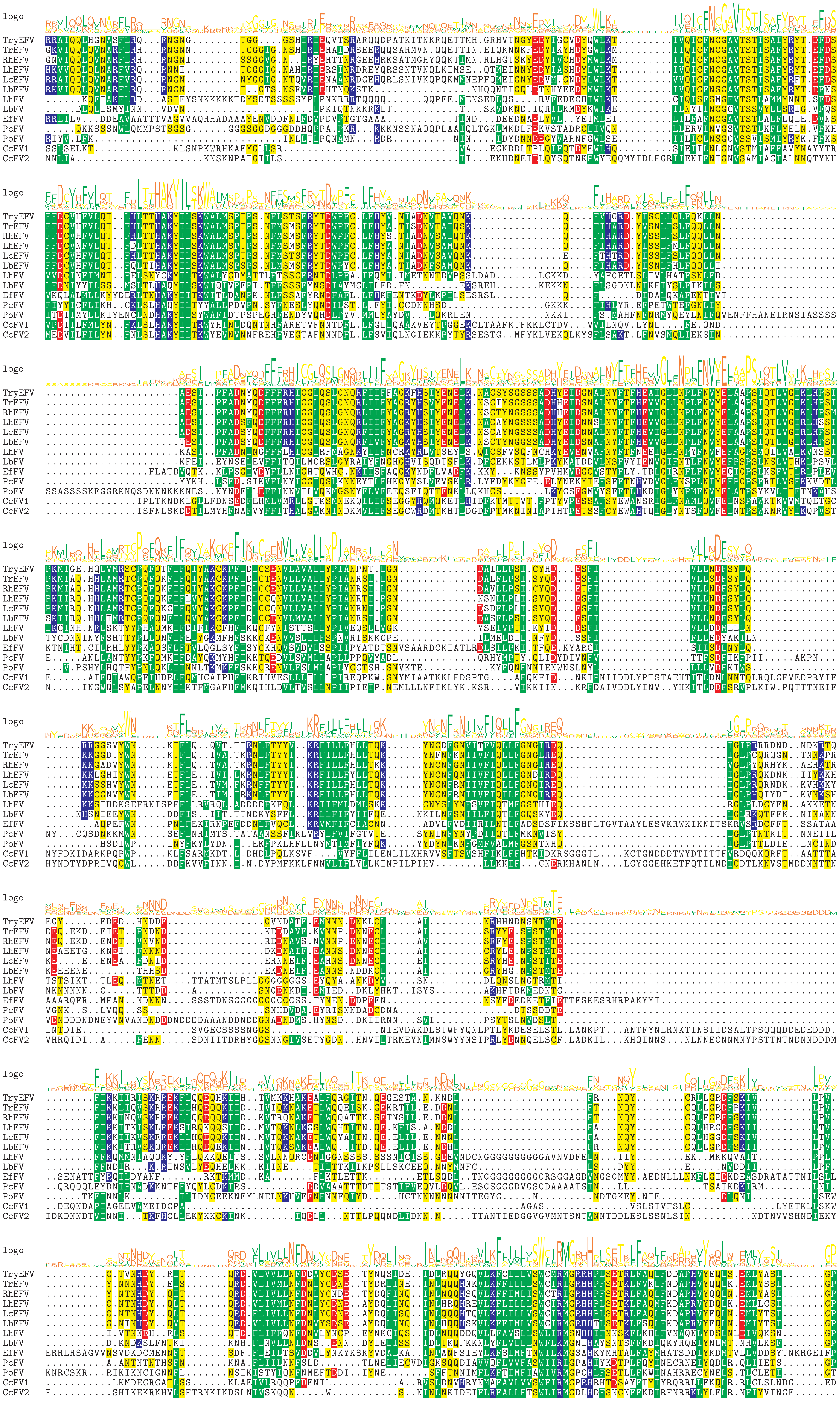

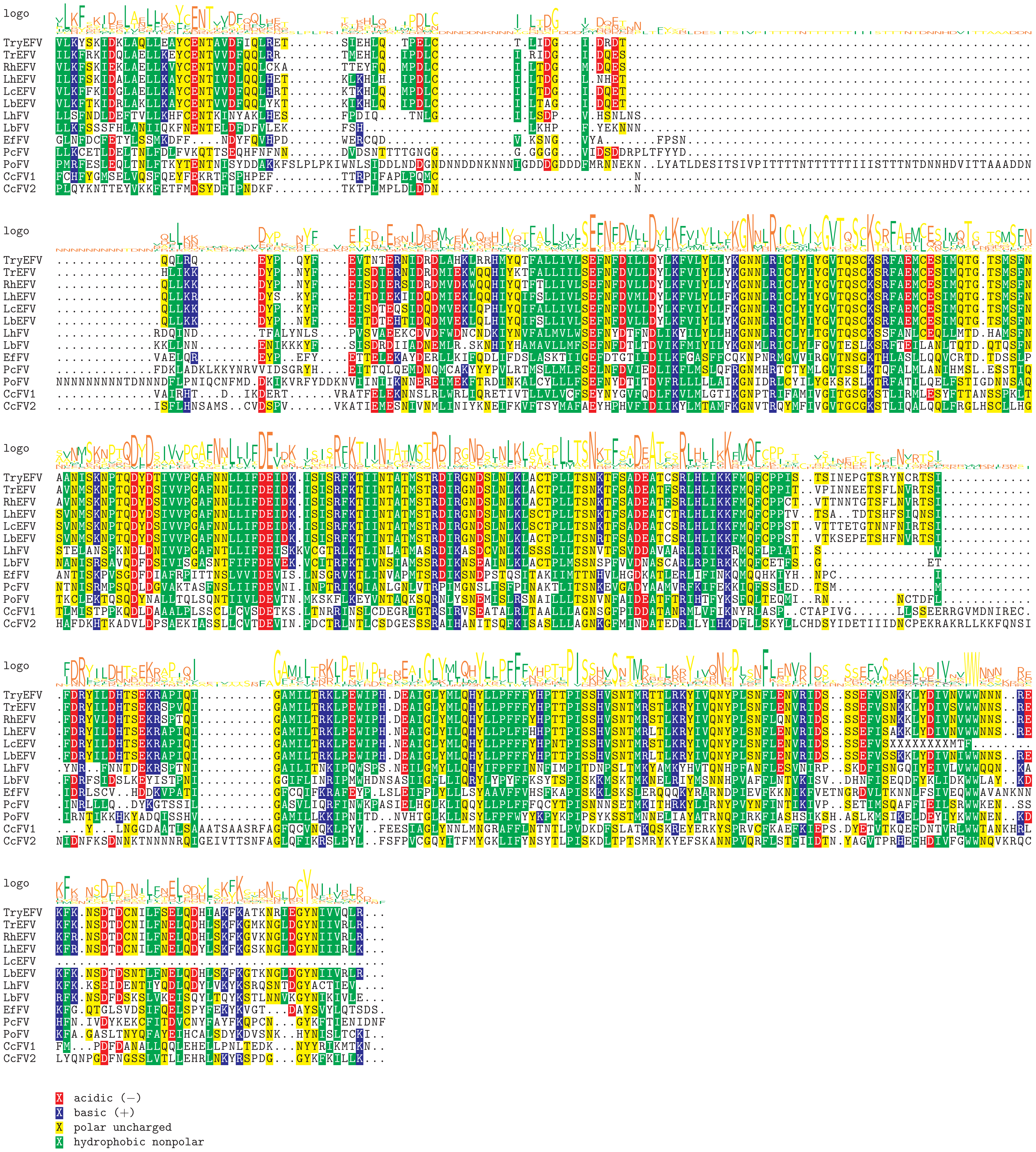

Supplement: FiguresS13 [file 595704_file03.zip › FiguresS13_All_Eucoilini_filamentous_alignments/Cluster2683-Cluster470-Cluster79-Cluster2592_LbFVorf92_[helicase]_MSA.pdf]

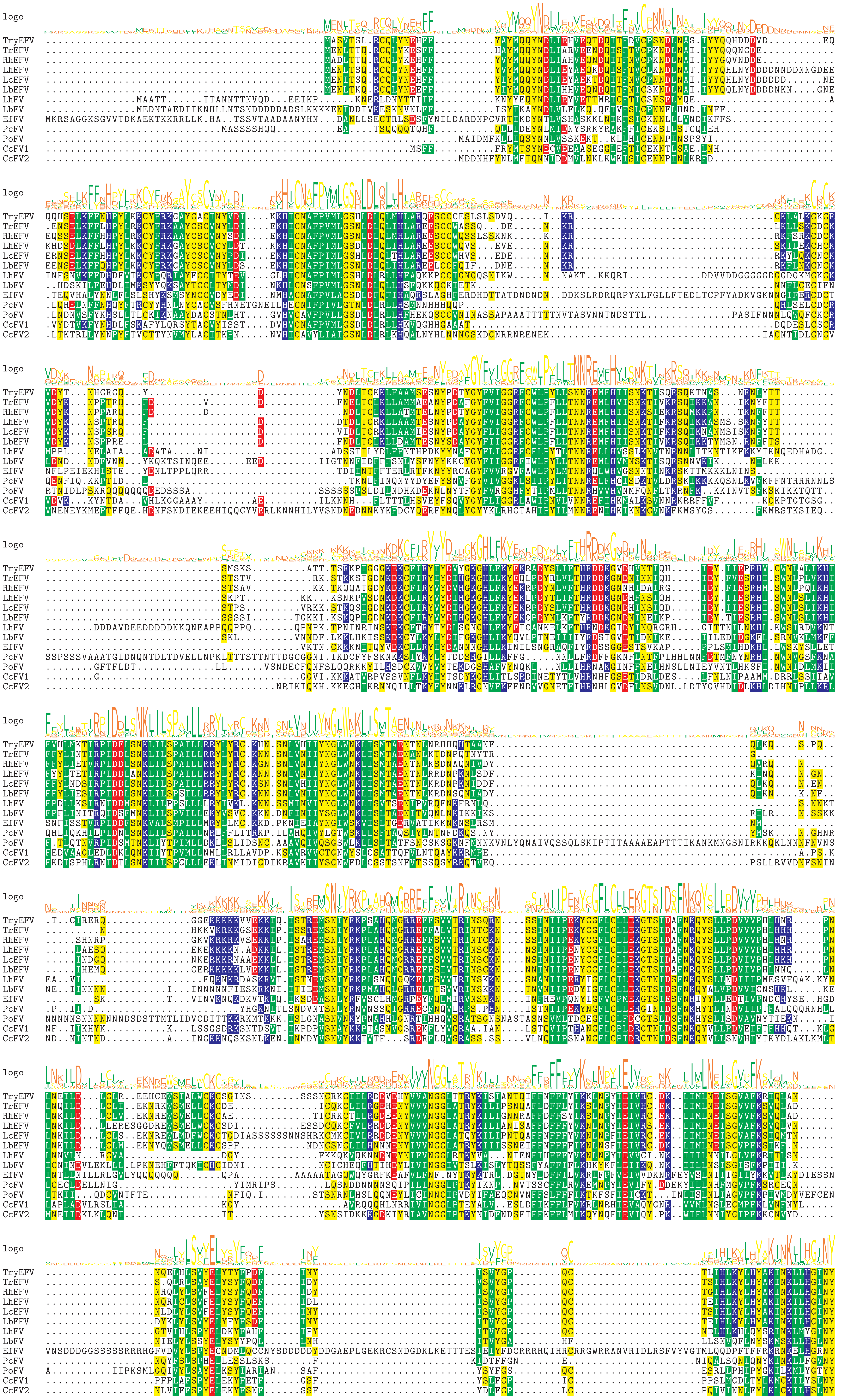

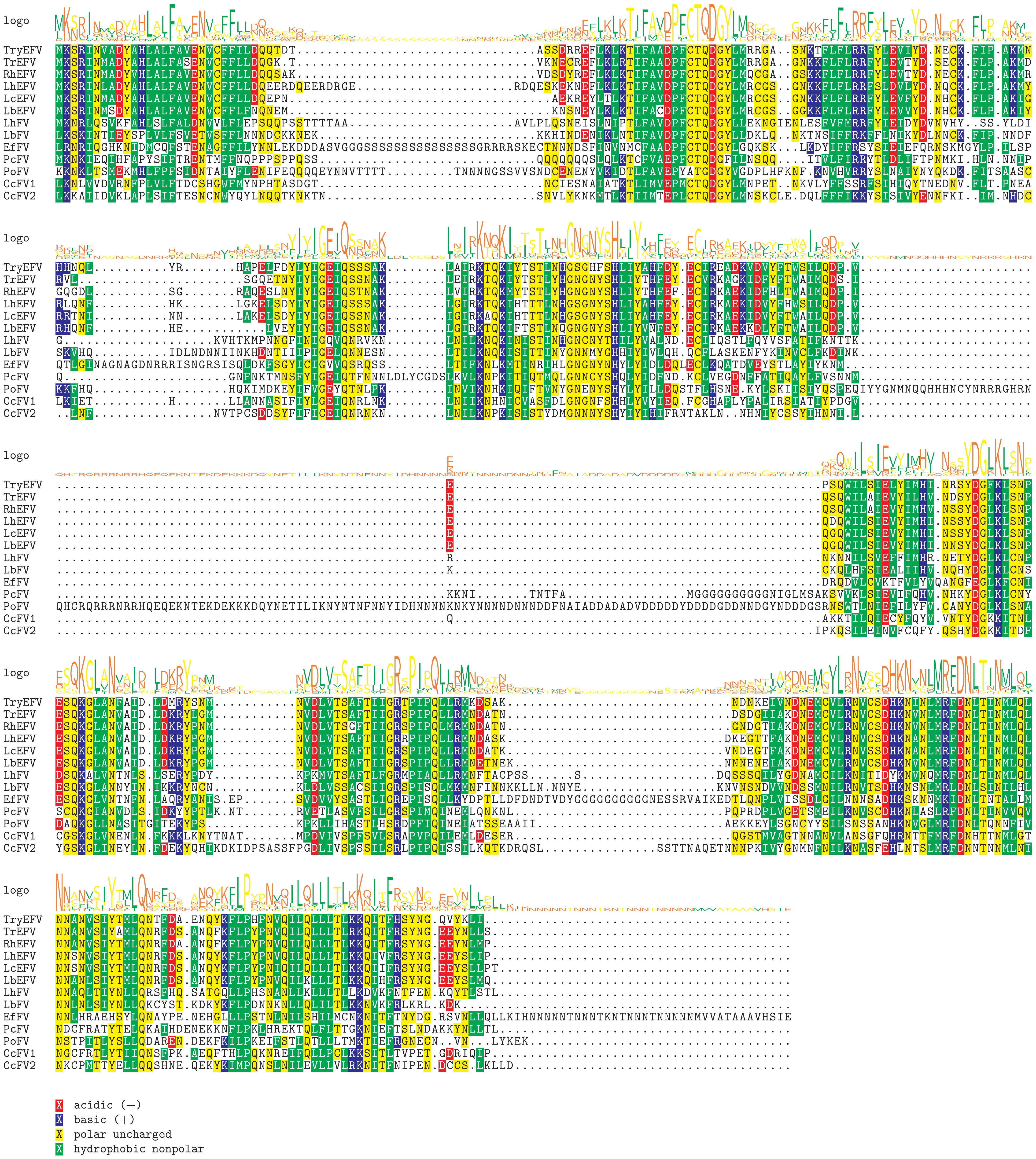

Supplement: FiguresS13 [file 595704_file03.zip › FiguresS13_All_Eucoilini_filamentous_alignments/Cluster1614-Cluster91_LbFVorf96_[lef-8]_MSA.pdf]

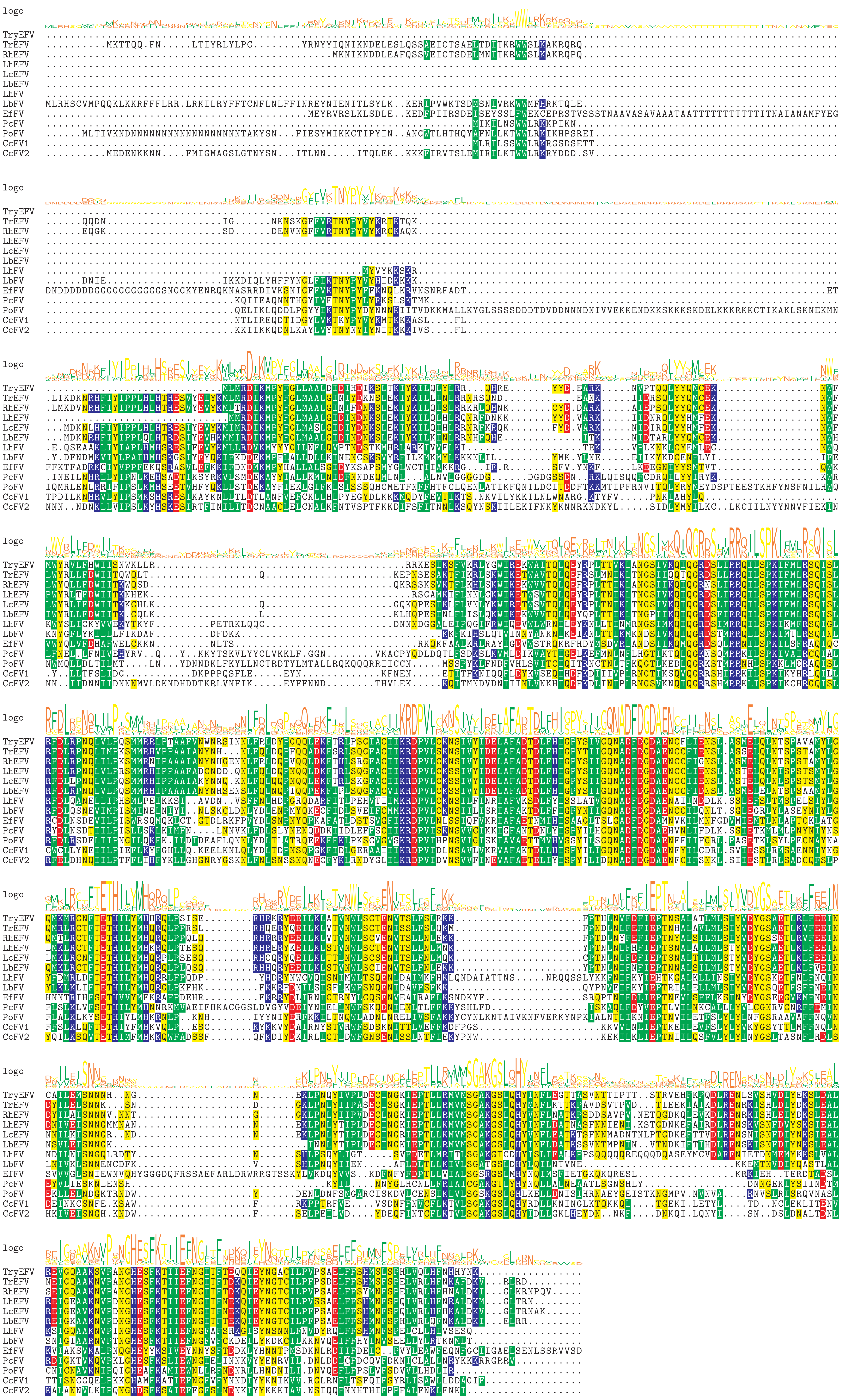

X acidic (−)
X basic (+)
X polar uncharged
X hydrophobic nonpolar

Supplement: FiguresS13 [file 595704_file03.zip › FiguresS13_All_Eucoilini_filamentous_alignments/Cluster15_LbFVorf78_[lef-9]_MSA.pdf]

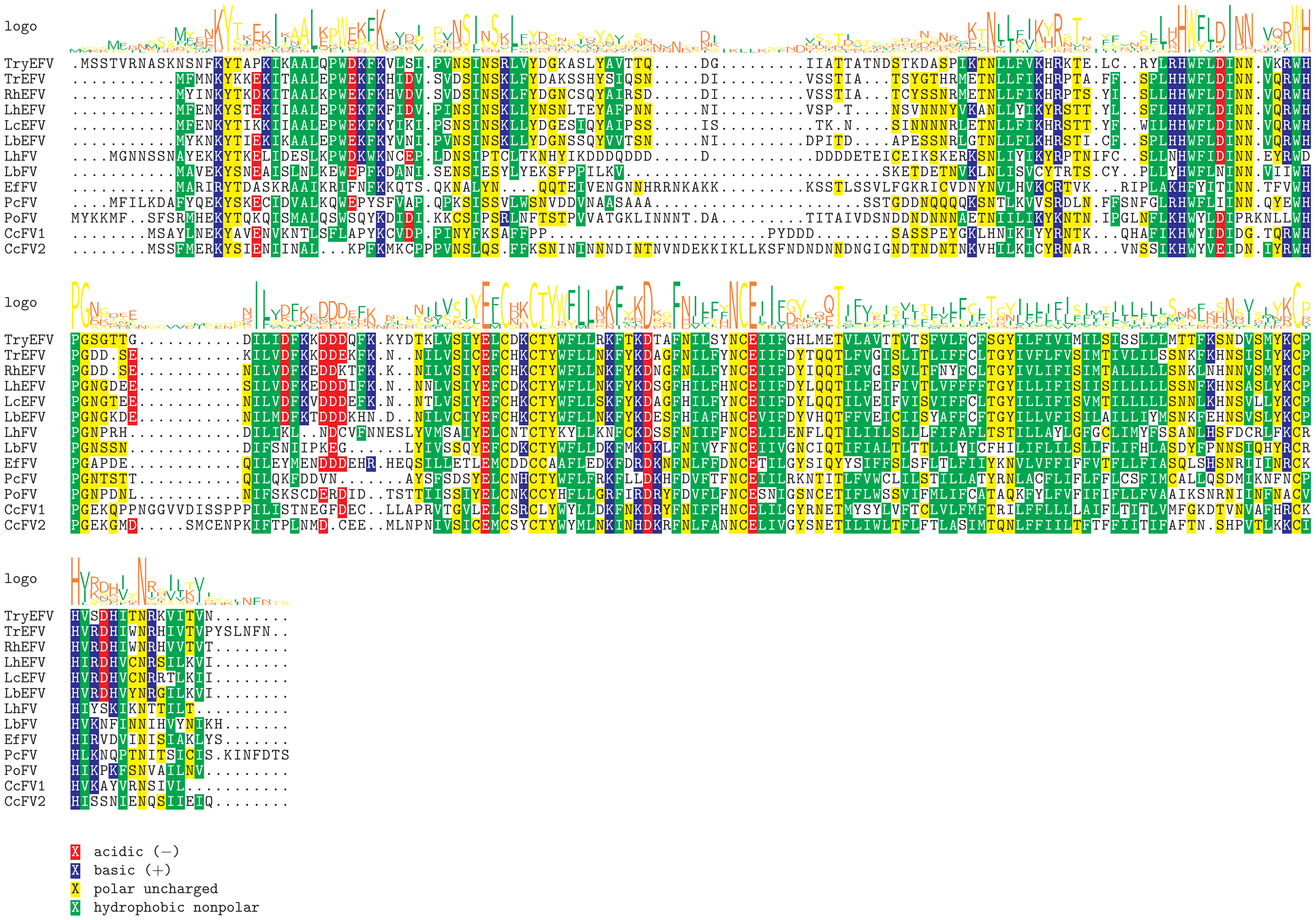

Supplement: FiguresS13 [file 595704_file03.zip › FiguresS13_All_Eucoilini_filamentous_alignments/Cluster965-Cluster60-Cluster519-Cluster44LbFVorf85_[Ac81]_MSA.pdf]
